# Assessment of statistical methods from single cell, bulk RNA-seq and metagenomics applied to microbiome data

**DOI:** 10.1101/2020.01.15.907964

**Authors:** Matteo Calgaro, Chiara Romualdi, Levi Waldron, Davide Risso, Nicola Vitulo

## Abstract

**Background:** The correct identification of differentially abundant microbial taxa between experimental conditions is a methodological and computational challenge. Recent work has produced methods to deal with the high sparsity and compositionality characteristic of microbiome data, but independent benchmarks comparing these to alternatives developed for RNA-seq data analysis are lacking.

**Results:** Here, we compare methods developed for single cell, bulk RNA-seq, and microbiome data, in terms of suitability of distributional assumptions, ability to control false discoveries, concordance, and power. We benchmark these methods using 100 manually curated datasets from 16S and whole metagenome shotgun sequencing.

**Conclusions:** The multivariate and compositional methods developed specifically for microbiome analysis did not outperform univariate methods developed for differential expression analysis of RNA-seq data. We recommend a careful exploratory data analysis prior to application of any inferential model and we present a framework to help scientists make an informed choice of analysis methods in a dataset-specific manner.

## Background

Study of the microbiome, the uncultured collection of microbes present in most environments, is a novel application of high-throughput sequencing that shares certain similarities but important differences from other applications of DNA and RNA sequencing. Common approaches for the microbiome studies are based on the deep sequencing of amplicons of universal marker-genes, such as the 16S rRNA genes, or on whole metagenome shotgun sequencing (WMS). Community taxonomic composition can be estimated from microbiome data by assigning each read to the most plausible microbial lineage using a reference annotated database, with a higher taxonomic resolution in WMS than in 16S [1,2]. The final output of such analyses usually consists of a large, highly sparse taxa per samples count table.

Differential abundance (DA) analysis is one of the primary approaches to identify differences in the microbial community composition between samples and to understand the structures of the microbial communities and the associations between the microbial compositions and the environment. DA analysis has commonly been performed using methods adapted from RNA sequencing (RNA-seq) analysis; however, the peculiar characteristics of microbiome data make differential abundance analysis challenging. Compared to other high-throughput sequencing techniques such as RNA-seq, metagenomic data are sparse, i.e., the taxa count matrix contains many zeros. This sparsity can be explained by both biological and technical reasons: some taxa are very rare and present only in a few samples, while others are very lowly represented and cannot be detected because of an insufficient sequencing depth or other technical reasons.

In recent years, single-cell RNA-seq (scRNA-seq) has revolutionized the field of transcriptomics, providing new insight on the transcriptional program of individual cells, shading light on complex, heterogeneous tissues, and revealing rare cell populations with distinct gene expression profiles [3–6]. However, due to the relatively inefficient mRNA capture rate, scRNA-seq data are characterized by dropout events, which leads to an excess of zero read counts compared to bulk RNA-seq data [7,8]. Thus, with the advent of this technology, new statistical models accounting for dropout events have been proposed. The similarities with respect to sparsity observed in both scRNA-seq and metagenomics data led us to pose the question of whether statistical methods developed for the differential expression of scRNA-seq data perform well on metagenomic DA analysis.

Some benchmarking efforts have compared the performance of methods [9–12] both adapted from bulk RNA-seq and developed for microbiome DA [13,14]. While some tools exist to guide researchers [15], a general consensus on the best approach is still missing, especially regarding the methods’ capability of controlling false discoveries. In this study, we benchmark several statistical models and methods developed for metagenomics [13,14,16–18], bulk RNA-seq [19–21] and, for the first time, single-cell RNA-seq [7,8,22–24] on a collection of manually curated 16S and WMS [25,26] real data as well as on a comprehensive set of simulations. We include in the comparison several tools that take into account the compositional nature of the data: they achieve this through the use of the Dirichlet-Multinomial Distribution (e.g., ALDEx2), Multinomial Distribution with reference frames (Songbird) or the Centered Log Ratio (CLR) transformation (e.g., ALDEx2, mixMC). The novelty of our benchmarking efforts is two-fold. First, we include in the comparison novel methods recently developed in the scRNA-seq and metagenomics literatures; second, unlike previous efforts, our conclusions are based on several performance metrics on real data that range from type I error control and goodness of fit to replicability across datasets, concordance among methods, and enrichment for expected DA microbial taxa.

## Results

We benchmarked a total of 18 approaches (Additional file 1: Supplementary Table 2) on 100 real dataset (Additional file 1: Supplementary Table 1), evaluating goodness of fit, type I error control, concordance, and power, through i) reliability of DA results in real data based on enrichment analysis; ii) specificity and sensitivity using 28,800 simulated datasets (Fig. 1; Additional file 2: Supplementary Table 4).

**Figure 1:**
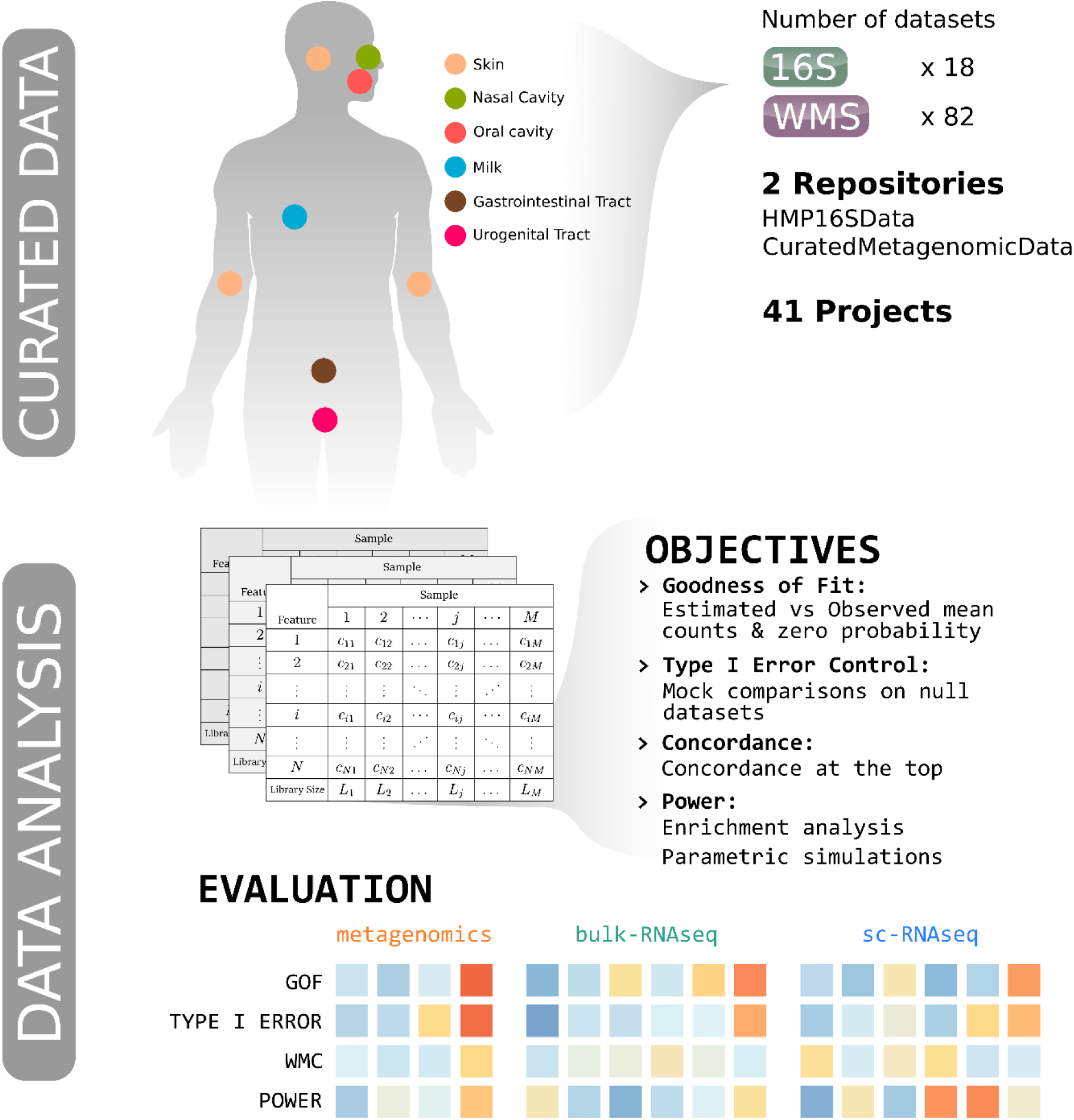
Starting from 41 Projects collected in 2 manually curated data repositories (*HMP16SData* and *curatedMetagenomicData* Bioconductor packages), 18 16S and 82 WMS datasets were downloaded. Biological samples belonged to several body sites (e.g. oral cavity), body subsites (e.g. tongue dorsum) and conditions (e.g. healthy vs disease). Feature per sample count tables were used in order to evaluate several objectives: goodness of fit (GOF) for 5 parametric distributions, type I error control, concordance, and power for 18 differential abundance detection methods. Methods, developed in metagenomics, bulk-RNAseq or sc-RNAseq, were ranked using empirical evaluations of the above cited objectives.

The benchmarked methods include both DA methods specifically proposed in the metagenomic literature and methods proposed in the single-cell and bulk RNA-seq fields. The manually curated real datasets span a variety of body sites and characteristics (e.g., sequencing depth, alpha and beta diversity). The diversity of the data allowed us to test each method on a variety of circumstances, ranging from very sparse, very diverse datasets, to less sparse, less diverse ones.

We first analyzed 18 16S, 82 WMS and 28 scRNA-seq public datasets in order to assess whether scRNA-seq and metagenomic data are comparable in terms of sparsity. We observed overlap in the fractions of zero counts between the scRNA-seq, WMS, and 16S, but with scRNA-seq datasets having a lower distribution of sparsities (ranging from 12% to 75%) as compared to 16S (ranging from 55% to 83%) and WMS datasets (ranging from 35% to 89%) whose distributions of zero frequencies were not significantly different from each other (Wilcoxon test, W = 734, p = 0.377, Fig. S1a-b). To establish whether the difference between scRNA-seq and metagenomic data was due to the different number of features and samples, which are intrinsically related to sparsity, we explored the role of library size and experimental protocol (Fig. S1c). scRNA-seq datasets showed a marked difference in terms of number of features and sparsity degree, as they are derived from different experimental protocols. Full-length data (e.g., Smart-seq) are on average sparser than droplet-based data (e.g., Drop-seq) but both are less sparse than 16S and WMS.

These results indicate that metagenomic data is even more sparse than scRNA-seq, and thus that zero-inflated models designed for scRNA-seq could at least in principle have good performance in a metagenomic context.

### Goodness of fit

As different methods rely on different statistical distributions to perform DA analysis, we started our benchmark by assessing the goodness of fit (GOF) of the statistical models underlying each method on the full set of 16S and WMS data. For each model, we evaluated its ability to correctly estimate the mean counts and the probability of observing a zero (Fig. 2). We evaluated five distributions: (1) the negative binomial (NB) used in edgeR [19] and DeSeq2 [20], (2) the zero-inflated negative binomial (ZINB) used in ZINB-WaVE [23], (3) the truncated Gaussian Hurdle model of MAST [7], (4) the zero-inflated Gaussian (ZIG) mixture model of metagenomeSeq [13], and (5) the Dirichlet-Multinomial (DM) distribution underlying ALDEx2 [14]. The truncated Gaussian Hurdle model was evaluated following two data transformations, the default logarithm of the counts per million (logCPM) and the logarithm of the counts rescaled by the median library size (see Methods). Similarly, the ZIG distribution was evaluated considering the scaling factors rescaled by either one thousand (as implemented in the metagenomeSeq Bioconductor package) and by the median scaling factor (as suggested in the original paper). We assessed the goodness of fit for each of these models using the stool samples from the Human Microbiome Project (HMP) as a representative dataset (Fig. 2a - d); all other datasets gave similar results (Additional file 1: Supplementary Fig. S2). A useful feature of this dataset is that a subset of samples was processed both with 16S and WMS and hence can be used to compare the distributional differences of the two data types. Furthermore, this dataset includes only healthy subjects in a narrow age range, providing a good testing ground for covariate-free models.

**Figure 2:**
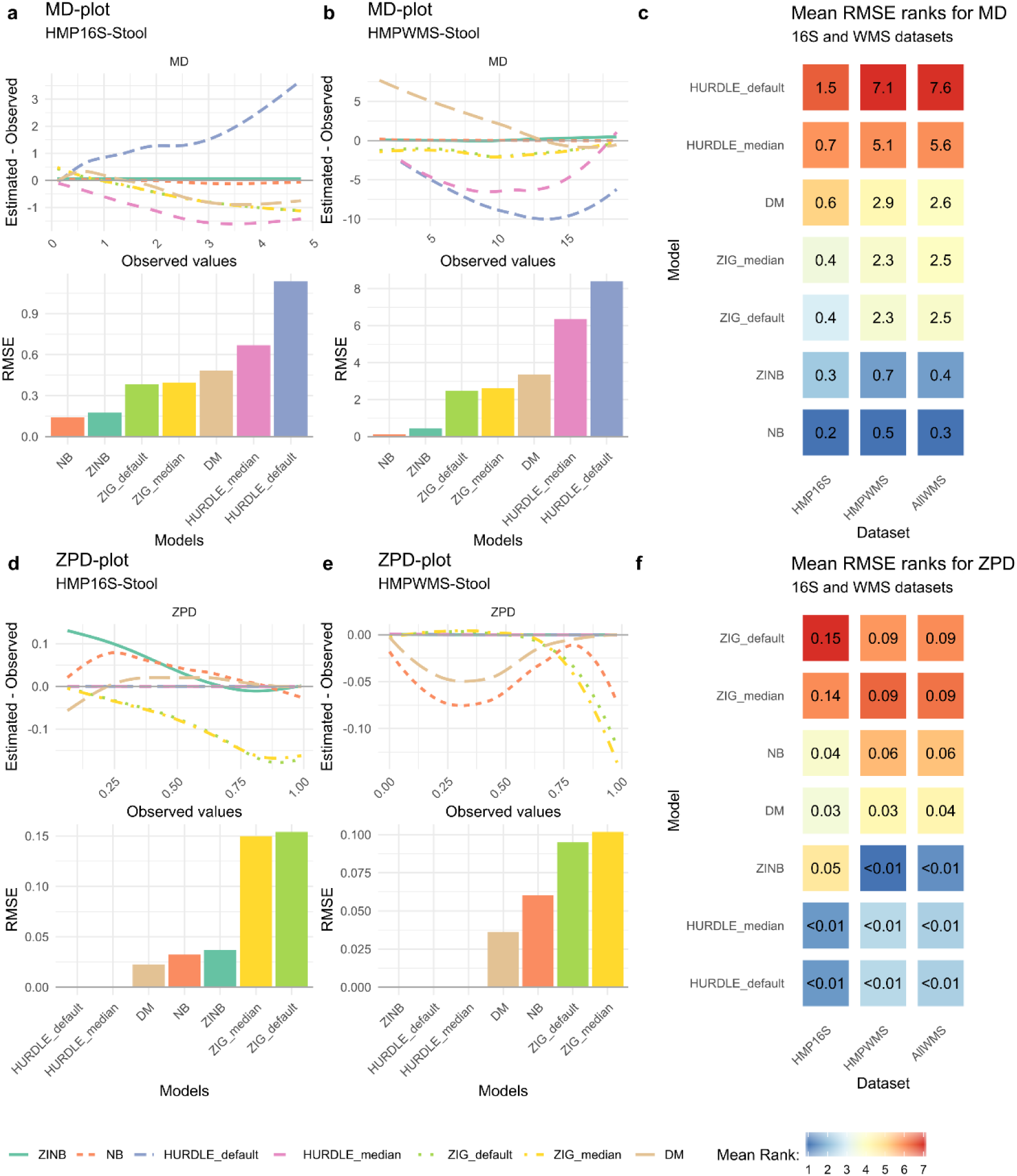
**a.** Mean-Differences (MD) plot and Root Mean Squared Errors (RMSE) for HMP 16S Stool samples. **b.** MD plot and RMSE for HMP WMS Stool samples. **c**. Average rank heatmap for MD performances in HMP 16S datasets, HMP WMS datasets and all other WMS project datasets. The value inside each tile refers to the average RMSE value on which ranks are computed. **d.** Zero Probability-Differences (ZPD; see Methods) plot and RMSE for HMP 16S Stool samples. **e.** ZPD plot and RMSE for HMP WMS Stool samples. **f**. Average rank heatmap for ZPD performances in HMP 16S datasets, HMP WMS datasets and all other WMS project datasets. The value inside each tile refers to the average RMSE value on which ranks are computed.

The NB distribution showed the lowest root mean square error (RMSE, see Methods) for the mean count estimation (MD), followed by the ZINB distribution (Fig. 2a-b). This was true both for 16S and for WMS data, in most of the considered datasets (Additional file 1: Supplementary Fig. S2). Moreover, for both distributions, the difference between the estimated and observed means were symmetrically distributed around zero, indicating that the models did not systematically under- or over-estimate the mean abundances (Fig. 2a-b; Additional file 1: Supplementary Fig. S2). Conversely, the ZIG distribution consistently underestimated the observed means, both for 16S and WMS and independently on the scaling factors (Fig. 2a-b). The Hurdle model was sensitive to the choice of the transformation: rescaling by the median library size rather than by one million reduced the RMSE in both 16S and WMS data (Fig. 2a-b). This was particularly evident in 16S data (Fig. 2a), in which the default logCPM values resulted in a substantial overestimation of the mean count, while the median library size scaling led to under-estimation. Given the clear problems with logCPM, we only used the median library size for MAST and the median scaling factor for metagenomeSeq in all subsequent analyses. The DM distribution overestimated observed means for low-mean count features and underestimated observed values for high-mean count features. This overestimation effect was more evident in WMS than in 16S.

Concerning the ability of models to estimate the probability of observing a zero (referred to as zero probability difference, ZPD), we found that Hurdle models provided good estimates of the observed zero proportion for 16S (Fig. 2c) and WMS datasets (Fig. 2d). The NB and ZINB distributions, on the other hand, tended to overestimate the zero probability for features with a low observed proportion of zero counts in 16S (Fig. 2c). In WMS data, the ZINB distribution perfectly fitted the observed proportion of zeros, while the NB and DM models tended to underestimate it (Fig. 2d). Finally, the ZIG distribution always underestimated the observed proportion of zeros, especially for highly sparse features (Fig. 2c-d).

In summary, across all datasets, the best fitting distributions were the NB and ZINB: the NB distribution seemed to be particularly well-suited for 16S datasets, while the ZINB distribution seemed to better fit WMS data (Fig. 2e). We hypothesize that this is due to the different sequencing depths of the two platforms. In fact, while our 16S datasets have an average of 4891 reads per sample, in WMS the mean depth is 3.6×10^8^ (3×10^8^ for HMP). To confirm this observation, we carried out a simulation experiment by down-sampling reads from deep-sequenced WMS samples (rarefaction): while the need for zero inflation seemed to diminish as we got closer to the number of reads typical of the corresponding 16S experiments, the profile did not completely match between approaches (Additional file 1: Supplementary Fig. S4b). This suggests that, while sequencing depth is an important contributing factor, it is not enough to completely explain the distributional differences between the two platforms.

### Type I error control

We next sought to evaluate type I error rate control of each method, i.e., the probability of the statistical test to call a feature DA when it is not. To do so, we considered mock comparisons between the same biological Stool HMP samples (using the same Random Sample Identifier in both 16S and WMS), in which no true DA is present. Briefly, we randomly assigned each sample to two experimental groups and compared them, repeating the process 1000 times (see Methods for additional details). In this setting, the p-values of a perfect test should be uniformly distributed between 0 and 1 (ref. [27]) and the false positive rate (FPR or observed *α*), which is the observed proportion of significant tests, should match the nominal value (e.g., *α* = 0.05).

To evaluate the impact of both the normalization step and the estimation and testing step in bulk RNA-seq inspired methods, we included in the comparison both edgeR with its default normalization (TMM), as well as with DESeq2 recommended normalization (“poscounts”, i.e., the geometric mean of the positive counts) and vice versa (Table S2). Similarly, because the zinbwave observational weights can be used to apply several bulk RNA-seq methods to single-cell data [24], we have included in the comparison edgeR, DESeq2, and limma-voom with zinbwave weights.

The qq-plots and Kolmogorov-Smirnov (KS) statistics in Figure 3 show that most methods achieved a p-value distribution reasonably close to the expected uniform. The notable exceptions in the 16S experiment were edgeR with TMM normalization and robust dispersion estimation (edgeR_TMM_robustDisp), metagenomeSeq, and ALDEx2 (Fig. 3a-b). While the former two appeared to employ liberal tests, the latter was conservative in the range of p-values that are typically of interest (0 - 0.1). In the WMS data, departure from uniformity was observed for metagenomeSeq and edgeR_TMM_robustDisp, and limma_voom_TMM_zinbwave, which employed liberal tests, as well as corncob_LRT, ALDEx2, and scde, which were conservative in the range of interest (Fig. 3c-d). We note that in the context of DA, liberal tests will lead to many false discoveries, while conservative tests will control the type I error at a cost of reduced power, potentially hindering true discoveries.

**Figure 3:**
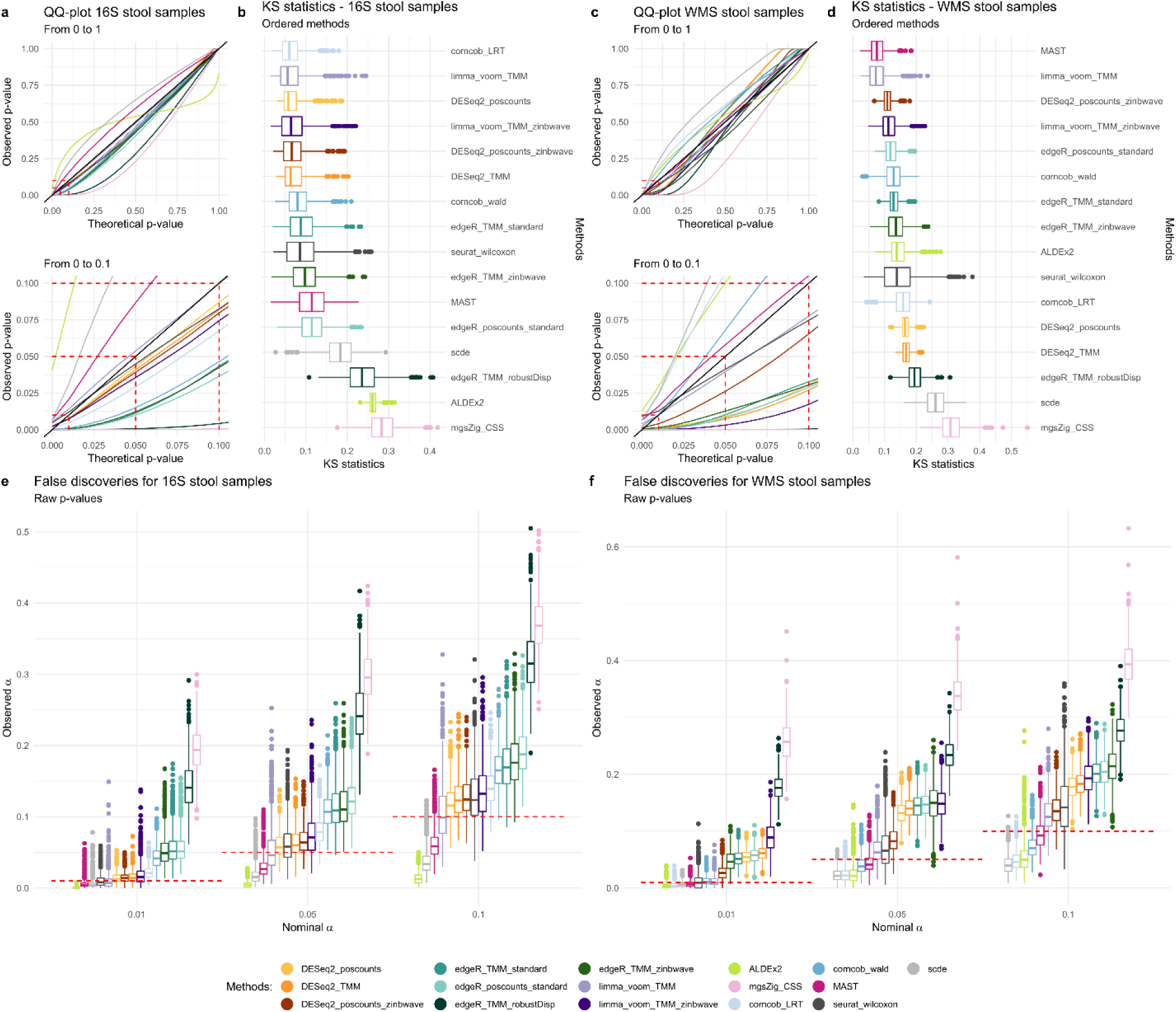
**a.** Quantile-quantile plot from 0 to 1 and 0 to 0.1 zoom for DA methods in 41 16S HMP stool samples. Average curves for mock comparisons reported. **b.** Kolmogorov-Smirnov statistic boxplots for DA methods in 41 16S HMP stool samples. **c.** Quantile-quantile plot from 0 to 1 and 0 to 0.1 zoom for DA methods in 41 WMS HMP stool samples. Average curves for mock comparisons reported. **d.** Kolmogorov-Smirnov statistic boxplots for DA methods in 41 WMS HMP stool samples. **e.** Boxplots for the proportion of raw p-values lower than 0.01, 0.05, 0.1 values of the commonly used thresholds of nominal *α* for 41 16S stool samples. **f.** Boxplots for the proportion of raw p-values lower than 0.01, 0.05, 0.1 values of the commonly used thresholds of nominal *α* for 41 WMS stool samples.

We next recorded the FPR by each method (by definition all discoveries are false positives in this experiment) and compared it to its expected nominal value. This analysis confirmed the tendencies observed in Figures 3a-b and 3c-d. In particular, edgeR_TMM_robustDisp and metagenomeSeq were very liberal in both 16S (Fig. 3e) and WMS data (Fig. 3f); in the case of metagenomeSeq, as much as 30% of the features were deemed DA in the 16S datasets when claiming a nominal FPR of 5% (Fig. 3e). ALDEx2, scde and MAST, albeit conservative, were able to control type I error. In between these two extremes, edgeR, DESeq2 and limma showed an observed FPR slightly higher than its nominal value. In particular, DESeq2-based methods, limma-voom, and MAST were very close to the nominal FPR for 16S (Fig. 3e), while limma-voom, MAST, and corncob (with Wald test) were the closest in WMS data (Fig. 3f). Of note, corncob seemed slightly conservative in WMS data and slightly liberal in 16S data, with LRT being closer than Wald to the nominal value in 16S (Fig. 3e) and vice versa in WMS data (Fig. 3f). The zinbwave weights showed mixed results: DESeq2 with zinbwave weights was better than the unweighted versions in WMS, while the weights did not help edgeR and limma in controlling the type I error rate. Taken together, these results suggest that the majority of the methods does not control the type I error rate, both in 16S and WMS data, confirming previous findings [10,12]. However, for most approaches, the observed FPR is only slightly higher than its nominal value, making the practical impact of this result unclear.

### Between-method concordance

To measure the ability of each method to produce replicable results in independent data, we looked at six datasets [25,26,28–30] (Additional file 1: Supplementary Table S3), with different alpha and beta diversity, as well as different amounts of DA between two experimental conditions (Additional file 1: Supplementary Figure S5). Each dataset was randomly split in two equally sized subsets and each method was separately applied to each subset. The process was repeated 100 times (see Methods for details). To assess the ability of methods to return concordant results from independent samples, we employed the Concordance At the Top [31](CAT) measure to assess between-method concordance (BMC) by comparing the list of DA features across methods in the subset (ranked by p-value when available or by importance in the case of the songbird and mixMC; see Methods). We used BMC to (i) group methods based on their degree of agreement, and (ii) identify those methods sharing the largest amount of discoveries with the majority of the other methods. Although concordance is not a guarantee of validity, it is a requirement of validity, so methods sharing the largest amount of discoveries with the majority of other methods may be more likely to also be producing valid results.

Concordance analysis performed on 16S Tongue Dorsum vs Stool dataset (Fig. 4a) showed that the methods clustered within two distinct groups: the first comprising all methods that include a TMM normalization step, songbird, and scde, the second containing all the other approaches (Fig. 4a). Even within the second group, methods segregated by normalization, as can be seen by the tight clustering of all the methods that include a poscount normalization step (Fig. 4a). This indicates that, in 16S data, the choice of the normalization has a pronounced effect on inferential results, even more so than the choice of the statistical test. A similar result was previously observed in bulk RNA-seq data [32]. The use of observational weights to account for zero inflation did not seem to matter in these data, and in general, scRNA-seq methods did not agree with each other (Fig. 4a). Similarly, the clustering did not separate compositional and non-compositional methods (Fig. 4a). We noted that metagenomeSeq was not concordant with any other method and that the two corncob approaches formed a tight group, confirming that modeling strategies have more impact than the choice of the test statistics in these data.

**Figure 4:**
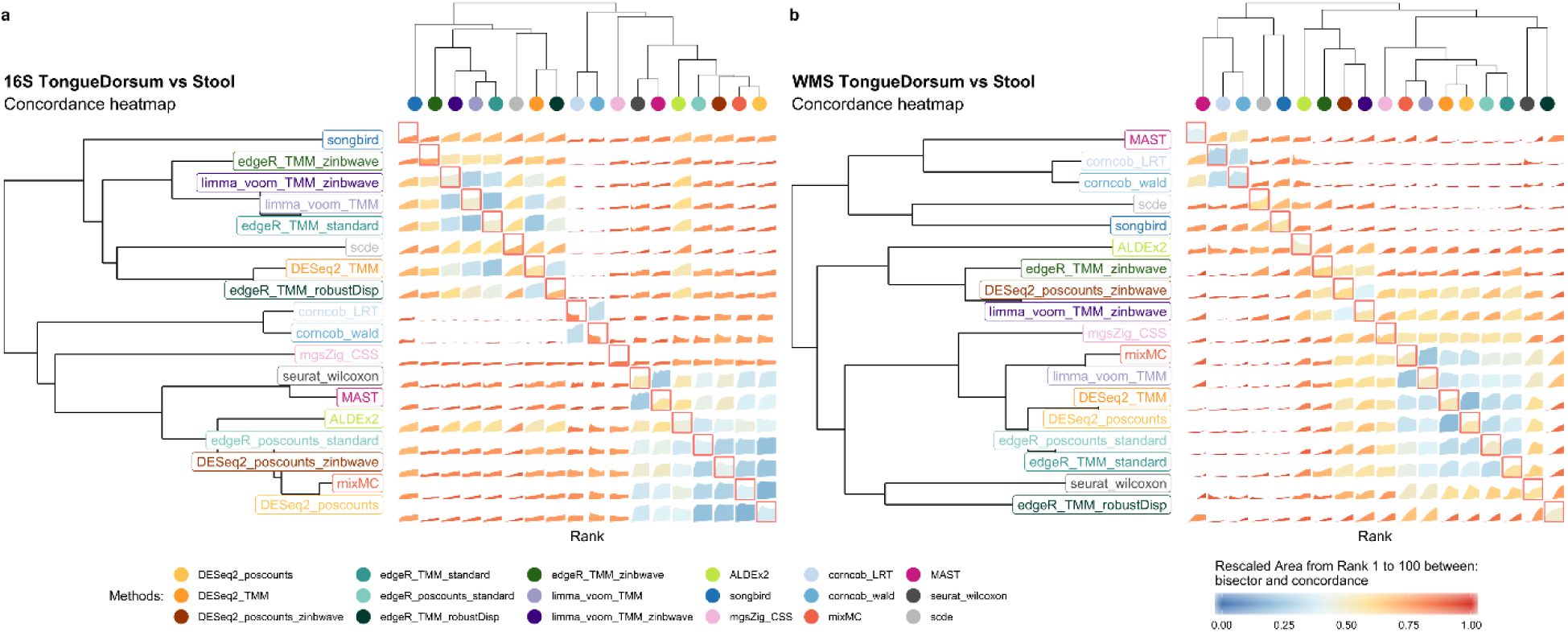
**a.** BMC and WMC (main diagonal) averaged values from rank 1 to 100 for DA methods evaluated in replicated 16S Tongue Dorsum vs Stool comparisons. **b.** BMC and WMC (main diagonal) averaged values from rank 1 to 100 for DA methods evaluated in replicated WMS Tongue Dorsum vs Stool comparisons.

A different picture emerged from the analysis of the WMS data (Fig. 4b). Here, methods clustered by the testing approach. The bottom cluster comprised the bulk RNA-seq methods with the inclusion of the Wilcoxon nonparametric approach, metagenomeSeq and mixMC. The middle cluster consisted of the zinbwave methods and ALDEx2. The top cluster comprised MAST, corncob, scde, and songbird. Overall, mixMC and the methods based on NB generalized linear models showed the highest BMC values. When observational weights were added to those models, the BMC decreased, but still a good level of concordance was observed with their respective unweighted version.

We noted that the BMC is highly dataset-specific and depends on the amount of DA between the compared groups. Indeed, BMC decreased with the beta diversity of the dataset, and the role of normalization became less clear (Additional file 1: Supplementary Fig. S6).

### Within-method concordance

The CAT metric was used again for assessing the within-method concordance (WMC), i.e., the amount of concordance of the results of each method on the two random subsets.

WMC was clearly dataset-dependent, showing high levels of concordance in datasets with a high differential signal (e.g., tongue vs. stool, Fig. 5a) and low concordance in datasets with a low differential signal (e.g., supragingival vs. subgingival, Fig. 5e). Overall, the reproducibility of the results in WMS studies was slightly higher than that of 16S datasets. In terms of method comparison, corncob showed high levels of concordance in WMS datasets but lower concordance in all 16S datasets (Fig. 5). Similarly, songbird showed the highest concordance in mid (Fig. 5d) and low (Fig. 5f) diversity WMS datasets but did not perform well in 16S (especially for the highly diverse TongueDorsum vs. Stool comparison; Fig 5a). The addition of zinbwave weights to edgeR, DESeq2 and limma-voom did not always help: it was sometimes detrimental, e.g., for edgeR in the schizophrenia dataset (Fig. 5d), and sometimes led to an improvement in replicability, e.g., for limma-voom in the Tongue Dorsum vs. Stool dataset (Fig. 5a). The schizophrenia dataset had the lowest numerosity among all the datasets evaluated, suggesting that sample size may play an important role in estimating zinbwave weights. While this analysis confirmed the unsatisfactory performance of metagenomeSeq (Fig 5a,b,f), ALDEx2, which was very conservative in terms of type I error control (Fig. 3), showed overall good performance, with the notable exception of the high-diversity WMS dataset (Fig. 5b), for which it was the worst performing method. To sum up, the highest concordance was measured, in all WMS datasets, by the corncob-based and songbird methods, while RNA-seq methods performed better in 16S datasets, confirming that the two platforms yield substantially different data. mixMC was the only method that never showed poor concordance regardless of the technology and of the diversity of the compared groups.

**Figure 5:**
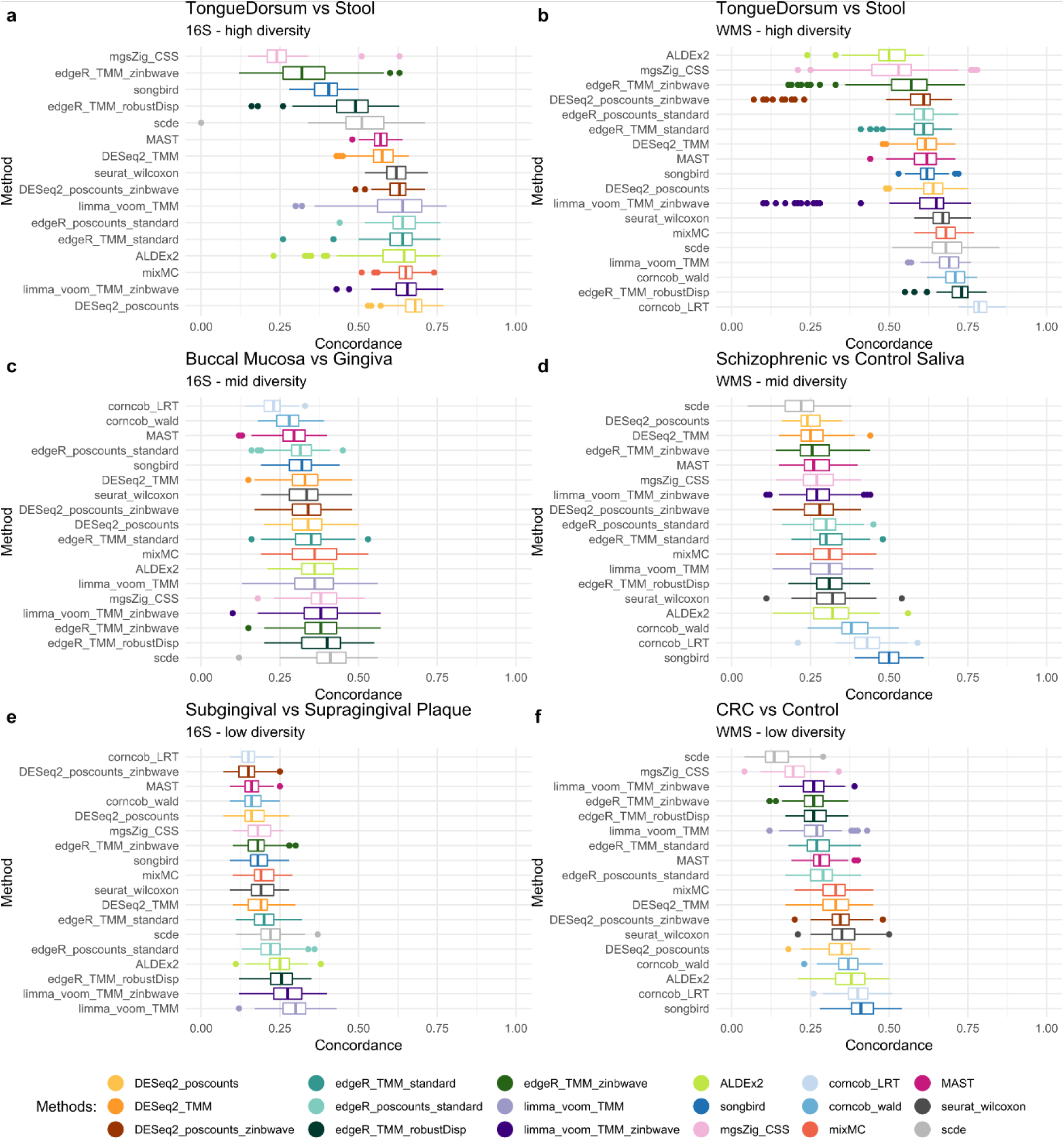
**a.** Boxplot of WMC on high level diversity, Tongue Dorsum vs Stool, 16S datasets. Due to the high sparsity and low sample size of the dataset, the CAT at rank 100 was not computable for corncob methods: only for a few features it was possible to estimate the model. **b.** Boxplot of WMC on high diversity, Tongue Dorsum vs Stool, WMS datasets. **c.** Boxplot of WMC on mid level diversity, Buccal Mucosa vs Attached Keratinized Gingiva, 16S datasets. **d.** Boxplot of WMC on mid level diversity, Schizophrenic vs Healthy Control saliva samples, WMS datasets. **e.** Boxplot of WMC on low level diversity, Supragingival vs Subgingival plaque, 16S datasets. **f.** Boxplot of WMC on low level diversity, Colon Rectal Cancer patient vs Healthy Control stool samples, WMS datasets.

Taken together, these analyses suggest that both BMC and WMC are highly dependent on the amount of DA observed in the dataset: higher DA leads to a higher concordance. Moreover, WMC was similar among the compared methods, indicating that the reproducibility of the DA results depends more on the strength of DA than on the choice of the method (Figure 5).

### Enrichment Analysis

While mock comparisons and random splits allowed us to evaluate model fit and concordance, these analyses did not assess the correctness of the discoveries. In fact, even the method with the highest WMC could nonetheless consistently identify false positive DA taxa.

While the lack of ground truth makes it challenging to assess the validity of DA results in real data, enrichment analysis [33] can provide an alternative solution to rank methods in terms of their ability to identify as significant the taxa that are known to be differentially abundant between two groups.

Here, we leveraged the peculiar environment of the gingival site: the supragingival biofilm is directly exposed to the open atmosphere of the oral cavity favoring the growth of aerobic or facultative anaerobic species. In the subgingival biofilm however, the atmospheric conditions gradually become strict anaerobic, favoring the growth of the associated species [34]. From the comparison of the two sites, we thus expected to find an abundance of aerobic microbes in the supragingival plaque and of anaerobic bacteria in the subgingival plaque. DA analysis should reflect this difference by finding an enrichment of aerobic (anaerobic) bacteria among the DA taxa with a positive (negative) log-fold-change.

We tested this hypothesis by comparing 38 16S supragingival and subgingival samples (for a total of 76 samples) from the HMP (see Methods for details). The DA methods showed a wide range of power, identifying 2 (ALDEx2) through 305 (metagenomeSeq) significantly DA taxa (Fig. 6a). However, almost all methods correctly found an enrichment of anaerobic microbes among the taxa under-abundant in supragingival and an enrichment of aerobic microbes among the over-abundant ones (Fig. 6a; Additional file 1: Supplementary Figure S7). Furthermore, as expected, no enrichment was found for facultative anaerobic microbes, which are able to switch between aerobic and anaerobic respiration (Fig. 6a).

**Figure 6:**
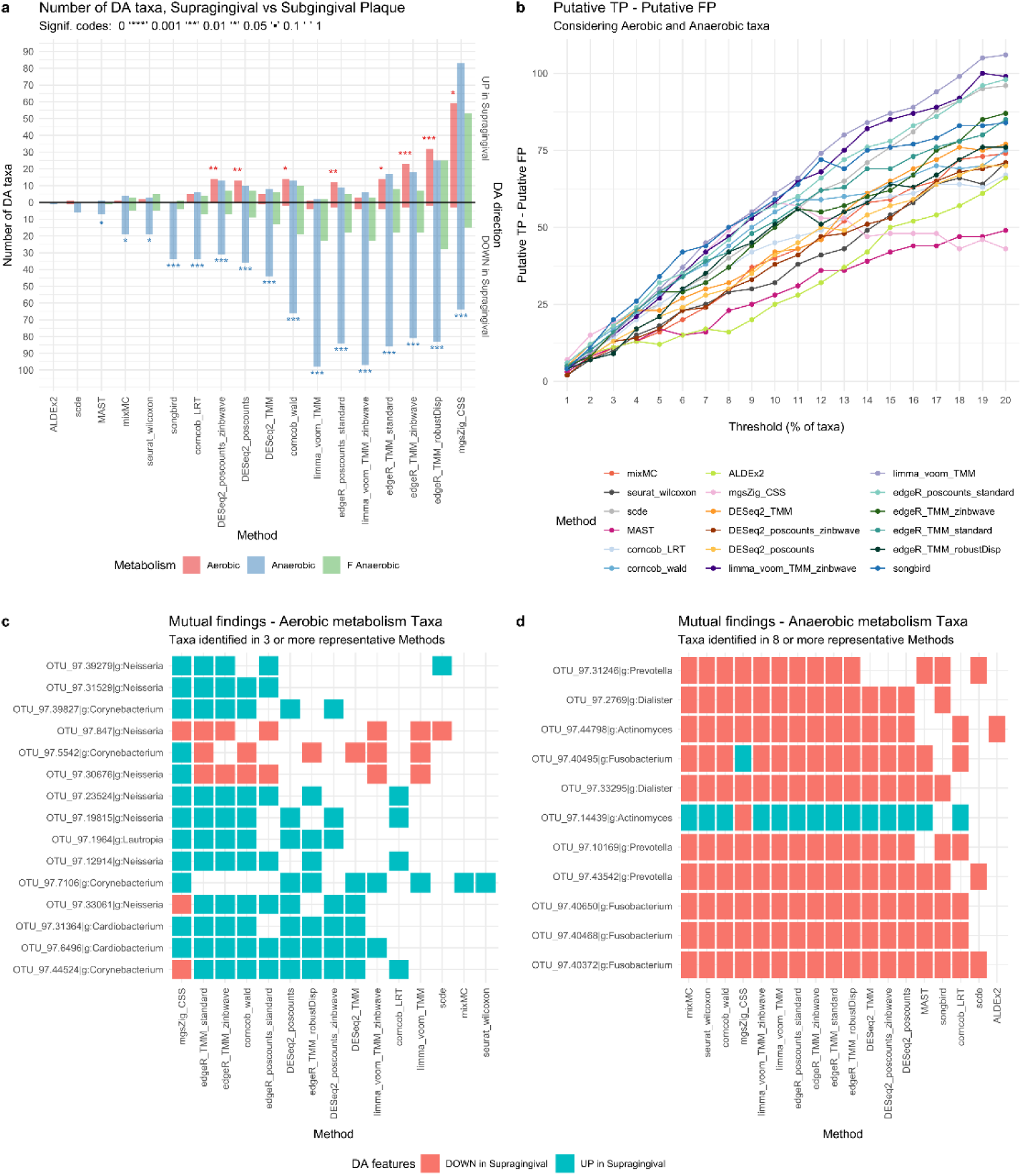
38vs38 Supragingival vs Subgingival Plaque 16S samples **a.** Barplot for the enrichment tests performed on the DA taxa found by each method using an adjusted p-value of 0.1 as threshold for significance (top 10% ranked taxa for *songbird*). Each bar represents the number of findings, UP in Supragingival or DOWN in Supragingival Plaque compared to Subgingival Plaque, regarding Aerobic, Anaerobic and Facultative Anaerobic taxa metabolism. A Fisher exact test is performed to establish the enrichment significance which is represented with signif. codes. **b.** Difference between putative True Positives (TP) and putative False Positives (FP) (y-axis) for several significance thresholds (x-axis). Each threshold represents the top percent ranked taxa, using the ordered raw p-value lists as reference (loading values for *mixMC* and differentials for *songbird*). **c.** Aerobic metabolism taxa mutually found by 3 or more methods from the subset of the representative methods. **d.** Anaerobic metabolism taxa mutually found by 8 or more methods from the subset of the representative methods.

Although most methods performed well, scde, ALDEx2, and MAST had too low power to detect any enrichment (at 0.05 significance level), as their number of identified DA taxa was very low (Fig. 6a). This analysis confirmed the conservative behavior of these methods in 16S data (Fig. 3e). Finally, metagenomeSeq and edgeR with robust dispersion estimation, found the correct enrichments, but they also identified many anaerobic taxa with a positive log-fold-change (Fig. 6a), confirming their liberal tendencies (Fig. 3e). Overall, these results were confirmed by the same comparison in WMS data (Additional file 1: Supplementary Figure S8), but the reduced sample size of our WMS dataset resulted in a reduced power to detect DA for all methods (see Methods).

To further explore the ability of each method to correctly rank the DA taxa independently of its power, we tested whether over-abundant aerobic taxa and under-abundant anaerobic taxa were more likely to be ranked at the top when ranking taxa by each method’s test statistics. To do so, we considered the top K taxa (with K from 1% to 20%; see Methods) and computed the difference between putative true positives (TP; over-abundant aerobic taxa and under-abundant anaerobic taxa) and putative false positives (FP; under-abundant aerobic taxa and over-abundant anaerobic taxa; Fig. 6b). Reassuringly, increasing the threshold resulted in a larger difference between TP and FP for most methods (Fig. 6b), indicating that independently of their power, the large majority of methods are able to highly rank true positive taxa. This becomes particularly important for the methods with a low power, suggesting that in these cases a more liberal p-value threshold may be applied. However, metagenomeSeq’s performance deteriorates after the 10% threshold, suggesting that this method starts to identify more false positives (Fig. 6b): this is particularly problematic since its adjusted p-value threshold identifies 34% of DA taxa. Among the other methods, MAST and ALDEx2 showed a consistently lower performance, while limma-voom was the best performer at permissive thresholds, and songbird was the best performer at strict thresholds (Fig. 6b).

The majority of aerobic taxa were found DA by just a handful of methods, with only 15 aerobic taxa, out of 75 unique taxa, identified as DA by 3 or more representative methods (see Methods; Fig. 6c). All of them belonged to the genera *Cardiobacterium, Neisseria, Lautropia, Corynebacterium*, found to be among the most prevalent genera in supragingival plaques in an independent study [35]. On the other hand, 57 anaerobic taxa, out of 161 unique taxa, were found DA by 5 or more representative methods (see Methods; Fig. 6d; Additional file 1: Supplementary Fig. S9). Among these, *Fusobacterium, Prevotella, Porphyromonas, Treponema* are known to be abundant in subgingival plaque [36,37]. Despite the small sample size for WMS data (n=10) of, enrichment and DA analysis were largely consistent, including several strains of *Neisseria* and several species of *Treponema* found to be DA (Additional file 1: Supplementary Fig. S8c,d). Overall, similar methods tended to identify a higher number of mutual taxa, confirming our previous findings in the concordance analysis (Additional file 1: Supplementary Fig. S6) and highlighting how different statistical test and normalization approaches have a big impact on the identified DA.

### Parametric simulations

To further validate the performances of the methods we turned to simulated data. Given the results of our GOF analysis (Fig. 2), we only used the NB and ZINB distributions to simulate 7200 and 19200 scenarios, respectively, mimicking both 16S and WMS data. The simulated data differ in sample size, proportion of DA features, effect size, proportion of zeros, and whether there was an interaction between the amount of zeros and DA (sparsity effect, see Methods for details).

In general, we found that the results confirmed our expectations (Additional file 2: Supplementary Fig. S11). The parametric distribution that generated the data had great influence on the method performances and the methods that rely on NB and ZINB generally performed better compared to the other methods. As an example, MAST, which showed overall good results in real data, did not behave in simulations, partly because of the misspecified model with respect to the data generating distribution.

As expected, all methods’ performances increased as the sample size and/or the effect size increased. Confirming our real data results, we finally observed that metagenomeSeq, scde, and edgeR-robust performed poorly. Details on the simulated data analysis can be found in Additional file 2.

## Discussion

We have investigated different theoretical and practical issues related to the analysis of metagenomic data. The main objective of the study was to compare several DA detection methods adapted from bulk RNA-seq, single-cell RNA-seq, or specifically designed for metagenomics. Unsurprisingly, there is no single method that outperforms all others in all the tested scenarios. As is often the case in high-throughput biology, the results are data-dependent and careful data exploration is needed to make an informed decision on which workflow to apply to a specific dataset. We recommend applying our explorative analysis framework to gain useful insights about the assumptions of each method and their suitability given the data at hand. To this end, we provide all the R scripts to easily reproduce the analyses of this paper on any given dataset (see code availability).

Our GOF analysis highlighted the advantages of using count models for the analysis of metagenomics data. The goodness of fit of zero inflated models seemed dependent on whether the data come from 16S or WMS experiments. The difference between these two approaches translates to different count data structures: while for WMS many features are characterized by a clearly visible bimodal distribution (with a point mass at zero and another mass, quite far from zero, at the second positive mode), 16S data are as sparse as or even more sparse than WMS data, presenting for many features a less clearly bimodal distribution (Additional file 1: Supplementary Fig. S4a). This difference is probably due to a mix of factors: primarily sequencing depth, but also different taxonomic classification between technologies (entire metagenomic sequences versus clusters of similar amplicon sequences), bioinformatics methods for data preprocessing, etc. However, comparing the distribution of several genera on the same samples assayed with 16S and WMS, we observed that many of the zero counts were consistent across platforms and very different read depths, suggesting that many observed zeros are biological and not technical in nature (Additional file 1: Supplementary Fig. S4a). Further analyses are needed to inspect this unsolved issue and related efforts are ongoing in the single-cell RNA-seq literature, where similar differences are observed between protocols with and without unique molecular identifiers [38,39].

Metagenomic data are inherently compositional, but whether incorporating compositionality into the statistical model provides benefits greater than tradeoffs they may introduce is a debated topic in the literature [9,13,40–42]. While other data resulting from sequencing can be thought of as compositional, too, some groups in the microbiome data analysis community believe that compositionality has greater relevance in metagenomics due to the potential presence of dominant microbes. Here, we found that compositional methods did not outperform non-compositional methods designed for count data, indicating that their benefits did not outweigh the drawbacks they may introduce. This can be explained by two considerations. First, some compositional methods assume that the data arise from a multinomial distribution, with *n* trials (reads) and a vector *p* indicating the probability of the reads to be mapped to each OTU. In metagenomic studies, we have a large *n* (number of sequenced reads) and small *p* (since there are many OTUs, the probability of each read to map to any given OTU is small). In this setting, the Poisson distribution is a good approximation of the multinomial. Similarly, the negative binomial is a good approximation of the Dirichlet-Multinomial [31]. Secondly, some normalizations, such as the geometric mean method implemented in DESeq2 or the trimmed mean of M-values of edgeR, have size factors mathematically equivalent or very similar to the centered log-ratio proposed by Aitchison [40,43]. This has been shown to reduce the impact of compositionality on DA results [44]. We did not test the ANCOM package [45] because it was too slow for assessment. However, we included three recent analysis methods that address compositionality, namely, ALDEx2, songbird, and mixMC. This allowed us to perform an adequate assessment of compositional vs non-compositional approaches. Similarly, multivariate methods, such as songbird and mixMC, did not outperform methods based on univariate tests, suggesting that these simpler approaches are often sufficient to detect the most relevant biological signals.

The lack of ground truth makes the assessment of DA correctness very challenging. However, we can rely on mock datasets, within-method concordance, and enrichment analysis to obtain a principled ranking of method performances (Fig. 7). Although each analysis by itself does not imply correctness, taken together these assessments are a good proxy to evaluate methods performances in terms of their ability to limit the amount of false discoveries, give replicable results in datasets contrasting the same groups, and identify as significant the taxa that are expected to be DA.

**Figure 7:**
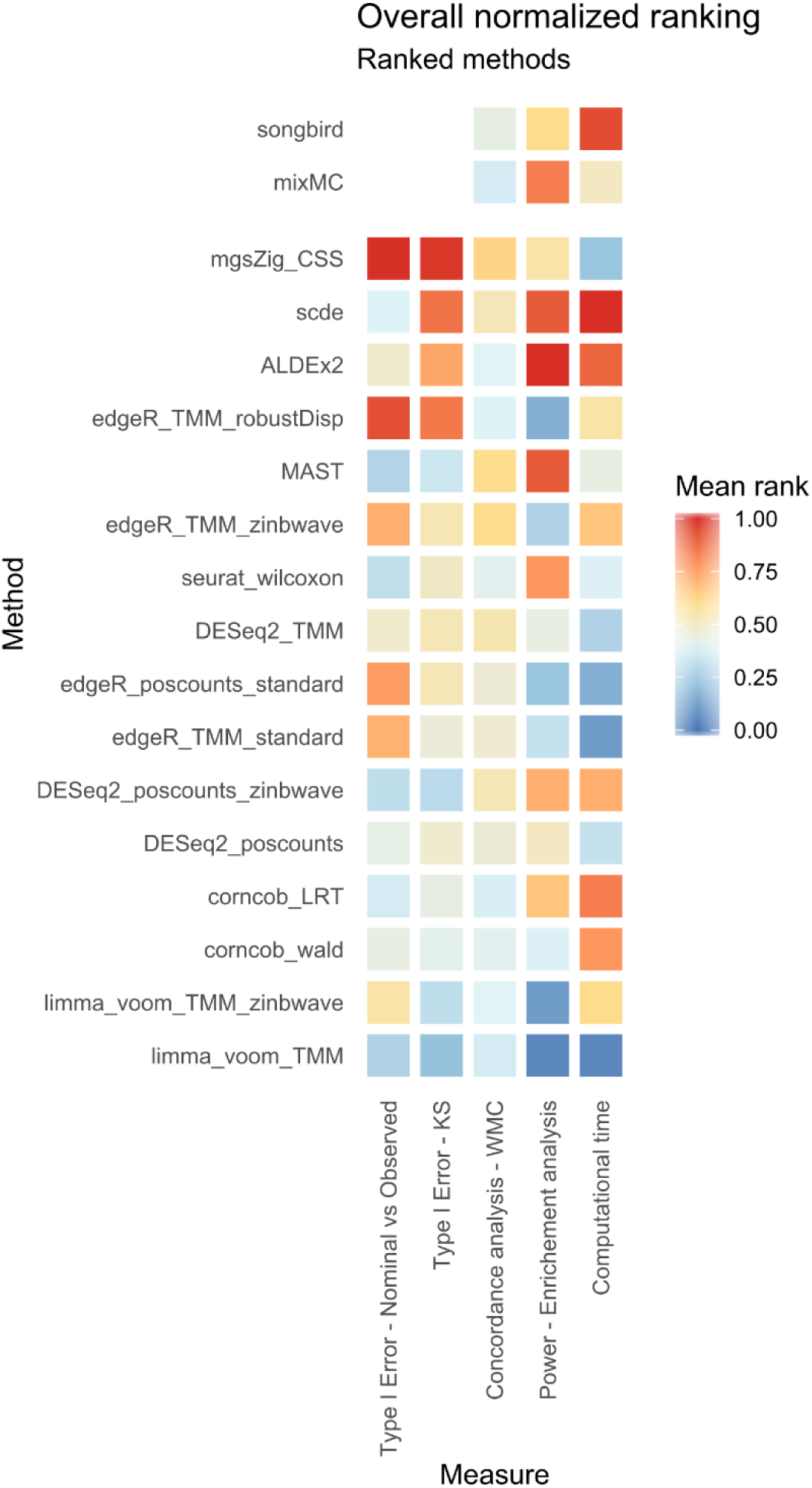
Overall method ranking regarding 5 evaluation criteria. Average normalized ranks range from 0 to 1, lower values correspond to better performances. Type I Error rows are based on the analysis of the 1000 mock comparisons from HMP 16S and WMS Stool dataset, Concordance analysis row is based on the average of WMC values obtained averaging performances in the 100 random subset comparisons for each of the 6 used datasets. Instead, Power - Enrichment analysis and Computational time columns, are based on the Supragingival vs Subgingival Plaque 16S dataset evaluations. Method’s ordering is computed using the first 4 column values. Because Type I error analysis was not available for songbird and mixMC, these methods were not included in the final ranking.

The parametric simulation framework is useful to inspect how individual characteristics of the data-generating distribution impact the sensitivity and specificity of the methods. As the entire analysis was supported by real data, we decided to focus only on a very simple but easily reproducible implementation of the NB and ZINB distributions for the simulations. The choice was justified by our GOF analysis on real datasets. Unsurprisingly, the sample size and the effect size were the characteristics that had the most impact on method performances. This translates into an evident suggestion for experimental design: large sample sizes are needed when dealing with low effect sizes. Our simulation framework can in principle be used for power calculations in the context of DA analysis.

In the 16S dataset used for the enrichment analysis, with a total of 76 samples and almost 900 unique taxa, the most time-consuming methods were scde and songbird with more than 5 minutes needed to identify DA taxa. ALDEx2 and corncob-based methods took about 40 seconds, zinbwave weighted methods took approximately 20 seconds while mixMC, MAST and seurat_wilcoxon around 10 seconds. DESeq2 and edgeR were under the 10 seconds with limma-voom which was the fastest method with less than a second (Fig. 7). A consistent ranking was found in simulated datasets with interesting changes determined by different sample-sizes (Additional file 2: Supplementary Table S5 and Supplementary Fig. S10).

## Conclusions

As already noted in recent publications [10–12], the perfect method does not exist. However, taken together, our analyses suggested that limma-voom, corncob, and DESeq2 showed the most consistent performance across all datasets, metagenomeSeq had the worst performance, and scde and ALDEx2 suffered from low power (Fig. 7). Among compositional data analysis methods, songbird showed a greater ability in identifying the correct taxa in the enrichment analysis, while mixMC had a better within-method concordance.

In general, we recommend a careful exploratory data analysis and we present a framework that can help scientists make an informed choice in a dataset-specific manner. In this study, we did not find evidence that bespoke differential abundance methods outperform methods developed for the differential expression analysis of RNA-seq data. However, our analyses also suggested that further research is required to overcome the limitations of currently available methods: in this respect, new directions in DA method development, e.g., leveraging the phylogenetic tree [46,47], log-contrast models [48], or compositional balances [49] are promising, but efforts to make these methods scalable are needed.

## Methods

### Datasets

The *HMP16Sdata* [25] (v1.2.0) and *CuratedMetagenomicData* [26] (v1.12.3) Bioconductor packages are used to download high-quality, uniformly processed, and manually annotated human microbiome profiles for thousands of people, using 16S and Whole Metagenome shotgun sequencing technologies respectively. *HMP16SData* comprises the collection of 16S data from the Human Microbiome Project (HMP), while *CuratedMetagenomicData* contains data from several projects. Gene-level counts for a collection of public scRNA-seq datasets are downloaded from *scRNAseq* (v 1.99.8) Bioconductor package.

While the latter datasets are used only for a comparison between technologies, the former are widely used for all the analyses. A complete index with dataset usage is reported in Additional file 1: Supplementary Table S1.

*Phyloseq* objects were obtained from the *HMP16SData* and *curatedMetagenomicData* packages using the function *as_phyloseq()* and setting the *bugs.as.phyloseq = TRUE* argument, respectively. The *otu_table* and *sample_data* slots of the *phyloseq* objects that contain, respectively, the taxa count table and the metadata associated to each sample were used for all downstream analyses. For the WMS datasets, absolute raw count data were estimated from the *metaPhlAn2-*produced relative count data by multiplying the columns of the *ExpressionSet* data by the number of reads for each sample, as found in the *pData* column *“number_reads”* (*counts = TRUE* argument).

*HMP16SData* was split by body subsite in order to obtain 18 separated datasets. Stool and Tongue Dorsum datasets were selected for example purposes thanks to their high sample size. The same was done on *CuratedMetagenomicData* HMP dataset, obtaining 9 datasets. Moreover, for the evaluation of type I error control, 41 stool samples with equal RSID, in both 16S and WMS, were used to compare DA methods. For each research project, *CuratedMetagenomicData* was split by body site and treatment or disease condition, in order to create homogeneous sample datasets. A total of 82 WMS datasets were created.

A total of 100 datasets were evaluated, however for the CAT analysis, non-split by condition or body subsite datasets were evaluated (e.g. Tongue Dorsum vs Stool in HMP, 2012 for both 16S and WMS).

To consider the complexity and the variety of several experimental scenarios, an attempt to select a wide variety of datasets for the analysis was done. The datasets were chosen based on several criteria: the sample size, the homogeneity of the samples or the availability of the same RSID for both technologies.

### Statistical Models

The following distributions were fitted to each dataset, either by directly modeling the read counts, or by first applying a logarithmic transformation:

- Negative Binomial (NB) model, as implemented in the *edgeR* (v3.24.3) Bioconductor package (on read counts);
- Zero Inflated Negative Binomial (ZINB), as implemented in the *zinbwave* (v1.4.2) Bioconductor package (on read counts);
- Truncated Gaussian hurdle model, as implemented in the *MAST* (v1.8.2) Bioconductor package (on log count);
- Zero Inflated Gaussian (ZIG), as implemented in the *metagenomeSeq* (v1.24.1) Bioconductor package (on log count).
- Dirichlet-Multinomial (DM), as implemented in the *MGLM* (v0.2.0) CRAN R package.

### Negative Binomial (NB)

We used the implementation of the NB model of the *edgeR* Bioconductor package. In particular, normalization factors were calculated with the Trimmed Mean of M-values (TMM) normalization [50] using the *calcNormFactors* function; common, trended and tagwise dispersions were estimated by *estimateDisp*, and a negative binomial generalized log-linear model was fit to the read counts of each feature, using the *glmFit* function.

### Zero-Inflated Negative Binomial (ZINB)

We used the implementation of the ZINB model of the *zinbwave* Bioconductor package. We fitted a ZINB distribution using the *zinbFit* function. As explained in the original paper, the method can account for various known and unknown, technical and biological effects [23]. However, to avoid giving unfair advantages to this method, we did not include any latent factor in the model (K = 0). We estimated a common dispersion for all features (common_dispersion = TRUE) and we set the likelihood penalization parameter epsilon to 1e10 (within the recommended set of values [24]).

### Truncated Gaussian Hurdle Model

We used the implementation of the *MAST* Bioconductor package. After a log2 transformation of the reascaled counts with a pseudocount of 1, a zero-truncated Gaussian distribution was modeled through generalized regression on positive counts, while a logistic regression modeled feature expression/abundance rate. As suggested in the MAST paper [7], cell detection rate (CDR) which is computed as the proportion of positive count features for each sample, was added as a covariate in the discrete and continuous model matrices as normalization factor.

### Zero-Inflated Gaussian

*metagenomeSeq* Bioconductor package was used to implement a ZIG model for log2 transformed counts with a pseudocount of 1, rescaled by the median of all normalization factors or by 1e03 which gives the interpretation of “count per thousand” to the offsets. The *CumNormStat* and *CumNorm* functions were used to perform Cumulative Sum Scaling (CSS) normalization, which accounts for specific data characteristics. Normalization factors were included in the regression through the *fitZig* function.

Note that both *MAST* and *metagenomeSeq* are applied to the normalized, log-transformed data. We evaluated both models, using their default scale factor 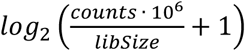 for *MAST* and 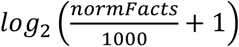 for *metagenomeSeq*, as well as by rescaling the data to the median library size [13], 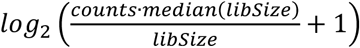 and 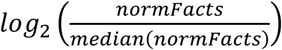, respectively.

### Dirichlet-Multinomial

The *MGLM* package was used to fit a Dirichlet-Multinomial regression model for counts. The *MGLMreg* function with *dist = “DM”*, allowed the implementation of the above model and the estimation of the parameter values.

### Goodness of Fit (GOF)

To evaluate the goodness of fit of the models, we computed the mean differences between the estimated and observed values for several datasets.

For each model, we evaluated two distinct aspects: its ability to correctly estimate the mean counts (plotted in logarithmic scale with a pseudo-count of 1) and its ability to correctly estimate the probability of observing a zero, computed as the difference between the probability of observing a zero count according to the model and the observed zero frequencies (Zero Probability Difference, ZPD). We summarized the results by computing the Root Mean Squared Error (RMSE) of the two estimators. The lower the RMSE, the better the fit of the model.

This analysis was repeated for 100 datasets available in *HMP16SData* and *CuratedMetagenomicData* (Table S1 and Additional file 1: Supplementary Figure S2).

Assuming homogeneity between samples inside the same body subsite or study condition, we specified a model consisting of only an intercept, or including a normalization covariate.

### Differential abundance detection methods

#### DESeq2

The *DESeq2* (v1.22.2) Bioconductor package fits a negative binomial model for count data. DESeq2 default data normalization is the so-called Relative Log Expression (RLE) based on scaling each sample by the median ratio of the sample counts over the geometric mean counts across samples. As 16S and WMS data sparsity may lead to a geometric mean of zero, it is replaced by n-th root of the product of the non-zero counts (which is the geometric mean of the positive count values) as proposed in *phyloseq* package [51] and implemented in the DESeq2 *estimateSizeFactors* function with option *type=“poscounts”*. We also tested DESeq2 with TMM normalization (see below). Moreover, as proposed in Van den Berge et al. [24], observational weights are supplied in the *weights* slot of the *DESeqDataSet* class object to account for zero inflation. Observational weights were computed by the *ComputeObservationalWeights* function of the *zinbwave* package. To test for DA, we used a Likelihood Ratio Test (LRT) to compare the reduced model (intercept only) to the full model with intercept and group variable. The p-values were adjusted for multiple testing via the Benjamini-Hochberg (BH) procedure. Some p-values were set to NA via the *cooksCutoff* argument that prevents rare or outlier features from being tested.

#### edgeR

The *edgeR* Bioconductor package fits a negative binomial distribution, similarly to DESeq2. The two approaches differ mainly in the normalization, dispersion parameter estimation, and default statistical test. We examined different procedures by varying the normalization and the dispersion parameter estimation: *edgeR_TMM_standard* involves TMM normalization and tagwise dispersion estimation through the *calcNormFactors* and *estimateDisp* functions respectively (with default values). Analogously to DESeq2, “*poscounts”* normalization was used in addition to TMM in *edgeR_poscounts_standard* to investigate normalization impact. We also evaluated the impact of employing a robust dispersion estimation, accompanied with a quasi-likelihood F test through the *estimateGLMRobustDisp* and *glmQLFit* functions respectively (*edgeR_TMM_robustDisp)*. As with DESeq2, *zinbwave* observational weights were included in the *weights* slot of the *DGEList* object in *edgeR_TMM_zinbwave* to account for zero inflation, through a weighted F test. Benjamini-Hochberg correction was used to adjust p-values for multiple testing.

#### Limma-voom

The *limma* Bioconductor package (v3.38.3) includes a *voom* function that (i) transforms previously normalized counts to logCPM, (ii) estimates a mean-variance relationship and (iii) uses this to compute appropriate observational-level weights[21]. To adapt *limma-voom* framework to zero-inflations, *zinbwave* weights have been multiplied by *voom* weights as done previously [24]. The residual degrees of freedom of the linear model were adjusted before the empirical Bayes variance shrinkage and were propagated to the moderated statistical tests. Benjamini-Hochberg correction method was used to correct p-values.

#### ALDEx2

*ALDEx2* is a Bioconductor package (v1.14.1) that uses a Dirichlet-multinomial model to infer abundance from counts [14]. The *aldex* method infers biological and sampling variation to calculate the expected False Discovery Rate, given the variation, based on several tests. Technical variation within each sample is estimated using Monte-Carlo draws from the Dirichlet distribution. This distribution maintains the proportional nature of the data while scale-invariance and sub-compositionally coherence of data, is ensured by centered log-ratio (CLR). This removes the need for a between sample normalization step. In order to obtain symmetric CLRs, the *iqlr* argument is applied, which takes, as the denominator of the log-ratio, the geometric mean of those features with variance calculated from the CLR between the first and the third quantile. Statistical testing is done through Wilcoxon Rank Sum test, even if Welch’s t, Kruskal-Wallis, Generalized Linear Models and correlation tests were available. Benjamini-Hochberg correction method was used to correct the p-values for multiple testing.

#### metagenomeSeq

*metagenomeSeq* is a Bioconductor package designed to address the effects of both normalization and under-sampling of microbial communities on disease association detection and testing feature correlations. The underlying statistical distribution for *log*_2_(*count* + 1) is assumed to be a zero-inflated Gaussian mixture model. The mixture parameter is modeled through a logistic regression depending on library sizes, while the Gaussian part of the model is a generalized linear model with a sample specific intercept which represent the sample baseline, a sample specific offset computed by Cumulative Sum Scaling (CSS) normalization and another parameter which represents the experimental group of the sample. We opted for the implementation suggested in the original publication [13], where CSS scaling factors are divided by the median of all the scaling factors instead of dividing them by 1000 (as done in the Bioconductor package). An EM algorithm is performed by *fitZig* function to estimate all parameters. An empirical Bayes approach is used for variance estimation and a moderated t-test is performed to identify differentially abundant features between conditions. Benjamini-Hochberg correction method was used to account for multiple testing.

#### Corncob

*corncob* is an R package (v0.1.0 [52]) for the differential abundance and differential variability analysis of microbiome data [17]. Specifically, *corncob* is designed to account for the challenges of modeling sequencing data from microbial abundance studies. It is based on a hierarchical model in which the latent relative abundance of each taxon is modelled as a beta distribution, and the observed absolute presence of a taxon is modelled as a binomial process with the previously specified beta as the probability of success. This hierarchical structure gives flexibility to the method, which can account for changes in the average count values as well as their dispersion. A generalized linear model framework, with a logit link function, is used to allow the study of covariates in the feature count distributions. The model fit is performed by maximum likelihood using the trust region optimization algorithm [17]. Likelihood-ratio or Wald tests can be used to test the null hypothesis of no DA.

#### Songbird

*songbird* is a python package [53] that ranks microbes that are changing the most relative to each other [16]. The method is based on a compositional approach in which the underlying count distribution is assumed to be multinomial. The coefficients from multinomial regression can be ranked to determine which taxa are changing the most between samples. The compositionality is addressed using the differential abundance of each taxon as reference to each other when they are ranked numerically. Since *songbird* has been developed as an extension tool for *Qiime2*, we converted all our data tables to the *biom* format to serve as input for this method. The authors’ suggested analysis pipeline requires several manual adjustments to the tuning parameters on the basis of the comparison of the results after several runs, making it difficult to implement this method within a benchmarking framework. For this reason, we used the default values for all the tuning parameters.

#### mixMC

*mixMC* is a multivariate framework implemented in mixOmics, a Bioconductor package (v6.6.1), for microbiome data analysis [18]. It handles compositional and sparse data, repeated-measures experiments and multiclass problems. After the addition of a pseudo-count value of 1, the TSS normalization is applied to the count table and the CLR transformation is performed to account for compositionality. The method is based on a Partial Least Squares (PLS) Discriminant Analysis (DA), a multivariate regression model which maximises the covariance between linear combinations of the feature counts and the outcome (in our case, a dummy matrix indicating the body site/group of each sample). Covariance maximization is achieved in a sequential manner via the use of latent component scores [18]. Each component is a linear combination of the feature counts and characterizes a source of covariation between the feature and the groups. The sparse version of PLS-DA, sPLS-DA uses Lasso penalizations to select the most discriminative features in the PLS-DA model. The penalization is applied component wise and the resulting selected features reflect the particular source of covariance in the data highlighted by each PLS component. We specified the number of features to select per component at 100 or more, and we optimized it using leave-one-out cross-validation. Since we always compared two groups in this manuscript, only the first component is necessary for the analysis. The multivariate regression coefficients, one for each feature, were ranked in order to obtain the most discriminant features for the first component.

#### MAST

*MAST* is a Bioconductor package for managing and analyzing qPCR and sequencing-based single-cell gene expression data, as well as data from other types of single-cell assays. The package also provides functionality for significance testing of differential expression using a Hurdle model. Zero rate represents the discrete part, modelled as a binomial distribution while 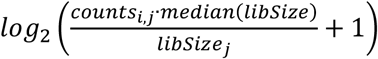 where i and j represents the i-th feature and the j-th sample respectively, is used for the continuous part, modelled as a Gaussian distribution. The kind of data considered, different from scRNA-seq, doesn’t allow the usage of the adaptive thresholding procedure suggested in the original publication [7]. Indeed, because of the amount of feature loss if adaptive thresholding is applied, the comparison of MAST with other methods would be unfair. However, a normalization variable is included in the model. This variable captures information about each feature sparsity related to all the others; hence, it helps to yield more interpretable results and decreases background correlation between features. The function *zlm* fits the Hurdle model for each feature: the regression coefficients of the discrete component are regularized using a Bayesian approach as implemented in the *bayesglm* function; regularization of the continuous model variance parameter helps to increase the robustness of feature-level differential expression analysis when a feature is only present in a few samples. Because the discrete and continuous parts are defined conditionally independent for each feature, tests with asymptotic χ^2^ null distributions, such as the Likelihood Ratio or Wald tests, can be summed and remain asymptotically χ^2^, with the degrees of freedom of the component tests added. Benjamini-Hochberg correction method was used to correct p-values.

#### Seurat with Wilcoxon Rank Sum Test

*Seurat* (v2.3.4) R package is a scRNA-Seq data analysis toolkit for the analysis of single-cell RNA-seq [22]. Briefly, counts were scaled, centered and LogNormalized. Wilcoxon Rank-Sum test for detecting differentially abundant features was performed via the *FindMarkers* function. Rare features, which are present in a fraction lower than 0.1 of all samples, and weak signal features, which have a log fold change between conditions lower than 0.25, are not tested. Benjamini-Hochberg correction method was used to correct p-values.

#### SCDE - Single Cell Differential Expression

The *scde* Bioconductor package (v1.99.1) with *flexmix* package (v2.3-13) implements a Bayesian model for scRNA-seq data [8]. Read counts observed for each gene are modeled using a mixture of a negative binomial (NB) distribution (for the amplified/detected transcripts) and low-level Poisson distribution (for the unobserved or background-level signal of genes that failed to amplify or were not detected for other reasons). The *scde.error.models* function was used to fit the error models on which all subsequent calculations rely. The fitting process is based on a subset of robust genes detected in multiple cross-cell comparisons. Error models for each group of cells were fitted independently (using two different sets of “robust” genes). Translating in a metagenomic context, cells correspond to samples and genes to OTU or amplicon sequence variants. Some adjustments were needed to calibrate some function default values such as the minimum number of features to use when determining the expected abundance magnitude during model fitting. This option, defined by the *min.size.entries* argument, set by default at 2000, was too big for many 16S or WMS experiment scenarios: as we usually observe around 1000 total features per dataset (after filtering out rare ones), we decided to replace 2000 with the 20% of the total number of features, obtaining a dataset-specific value. Particularly poor samples may result in abnormal fits and were removed as suggested in the *scde* manual. To test for differential expression between the two groups of samples a Bayesian approach was used: incorporating evidence provided by the measurements of individual samples, the posterior probability of a feature being present at any given average level in each subpopulation was estimated. To moderate the impact of high-magnitude outlier events, bootstrap resampling was used and posterior probability of abundance fold-change between groups was computed.

#### Type I error control

For this analysis, we used the collection of HMP Stool samples in *HMP16SData* and *CuratedMetagenomicData*. The multidimensional scaling (MDS) plot of the beta diversity did not show patterns associated with known variables (Additional file 1: Supplementary Fig. S3), hence we assumed no differential abundance. All samples with the same Random Subject Identifier (RSID) in 16S and WMS were selected in order to easily compare the two technologies. 41 biological samples were included.

Starting from the 41 samples, we randomly split the samples in two groups: 21 assigned to Group 1 and 20 to Group 2. We repeated the procedure 1000 times. We applied the DA methods to each randomly split dataset. Every method returned a p-value for each feature. DESeq2, seurat_wilcoxon and corncob methods returned some NA p-values. This is due to feature exclusion criteria, based on distributional assumptions, performed by these methods (see above), or convergence issues.

We compared the distribution of the observed p-values to the theoretical uniform distribution, as no differential abundant features should be present. This was summarized in the qq-plot where the bisector represents a perfect correspondence between observed and theoretical quantiles of p-values. For each theoretical quantile, the corresponding observed quantile was obtained averaging the observed p-values’ quantiles from all 1000 datasets. Departure from uniformity was evaluated with a Kolmogorov-Smirnov statistic. P-values were also used to compare the number of false discoveries with 3 common thresholds: 0.01, 0.05 and 0.1.

#### Concordance

We used the Concordance At the Top (CAT) to evaluate concordance for each differential abundance method. Starting from two lists of ranked features (by p-values, fold-changes or other measures), the CAT statistic was computed in the following way. For a given integer *i*, concordance is defined as the cardinality of the intersection of the top *i* elements of each list, divided by *i*, i.e. 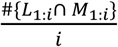, where *L* and *M* represent the two lists. This concordance was computed for values of *i* from 1 to *R*.

Depending on the study, only a minority of features may be expected to be differentially abundant between two experimental conditions. Hence, the expected number of differentially abundant features is a good choice as the maximum rank *R*. In fact, CAT displays high variability for low ranks as few features are involved, while concordance tends to 1 as *approaches* the total number of features, becoming uninformative. We set *R* = 100, considering this number biologically relevant and high enough to permit an accurate concordance evaluation. In our filtered data, the total number of features was close to 1000, and 100 corresponds to 10% of total taxa.

We used CAT for two different analyses:

- Between Method Concordance (BMC), in which a method was compared to other methods in the same dataset;
- Within Method Concordance (WMC), in which a method is compared to itself in random splits of the datasets.

To summarize this information for all pairwise method comparisons, we computed the Area Under the Curve, hence giving a better score to two methods that are consistently concordant for all values of *i* from 1 to 100.

We selected several datasets, with different alpha and beta diversity, for our concordance analysis. Table S3 describes the six datasets used. For each dataset, the same sample selection step, described next, was used.

The concordance evaluation algorithm can be easily summarized by the following steps:

1. Each dataset was randomly divided in half to obtain two subsets (Subset1 and Subset2) with two balanced groups;
2. DA analysis between the groups was performed with all evaluated methods independently on each subset;
3. For each method, the list of features ordered by p-values (or differentials, or loadings) obtained from Subset1 was compared to the analogous list obtained from Subset2 and used to evaluate WMC;
4. For each method, the list of features ordered by p-values (or differentials, or loadings) obtained from Subset1 was compared to the analogous list obtained from Subset1 by all the other methods and used to evaluate BMC for Subset1. The same was done in Subset2.
5. Steps 1-4 were repeated 100 times;
6. WMC and BMC were averaged across the 100 values (and between Subset1 and Subset2 for BMC) to obtain the final values.

#### Sample selection step

For each dataset, a subset was chosen in order to have a balanced number of samples for each condition. In lower diversity studies (e.g. Subgingival vs Supragingival Plaque) different biological samples from the same subject may be strongly correlated. Hence, we selected only one sample per individual, no matter the condition. To further increase the homogeneity of the datasets, we selected only samples from the same sequencing center.

#### Enrichment analysis

The same low-diversity dataset used in the concordance analysis (i.e., 16S Subgingival vs Supragingival Plaque), was used for the enrichment analysis. The dataset is balanced as it is composed of 38 samples for each body subsite, for a total of 76 samples. DA analysis was performed using Subgingival Plaque as the reference level. Taxa with an adjusted p-value less than 0.1 were chosen as DA, for all the methods except songbird and mixMC that return a list of differentials and loadings, respectively. For songbird, a threshold corresponding to the 10% of the total number of taxa was chosen to select the most associated taxa for the considered comparison. mixMC implements a variable selection procedure that automatically selects the most discriminant taxa. We annotated each taxon with the information on genus-level metabolism (available at https://github.com/waldronlab/nychanesmicrobiome), classifying each taxon in aerobic, anaerobic, facultative anaerobic, or unassigned.

Enrichment analysis was performed via a Fisher exact test, using the function *fisher.test(table, alternative = “greater”*) where *table* is a contingency table. Six contingency tables were built for each method to inspect enrichment of:

- over-abundant (UP) aerobic taxa in Supragingival Plaque;
- under-abundant (DOWN) aerobic taxa in Subgingival Plaque;
- over-abundant (UP) anaerobic taxa in Supragingival Plaque;
- under-abundant (DOWN) anaerobic taxa in Subgingival Plaque;
- over-abundant (UP) facultative anaerobic taxa in Supragingival Plaque;
- under-abundant (DOWN) facultative anaerobic taxa in Subgingival Plaque.

All the information retrieved from the enrichment analysis were summarized in a bar plot, where for each method, the number of differentially abundant taxa together with their direction were represented as a positive (negative) bar for over- (under-) abundant taxa in Supragingival Plaque samples, colored by genus level metabolism.

To calculate log odds-ratio for each contingency table, the Haldane-Anscombe correction is applied since it allows the odds-ratio calculation in presence of zero cells. Briefly, it consists in adding a pseudo-count value of 0.5 to each cell of the contingency table to calculate the odds-ratio and a pseudo-count value of 1 to calculate the variance.

To compare all the evaluated methods without considering their power, the followings steps were followed:

1. raw p-values, songbird’s differentials and mixMC’s loadings were properly ordered;
2. several thresholds from 1% to 20% of the top ranked taxa in the previously ordered lists were used to select the DA taxa for each method;
3. Putative true positives (TP) were calculated as the sum of Aerobic taxa over-abundant in Supragingival Plaque and Anaerobic taxa under-abundant in Supragingival Plaque;
4. Putative false positives (FP) were calculated as the sum of Aerobic taxa under-abundant in Supragingival Plaque and Anaerobic taxa over-abundant in Supragingival Plaque;
5. The differences between Putative TP and Putative FP were plotted

To rank all the methods, the same difference was computed, this time using the list of DA taxa based on the adjusted p-values less than 0.1 and the 10% threshold for songbird.

To inspect the concordance of DA taxa between methods, mutual findings were collected and added between the methods. As similar methods tend to identify the same taxa, only one method for each normalization or weighting procedure was considered as representative. This subset contains: edgeR with TMM normalization, DESeq2 with poscounts normalization, limma-voom with TMM normalization, MAST, scde, seurat-wilcoxon, corncob (Wald test), mgsZig, ALDEx2, mixMC and songbird. The taxa found by most methods in this subset were extracted, but for the graphical representation, all methods were reintroduced.

The same analysis was performed in the WMS dataset. However, the sample size was limited to only 5 for the subgingival body subsite, while 88 (with unique RSID) for the supragingival site. For this reason, a 5 vs 5 samples analysis was performed, randomly selecting five samples from the supragingival dataset. Songbird was not included in the analysis because of an error during the parameter estimation that we were not able to solve. Given the low sample-size, corncob methods with bootstrap were added to the analysis.

#### Parametric simulations

Several real datasets were used as templates for the simulations:

- 41 Stool samples available for both 16S and WMS from HMP;
- 208 16S samples and 90 WMS samples of Tongue Dorsum body subsite from HMP.
- 67 Stool and 56 Oral cavity WMS data of Fijian adult women from BritoIl_2016.

Each dataset was filtered to obtain only a sample per individual. 16S and WMS samples were pruned to keep sequencing runs with library sizes of more than 10^3^ and 10^6^, respectively. Moreover, only features present in more than 1 sample with more than 10 reads were kept. After the data filtering step, the simulation framework was established, by specifying the parametric distribution and other data characteristics, described in Additional file 2: Supplementary Table S4.

For each combination of parameters, we simulated 50 datasets, yielding a total of 28,800 simulations. Variables to be included in the simulation framework were chosen based on the role they may play in the analysis of a real experiment.

NB and ZINB are simple parametric distributions, easy to fit on real data through a reliable Bioconductor package and above all, seemed to fit 16S and WMS data better than other statistical models (see Figure 2). The *zinbSim* function from the *zinbwave* Bioconductor package easily allows the user to generate both NB and ZINB counts after the *zinbFit* function estimates model parameters from real data. The user can set several options in *zinbFit*, we used *epsilon=1e14, common_dispersion=TRUE*, and *K=0*.

Generating two experimental groups requires the specification of enough samples for each condition and a more or less substantial biological difference between them.

Sample size is a crucial parameter: many pilot studies start with 10 or even fewer samples per condition, while clinical trials and case-control studies may need more samples in order to achieve the needed power. We included 10, 20 and 40 samples per condition in our simulation framework.

We considered two different scenarios for the number of features simulated as DA: 10%, representing a case where the majority of the features are not DA, a common assumption made by analysis methods; and 50%, a more extreme comparison. Similarly, we simulated a fold change difference for the DA features of 2 or 5. This is obviously a simplification, since in reality a continuum gradient of fold effects is present. Nevertheless, it allowed us to characterize the role of the effect size in the performance of the methods. For the DA features, the fold change between conditions was applied to the mean parameter of the ZINB or NB distributions, with or without “compensation” as introduced by Hawinkel et al [10]. Without compensation, the absolute abundance of a small group of features responds to a physiological change. This simple procedure modifies the mean relative abundances of all features, a microbiologist would only want to detect the small group that initially reacted to the physiological change. For this reason, significant results for other features will be considered as false discoveries. Compensation prevents the changes in DA features to influence the other, non-DA, features. The procedure comprises the following steps:

1. The relative mean for each feature is computed using estimated mean parameter of NB;
2. 10% or 50% of features are randomly sampled;
3. If there is no compensation, half of their relative means are multiplied by *foldEffect* while the remainings are divided by *foldEffect* generating up and down regulated features respectively. If there is compensation, 1/(1+*foldEffect*) of the selected feature relative means are multiplied by *foldEffect* while the remaining ones are multiplied by *(a/b)*(1-foldEffect)+1*, where *a* is the sum of the relative means of the features that will be up-regulated while *b* is the sum of the features that will be down-regulated.
4. The resulting relative means are normalized to sum to 1.

Sparsity is a key characteristic of metagenomic data. The case in which a bacterial species presence rate varies between conditions was emulated in the simulation framework via the so called *sparsityEffect* variable. Acting on the mixture parameter of the ZINB model it is possible to exacerbate down-regulation and up-regulation of a feature, adding zeros for the former and reducing zeroes for the latter. This scenario provided by 0 (no sparsity change at all), 0.05 and 0.15 of sparsity change should help methods to identify more differentially abundant features. As the mixing parameter can only take values between 0 and 1, when the additive sparsity effect yielded a value outside this range, it was forced to the closer limit.

The previously described DA methods were tested in each of the simulated datasets (50 for each set of simulation framework parameters) and the adjusted p-values were used to compute the False Positive Rate (FPR = 1 - Specificity) and the True Positive Rate (TPR = Sensitivity). Partial areas under the Receiver Operating Characteristic (pAUROC) curve with an FPR from 0 to 0.1 values were computed and then averaged in order to obtain a single value for each set of variables.

#### Computational complexity

To measure the computational times for all the 18 methods, we used the Subgingival vs Supragingival Plaque HMP 16S dataset where a total of 76 samples and approximately 900 taxa were available. The evaluation was performed on a laptop computer with O.S. Windows 10 64bit, Intel® i7-8th Gen CPU with 16GB of RAM. Moreover, the Stool 16S and WMS parametric simulation datasets (9200 total datasets), were used in order to measure each method’s computational complexity (except for mixMC and songbird). Time evaluation was performed on a single core for each dataset where all methods are tested sequentially and then properly averaged with the values of all the simulations. The methods’ performance evaluations in power analysis on the 28800 total parametric simulations were performed in the same way, equally dividing the simulated datasets across 30 cores. The working machine was a Linux x86_64 architecture server with 2 Intel® Xeon® Gold 6140 CPU with 2.30 GHz for a total of 72 CPUs and 128 GB of RAM.

## Declarations

### Ethics approval and consent to participate

Not applicable.

### Consent for publication

Not applicable.

### Availability of data and materials

The real datasets used in this article are available in the *HMP16SData* Bioconductor package, available at http://bioconductor.org/packages/HMP16SData, and in the *CuratedMetagenomicData* Bioconductor packages, available at http://bioconductor.org/packages/CuratedMetagenomicData. The scripts to reproduce all analyses and figures of this article are available at https://github.com/mcalgaro93/sc2meta

### Authors’ contributions

DR, CR, NV conceived the project, LW co-developed the evaluation strategies, MC and DR drafted the manuscript, LW CR NV reviewed and edited the manuscript, MC performed the data analyses and curated the code repository. All Authors read and approved the final manuscript.

