## Additional file 1 for "Assessment of statistical methods from single cell, bulk RNA-seq and metagenomics applied to microbiome data"

### SUPPLEMENTARY FIGURES AND TABLES

**Supplementary Table S1**: List of datasets used.

| | | | | | | | | Figures | | | | | |
|---|---|---|---|---|---|---|---|---|---|---|---|---|---|
| **Technology** | **Dataset name** | **Year** | **Bodysite** | **Condition/Subsite** | **Samples** | **Taxa** | **Sparsity** | **2** | **3** | **4** | **5** | **6** | **7** |
| 16S from HMP16SData | HMP | 2012 | Gastrointestinal Tract | Stool | 202 | 1959 | 0.76 | | | | | | |
| | | | Oral | Saliva | 185 | 1173 | 0.61 | | | | | | |
| | | | | Tongue Dorsum | 208 | 1133 | 0.60 | | | | | | |
| | | | | Hard Palate | 197 | 1046 | 0.62 | | | | | | |
| | | | | Buccal Mucosa | 200 | 936 | 0.63 | | | | | | |
| | | | | Attached Keratinized Gingiva | 207 | 1081 | 0.70 | | | | | | |
| | | | | Palatine Tonsils | 207 | 1478 | 0.69 | | | | | | |
| | | | | Throat | 196 | 1210 | 0.65 | | | | | | |
| | | | | Supragingival Plaque | 206 | 1414 | 0.65 | | | | | | |
| | | | | Subgingival Plaque | 205 | 1587 | 0.68 | | | | | | |
| | | | Airways | Anterior Nares | 169 | 798 | 0.68 | | | | | | |
| | | | Skin | Left Antecubital Fossa | 67 | 475 | 0.71 | | | | | | |
| | | | | Right Antecubital Fossa | 70 | 463 | 0.70 | | | | | | |
| | | | | Left Retroauricular Crease | 175 | 727 | 0.70 | | | | | | |
| | | | | Right Retroauricular Crease | 181 | 760 | 0.71 | | | | | | |
| | | | Urogenital Tract | Vaginal Introitus | 86 | 634 | 0.76 | | | | | | |
| | | | | Mid Vagina | 95 | 590 | 0.77 | | | | | | |
| | | | | Posterior Fornix | 95 | 582 | 0.77 | | | | | | |
| WMS from CuratedMetagenomicData | HMP | 2012 | nasalcavity* | Anterior Nares | 74 | 388 | 0.83 | | | | | | |
| | | | oralcavity | Tongue Dorsum | 90 | 783 | 0.53 | | | | | | |
| | | | | Buccal Mucosa | 84 | 708 | 0.62 | | | | | | |
| | | | | Supragingival Plaque | 88 | 811 | 0.56 | | | | | | |
| | | | | Subgingival Plaque | 5 | 423 | 0.20 | | | | | | |
| | | | skin | Left Retroauricolar Crease | 8 | 248 | 0.53 | | | | | | |
| | | | | Right Retroauricolar Crease | 15 | 265 | 0.63 | | | | | | |
| | | | stool* | Stool | 95 | 796 | 0.69 | | | | | | |
| | | | vagina* | Posterior Fornix | 42 | 281 | 0.84 | | | | | | |
| WMS from CuratedMetagenomicData | AsnicarF | 2017 | milk | control | 4 | 25 | 0.36 | | | | | | |
| | | | stool | | 10 | 373 | 0.59 | | | | | | |
| | Bengtsson-PalmeJ | 2015 | stool | control | 36 | 750 | 0.52 | | | | | | |
| | BritoIL | 2016 | oralcavity | control | 140 | 867 | 0.59 | | | | | | |
| | | | stool | | 172 | 1197 | 0.72 | | | | | | |
| | Castro-NallarE | 2015 | oralcavity | schizophrenia | 16 | 498 | 0.48 | | | | | | |
| | | | | control | 16 | 507 | 0.47 | | | | | | |
| | ChngKR | 2016 | skin | AD | 19 | 485 | 0.54 | | | | | | |
| | | | | control | 20 | 566 | 0.61 | | | | | | |
| | DavidLA | 2015 | stool | control | 33 | 518 | 0.77 | | | | | | |
| | | | | infectiousgastroenteritis | 14 | 345 | 0.63 | | | | | | |
| | FengQ | 2015 | stool | CRC | 46 | 990 | 0.61 | | | | | | |
| | | | | control | 61 | 902 | 0.64 | | | | | | |
| | | | | adenoma | 47 | 913 | 0.64 | | | | | | |
| | HanniganGD | 2017 | stool | adenoma | 27 | 370 | 0.66 | | | | | | |
| | | | | control | 28 | 407 | 0.64 | | | | | | |
| | | | | CRC | 27 | 418 | 0.66 | | | | | | |
| | Heitz-BuschartA | 2016 | stool | T1D | 10 | 487 | 0.43 | | | | | | |
| | | | | control | 10 | 463 | 0.41 | | | | | | |
| | HMP | 2012 | nasalcavity* | control | 74 | 388 | 0.83 | | | | | | |
| | | | oralcavity | | 90 | 814 | 0.58 | | | | | | |
| | | | skin | | 17 | 325 | 0.68 | | | | | | |
| | | | stool* | | 95 | 796 | 0.69 | | | | | | |
| | | | vagina* | | 42 | 252 | 0.83 | | | | | | |
| | KarlssonFH | 2013 | stool | IGT | 49 | 777 | 0.64 | | | | | | |
| | | | | control | 43 | 651 | 0.57 | | | | | | |
| | | | | T2D | 53 | 777 | 0.63 | | | | | | |
| | KosticAD | 2015 | stool | control | 15 | 402 | 0.57 | | | | | | |
| | LeChatelierE | 2013 | stool | control | 292 | 1215 | 0.74 | | | | | | |
| | LiJ | 2014 | stool | control | 260 | 1275 | 0.74 | | | | | | |
| | LiJ | 2017 | stool | control | 41 | 570 | 0.63 | | | | | | |
| | | | | pre-hypertension | 56 | 589 | 0.65 | | | | | | |
| | | | | hypertension | 99 | 768 | 0.72 | | | | | | |
| | LiSS | 2016 | stool | control | 5 | 375 | 0.20 | | | | | | |
| | | | | metabolic syndrome | 10 | 495 | 0.41 | | | | | | |
| | | | | FMT | 10 | 496 | 0.38 | | | | | | |
| | LiuW | 2016 | stool | control | 110 | 801 | 0.67 | | | | | | |
| | LomanNJ | 2013 | stool | STEC | 43 | 582 | 0.72 | | | | | | |
| | LoombaR | 2017 | stool | fatty liver | 86 | 826 | 0.71 | | | | | | |
| | LouisS | 2016 | stool | control | 16 | 382 | 0.47 | | | | | | |
| | NielsenHB | 2014 | stool | control | 236 | 1271 | 0.75 | | | | | | |
| | | | | IBD | 82 | 1072 | 0.70 | | | | | | |
| | Obregon-TitoAJ | 2015 | stool | control | 58 | 845 | 0.69 | | | | | | |
| | OhJ | 2014 | skin | control | 20 | 909 | 0.78 | | | | | | |
| | PasolliE | 2018 | stool | control | 112 | 850 | 0.72 | | | | | | |
| | QinJ | 2012 | stool | control | 174 | 1035 | 0.73 | | | | | | |
| | | | | T2D | 170 | 1090 | 0.73 | | | | | | |

| | | | | | | | |
|---|---|---|---|---|---|---|---|
| | QinN | 2014 | stool | control | 114 | 832 | 0.65 |
| | | | | cirrhosis | 123 | 1192 | 0.71 |
| | RampelliS | 2015 | stool | control | 38 | 517 | 0.65 |
| | RaymondF | 2016 | stool | control | 24 | 564 | 0.52 |
| | | | | cephalosporins | 18 | 446 | 0.53 |
| | SchirmerM | 2016 | stool | control | 471 | 958 | 0.74 |
| | ShiB | 2015 | oralcavity | periodontitis | 12 | 503 | 0.28 |
| | | | | SRP | 12 | 524 | 0.36 |
| | SmitsSA | 2017 | stool | control | 40 | 250 | 0.49 |
| | TettAJ | 2016 | skin | control | 26 | 383 | 0.75 |
| | | | | psoriasis | 26 | 243 | 0.78 |
| | ThomasAM | 2018 | stool | control | 24 | 505 | 0.55 |
| | | | | adenoma | 27 | 547 | 0.60 |
| | | | | CRC | 29 | 673 | 0.66 |
| | VatanenT | 2016 | stool | control | 201 | 913 | 0.81 |
| | | | | bronchitis | 14 | 379 | 0.56 |
| | | | | otitis | 59 | 550 | 0.70 |
| | | | | respiratoryinf | 6 | 261 | 0.38 |
| | VincentC | 2016 | stool | control | 90 | 909 | 0.74 |
| | | | | CDI | 8 | 434 | 0.44 |
| | VogtmannE | 2016 | stool | control | 52 | 822 | 0.59 |
| | | | | CRC | 52 | 920 | 0.62 |
| | WenC | 2017 | stool | AS | 97 | 834 | 0.68 |
| | XieH | 2016 | stool | control | 250 | 1198 | 0.72 |
| | YuJ | 2015 | stool | CRC | 75 | 1017 | 0.65 |
| | | | | control | 53 | 843 | 0.61 |
| | ZellerG | 2014 | stool | control | 66 | 991 | 0.63 |
| | | | | CRC | 91 | 1110 | 0.67 |
| | | | | adenoma | 42 | 875 | 0.60 |

[*] These datasets are reported twice but repeated for completeness.

### **Supplementary Table S2**: List of Differential abundance detection methods.

| R library | Description | Denomination | Input | Normalization | Dispersion | Weights | Test | Application |
|---|---|---|---|---|---|---|---|---|
| DESeq2 | Negative Binomial (NB) generalized linear model. Several normalizations used. | DESeq2_poscounts | counts | RLE poscounts | tagwise | none | Empirical Bayes + LRT | RNA-seq |
| | | DESeq2_TMM | | TMM | | | | |
| edgeR | | edgeR_TMM_standard | | | | | | |
| | | edgeR_poscounts_standard | | RLE poscounts | | | | |
| | | edgeR_TMM_RobustDisp | | TMM | Robust tagwise | | Quasi-likelihood F | |
| limma | Linear model with weights | limma_voom_TMM | log2((counts + 0.5) * 1e06 / (lib.size + 1)) | | none | voom | Empirical Bayes + moderated t | microArray and RNA-seq |
| ALDEx2 | Compositional approach - Monte-Carlo simulations from a Dirichlet distribution | ALDEx2 | log2(counts+0.5) | IQLR | none | none | t | Microbiome |
| metagenomeSeq | Zero-Inflated Gaussian (ZIG) mixture model | mgsZig_CSS | log2(counts+1) | CSS + In model | | none | Empirical Bayes + moderated t | |
| Corncob | Beta-Binomial regression model | corncob_LRT | counts | none | tagwise | none | LRT | |
| | | Corncob_wald | | | | | Wald | |
| Songbird | Compositional approach - Multinomial regression model | songbird | relative counts | CLR | none | reference frames | ranking | |
| mixMC | Compositional approach – sparse Partial Least Squares Discriminant Analysis (sPLS-DA) | mixMC | (counts + 1)/ lib.size | CLR | none | none | | |
| zinbwave + edgeR, DESeq2 or limma | Zero-Inflated Negative Binomial (ZINB) estimation of observational weights to use in DESeq2, edgeR and limma-voom. | DESeq2_poscounts_zinbwave | counts | RLE poscounts | tagwise | zinbwave | Empirical Bayes + LRT | scRNA-seq |
| | | edgeR_TMM_zinbwave | | TMM | | | Weighted F | |
| | | limma_voom_TMM_zinbwave | log2((counts + 0.5) * 1e06 / (lib.size + 1)) | | none | voom * zinbwave | Empirical Bayes + moderated t | |
| MAST | Truncated Gaussian hurdle model. | MAST | log2(counts*median(lib.size)/ lib.size + 1) | In model | | none | Empirical Bayes + LRT | |
| seurat | Data centered, scaled and filtered before performing a Wilcox test. | seurat_wilcoxon | counts | LogNormalize + centering + scaling | LogVMR | | Wilcoxon | |
| scde | Bayesian approach based on a Negative Binomial and Poisson mixture model. | scde | | none | none | | Bayes + bootstrap resampling | |

**Supplementary Table S3:** Metagenomic approach and microbial diversity of datasets used for concordance analysis.

| Microbial diversity/ technology | 16S (HMP16SData) | WMS (CuratedMetagenomicData) |
|---|---|---|
| **Low** | 38 Subgingival and 38 Supragingival Plaque samples sequenced in WUGC from the HMP data. | 66 healthy controls and 66 CRC patients (a random subset from 91) Stool samples [28]. |
| **Mid** | 36 Attached Keratinized Gingiva and 35 Buccal Mucosa samples sequenced in WUGC from HMP data. | 16 healthy controls and 16 Schizophrenic patients Oral Cavity samples [29]. |
| **High** | 39 Stool and 39 Tongue Dorsum samples sequenced in WUGC from HMP data. | 45 Stool and 45 Tongue Dorsum samples from HMP data. |

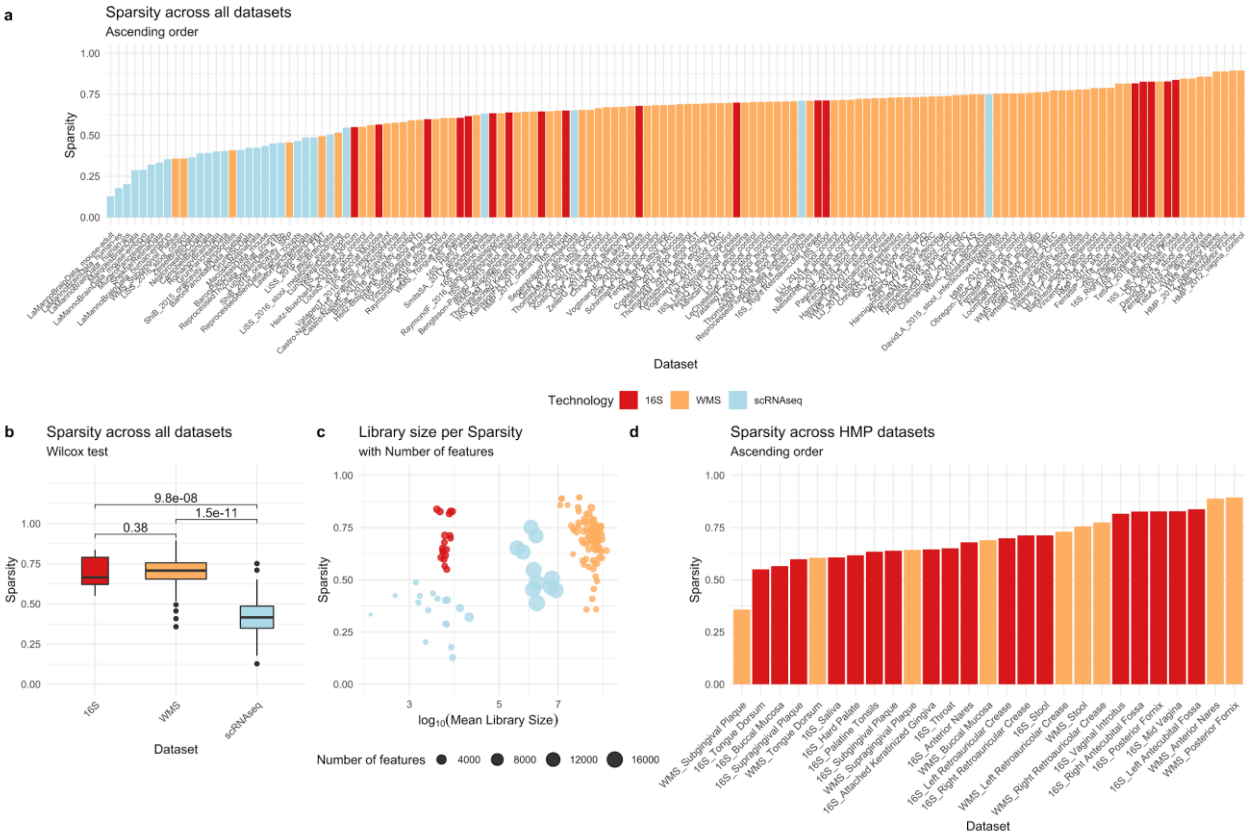

**Supplementary Figure S1**: **a.** Sparsity distribution across all 16S, WMS and scRNA-seq datasets. **b.** Boxplots and Wilcoxon tests for sparsity distributions between 16S, WMS and scRNA-seq technologies. **c.** Logarithm of mean library size vs. sparsity with point size proportional to the number of features for each dataset. **d.** Sparsity distribution for all HMP 16S and WMS datasets.

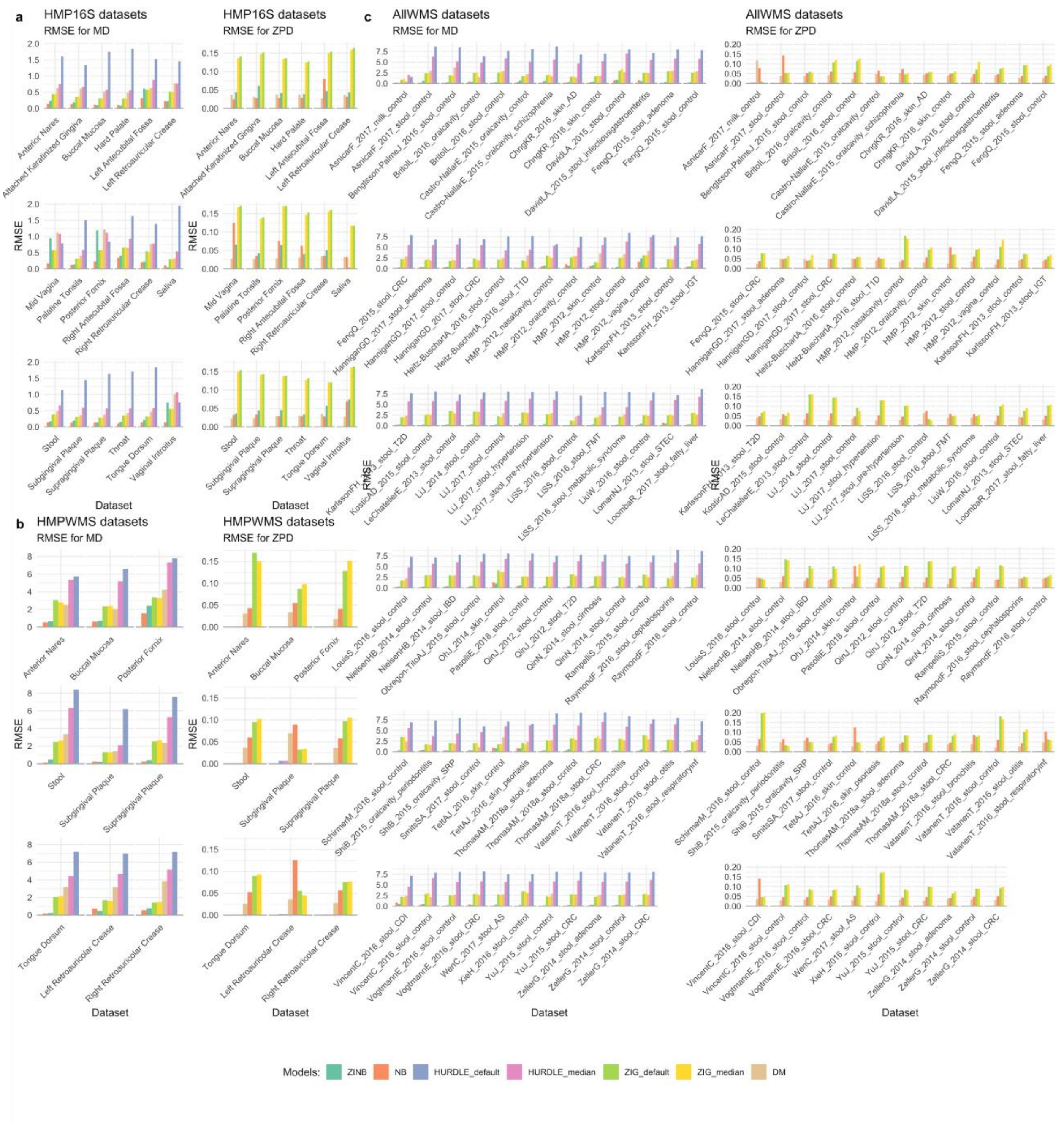

**Figure S2**: Mean differences (MD) and Zero Probability differences (ZPD) Root Mean Square Errors (RMSE) for HMP 16S datasets (a), HMP WMS datasets (b) and all the other WMS datasets (c).

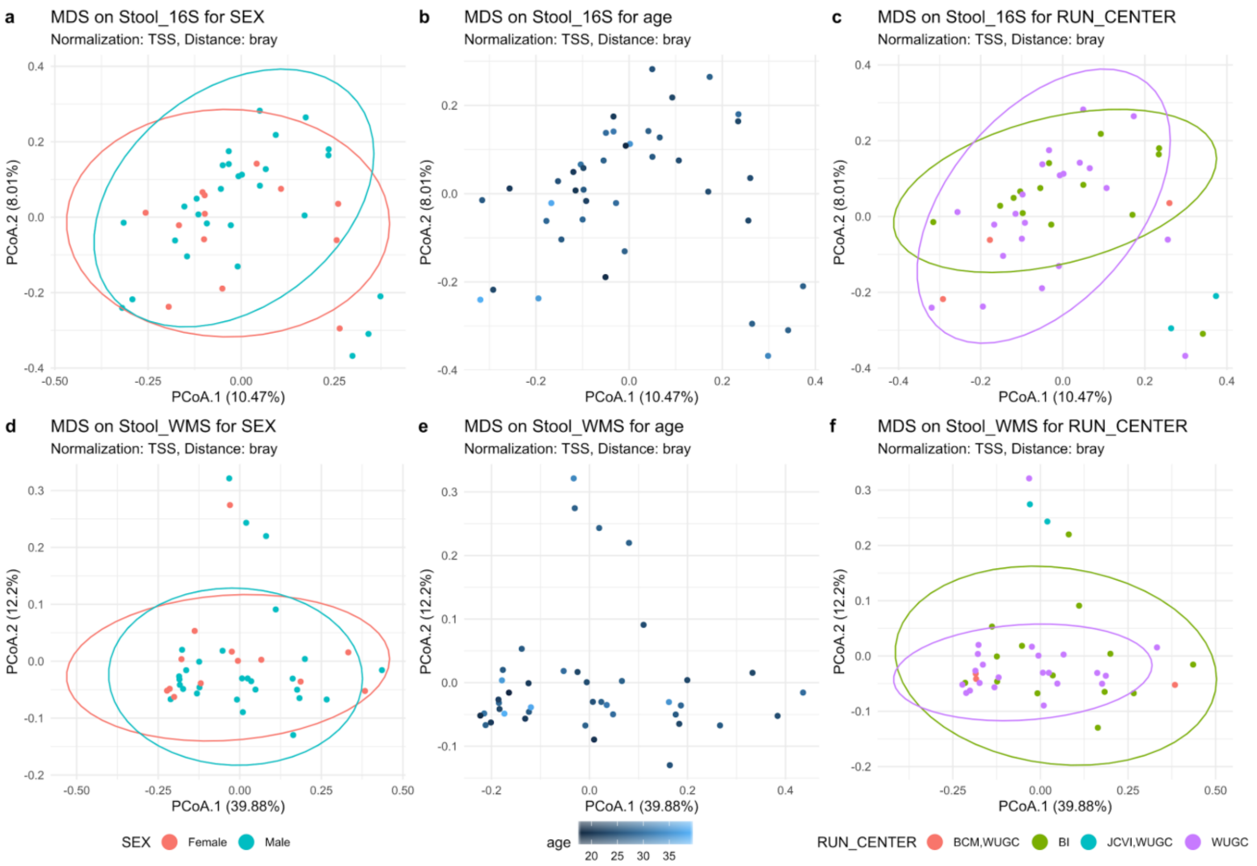

**Supplementary Figure S3:** MDS for Bray-Curtis distances on TSS normalized count for the HMP stool samples with the same RSID between 16S and WMS. **a.** 16S stool samples colored by SEX. **b.** 16S stool samples colored by age. **c.** 16S stool samples colored by RUN_CENTER. **d.** WMS stool samples colored by SEX. **e.** WMS stool samples colored by age. **f.** WMS stool samples colored by RUN_CENTER.

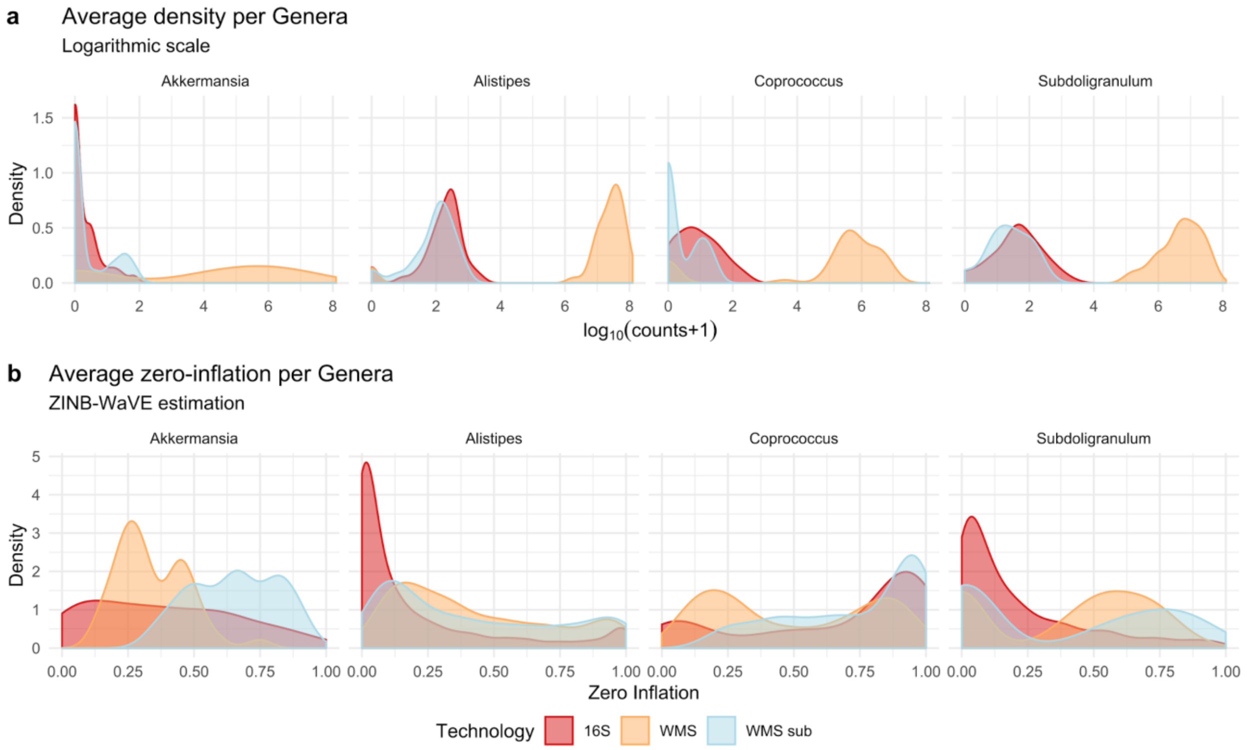

**Figure S4**: **a.** Average densities of log10(counts+1) for Akkermansia, Alistipes, Coprococcus and Subdoligranulum genera taken as representatives. **b.** Zero-inflation estimation through ZINB-WaVE package for the counts of Akkermansia, Alistipes, Coprococcus and Subdoligranulum genera taken as representatives. The difference between 16S and WMS approaches translates to different count data structures. This diversity is probably due to a mix of factors: sequencing depth, different taxonomic classification between technologies (entire metagenomic sequences versus clusters of similar amplicon sequences), bioinformatics methods for data preprocessing, etc. To investigate these differences, we studied the same RSID stool samples from HMP data sequenced in both 16S and WMS approaches. As the sequencing depth was the most visible difference between them (Supplementary Fig. S1c), we subsampled the WMS samples to make their library sizes equal to the 16Ss (rerefaction). Observing Supplementary Figure S4a, bimodality is clearly visible in both 16S and WMS average log count distributions for the Akkermansia and Alipstipes genera, while it is visible only for the WMS approach in Coprococcus and it seems not to be present for Subdoligranulum genera. After WMS subsampling, density profiles for 16S and WMS sub become very similar. However, bimodality in Akkermansia and Coprococcus density profiles are more visible in WMS sub rather than 16S.

The averaged zero-inflation probability estimates are reported in Figure S4b. After WMS subsampling, a difference between 16S and WMS sub profiles still exists, indicating the possible involvement of other factors explaining differences in the data structures.

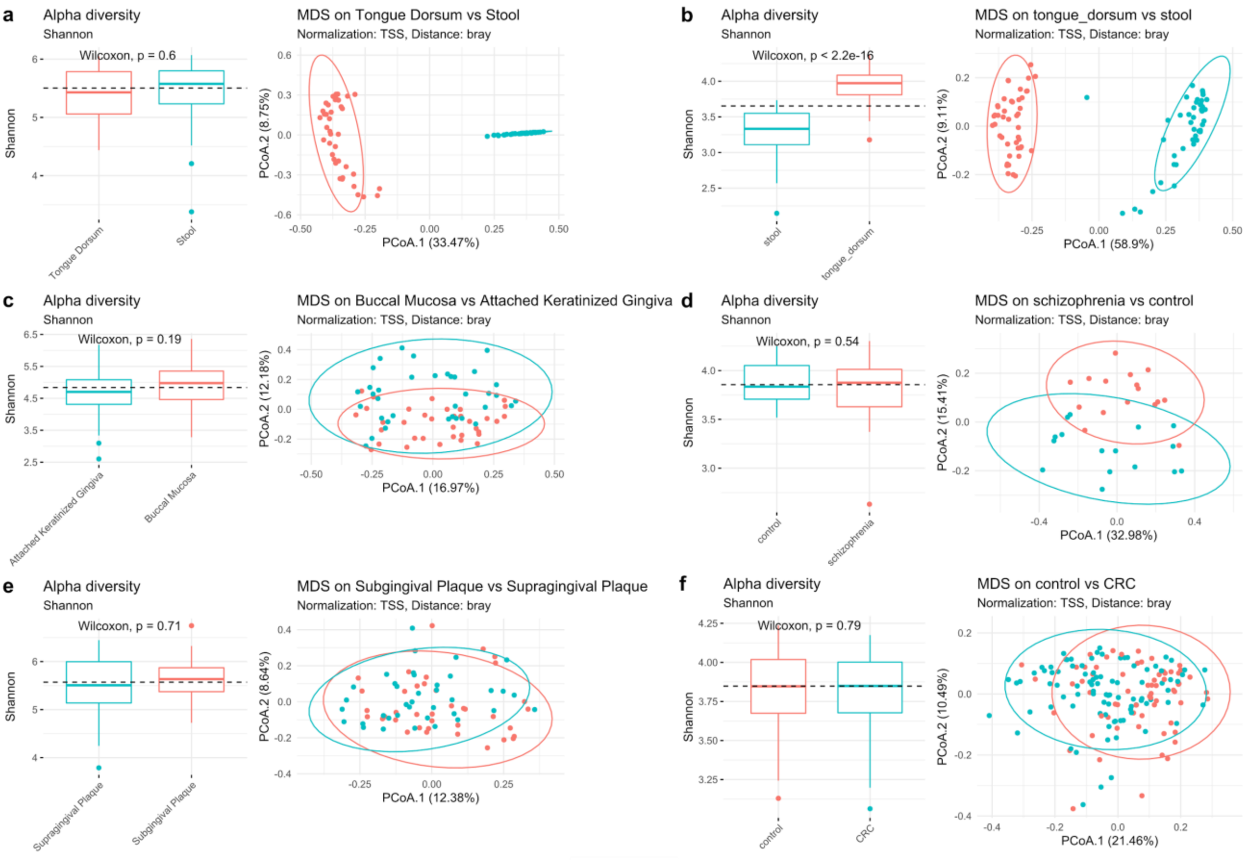

**Figure S5**: Alpha diversity (Shannon) and Beta diversity (Multidimensional Scaling ordination method and Bray-Curtis dissimilarity measure) of the original data used for concordance evaluation. **a.** High diversity, 16S, Stool vs Tongue Dorsum samples comparison. **b.** High diversity, WMS, Stool vs Tongue Dorsum samples comparison. **c.** Mid diversity, 16S, Buccal Mucosa vs Attached Keratinized Gingiva comparison. **d.** Mid diversity, WMS, Schizophrenia vs Healthy control saliva samples comparison. **e.** Low diversity, 16S, Supragingival vs Subgingival plaque samples comparison. **f.** Low diversity, WMS, Colon Rectal Cancer vs Healthy control stool samples comparison.

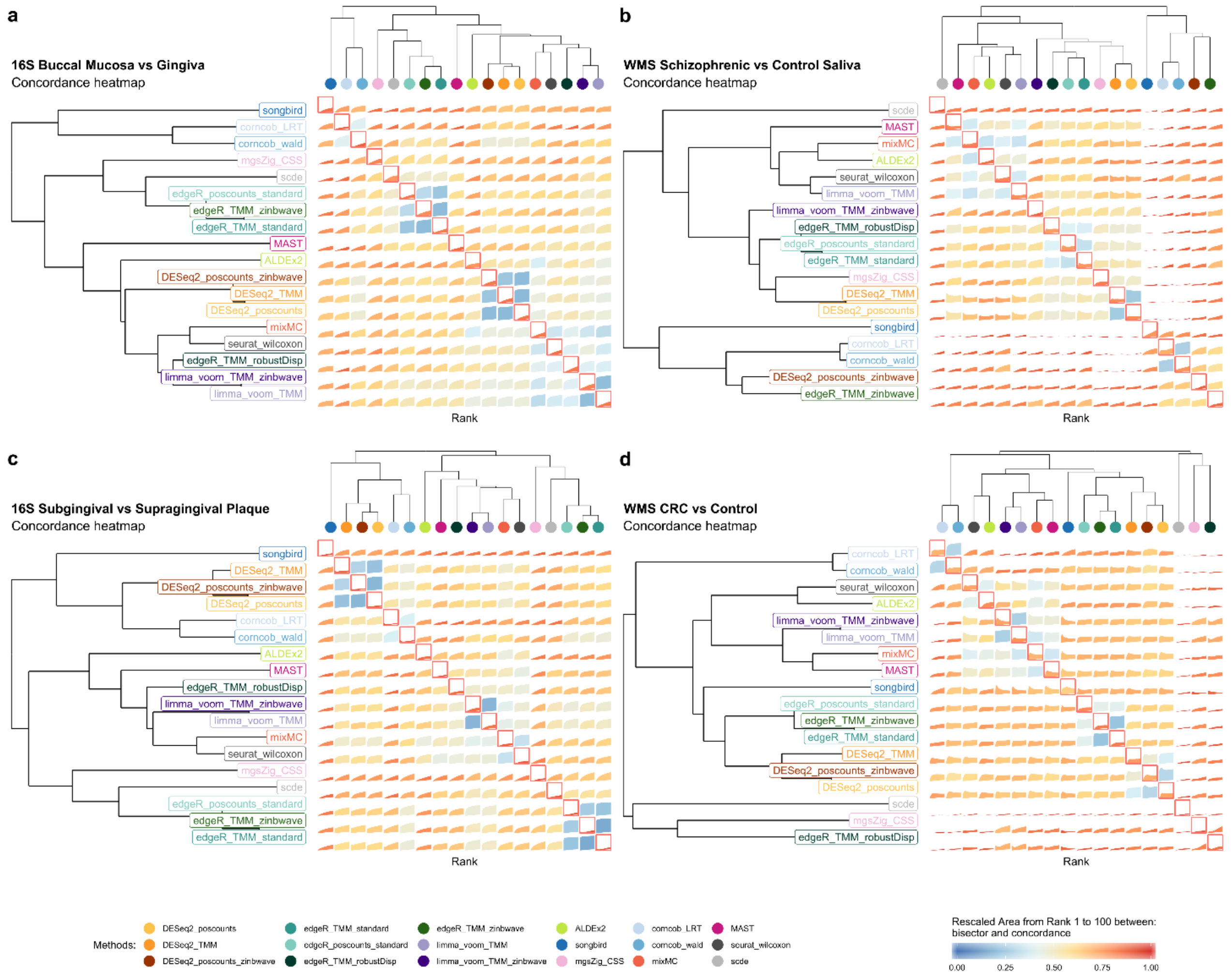

**Figure S6: a.** BMC and WMC (main diagonal) averaged values from rank 1 to 100 for DA methods evaluated in replicated 16S Buccal Mucosa vs Attached Keratinized Gingiva comparisons. **b.** BMC and WMC (main diagonal) averaged values from rank 1 to 100 for DA methods evaluated in replicated WMS Schizophrenia vs Healthy control stool samples comparison. **c.** BMC and WMC (main diagonal) averaged values from rank 1 to 100 for DA methods evaluated in replicated 16S Supragingival vs Subgingival plaque samples comparison. **d.** BMC and WMC (main diagonal) averaged values from rank 1 to 100 for DA methods evaluated in replicated WMS Colon Rectal Cancer vs Healthy control stool samples comparison.

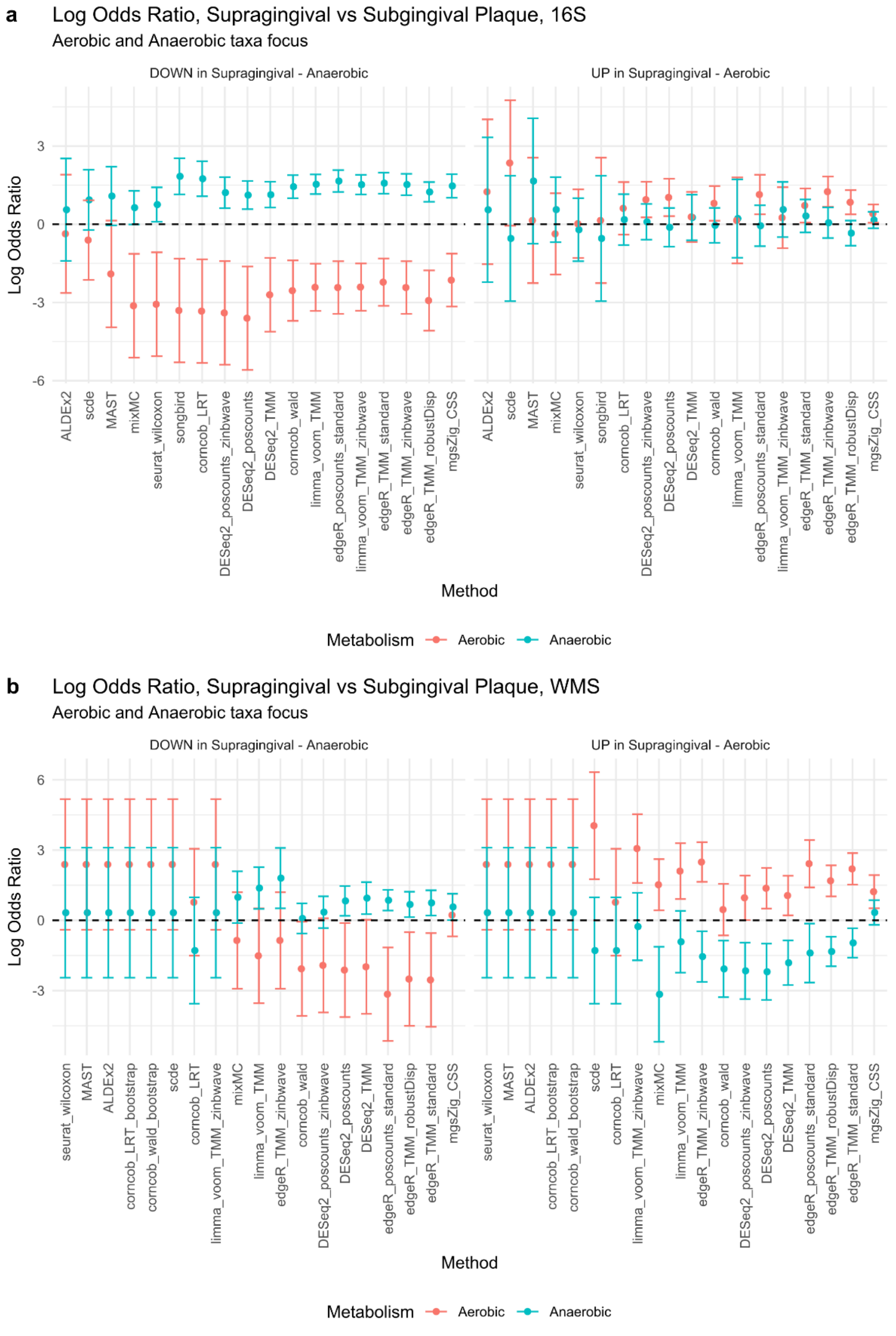

**Supplementary Figure S7: a.** Log Odds Ratio for Aerobic and Anaerobic taxa, over-abundant and under-abundant in Supragingival Plaque respectively, in HMP 16S samples. **b.** Log Odds Ratio for Aerobic and Anaerobic taxa, over-abundant and under-abundant in Supragingival Plaque respectively, in HMP WMS samples.

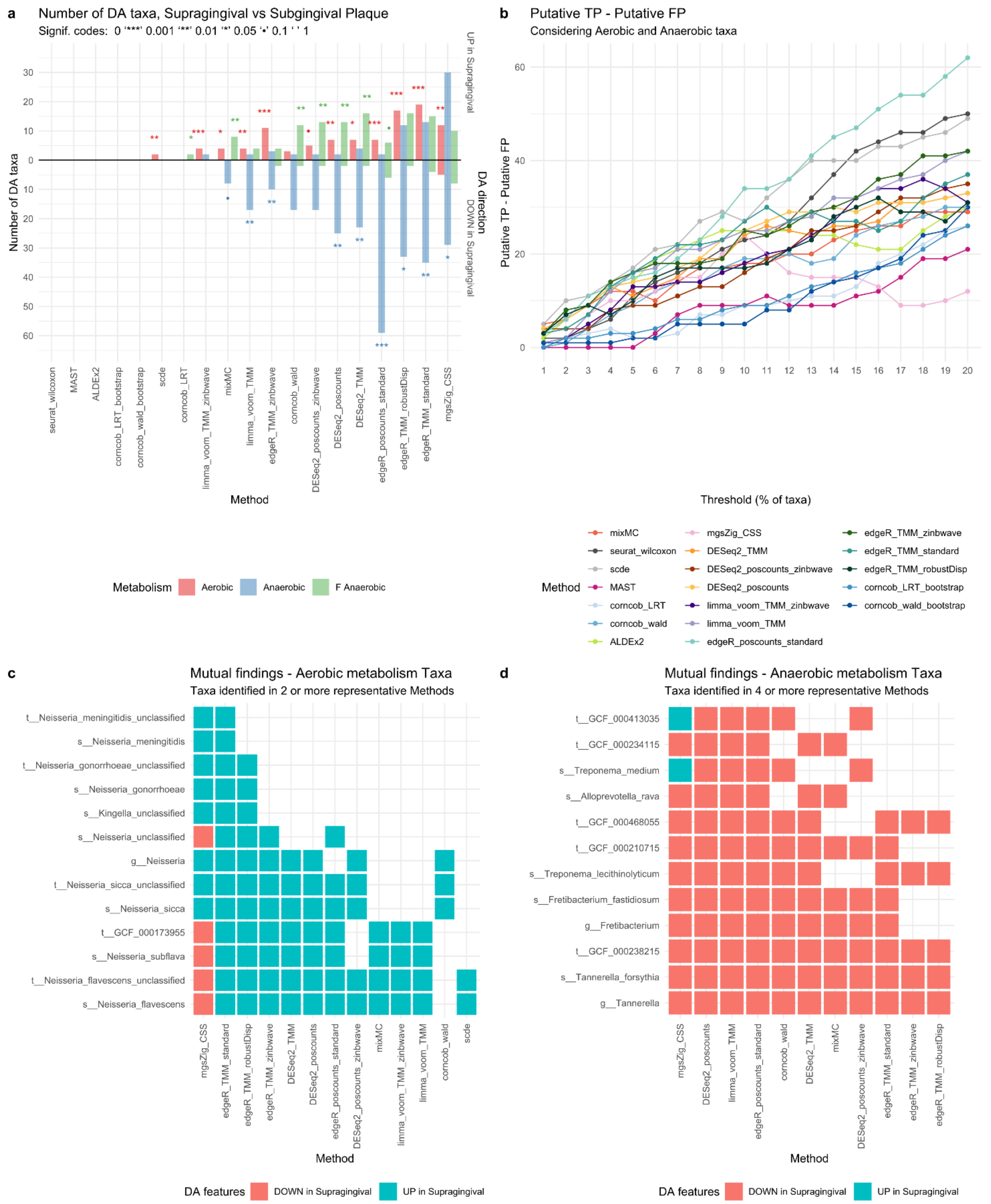

**Supplementary Figure S8:** 5vs5 Supragingival vs Subgingival Plaque HMP WMS samples **a.** Barplot for the enrichment tests performed on the DA taxa found by each method using an Adjusted p-value of 0.1 as threshold for significance (top 10% ranked taxa for songbird). Each bar represents the number of findings, UP in Supragingival or DOWN in Supragingival Plaque compared to Subgingival Plaque, regarding Aerobic, Anaerobic and Facultative Anaerobic taxa metabolism. A Fisher exact test is performed to establish the enrichment significance which is represented with signif. codes. **b.** Difference between putative True Positives (TP) and putative False Positives (FP) is drawn for several significance thresholds. Each threshold represents the top percent ranked taxa, using the ordered raw p-value lists as reference (loading values for mixMC and differentials for songbird). **c.** Aerobic metabolism taxa mutually found by 2 or more methods from the subset of the representative methods. **d.** Anaerobic metabolism taxa mutually found by 4 or more methods from the subset of the representative methods.

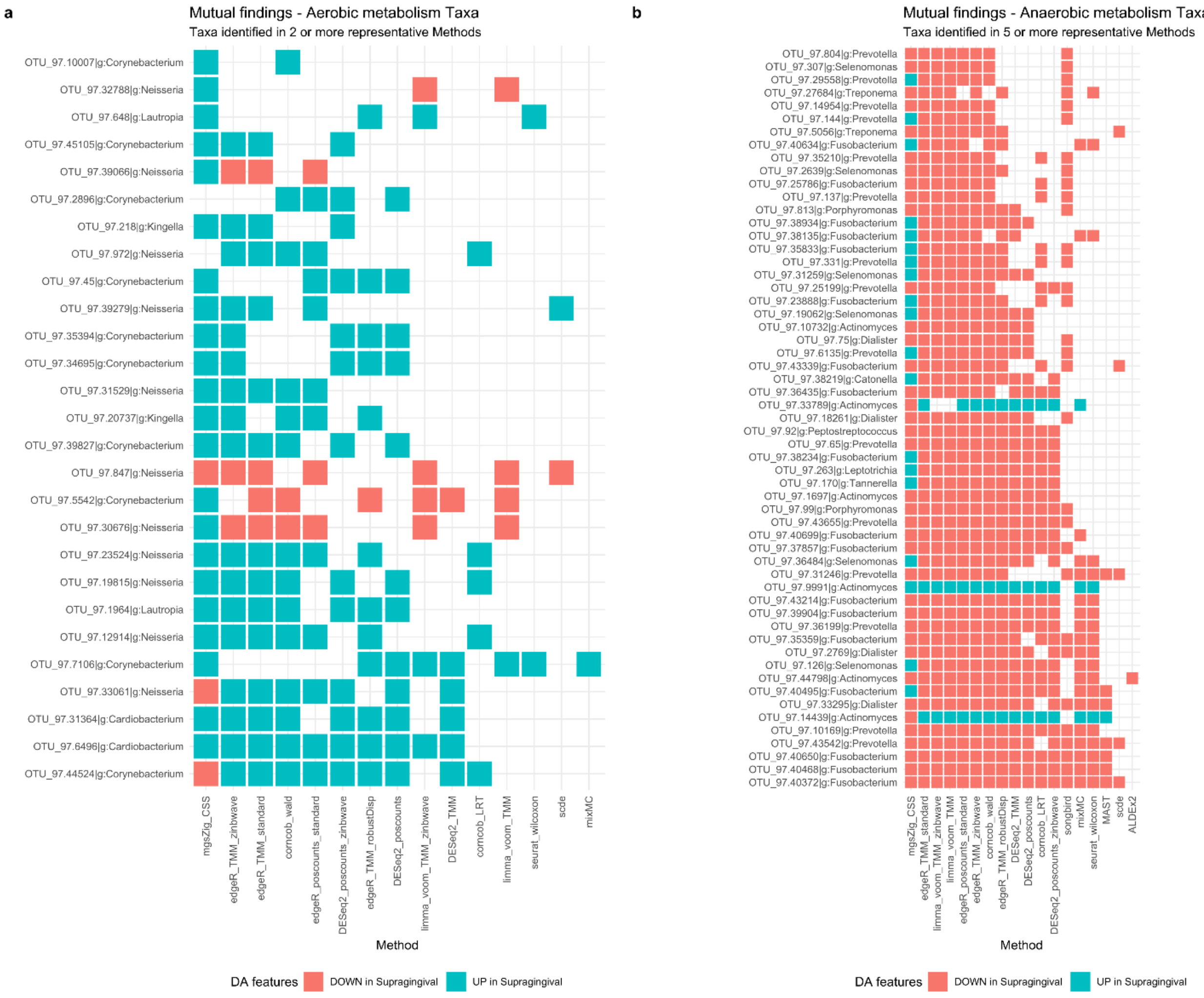

**Supplementary Figure S9:** 38vs38 Supragingival vs Subgingival Plaque HMP 16S samples **a.** Aerobic metabolism taxa mutually found by 2 or more methods from the subset of the representative methods. **d.** Anaerobic metabolism taxa mutually found by 5 or more methods from the subset of the representative methods.
