## Additional file 2 for "Assessment of statistical methods from single cell, bulk RNA-seq and metagenomics applied to microbiome data"

### Parametric simulations

While the results of the experimental datasets are best for assessing model fit and consistency of discoveries, the lack of ground truth makes it impossible to assess the validity of discoveries. For this reason, in addition to the enrichment analysis, which is restricted to one dataset, we turned to simulated data to explore the properties of the methods in more detail. Here, we specifically asked whether it was important to model zero inflation and, given the results of our GOF analysis (Fig. 2), we only used the NB and ZINB distributions to simulate the data. Briefly, for each distribution we simulated 7200 and 19200 scenarios respectively, mimicking both 16S and WMS data, and varying the sample size, the proportion of DA features, and the amount of the effect. We also varied the proportion of zeros and whether there was an interaction between the number of zeros and DA (sparsity effect, see Methods for details; Additional file 2: Supplementary Table S4).

Supplementary Figure S11 summarizes the performance of all methods according to all the different variables involved in the simulation procedure. To condense all the results into a single figure, we ranked the methods summarizing the pAUROC performance independently for each simulation parameter. Importantly, this summary ignored many interaction effects (e.g., whether the fold effect influences the results differently depending on the technology). Albeit simplified, this summary is nonetheless useful to get an overview of each method's performance. We used the partial Area Under the Receiver Operating Characteristic Curve (pAUROC) between 0 and 0.1 of False Positive Rate (FPR) values as an indicator of the method performances, since it only considers the range of FPR values that are important in practice and measures the ability of methods to correctly detect true differential abundant features. A method-specific pattern is clearly visible, indicating the robustness and coherence of each method across different simulation scenarios (Additional file 2: Supplementary Figure S11).

Briefly, edgeR with TMM normalization (with and without zinbwave weights) and DESeq2 with poscount normalization and zinbwave weights were the overall best methods (Fig. 6a). The other

DESeq2-based methods were close second together with corncob (with Wald test). Unsurprisingly, the parametric distribution that generated the data had great influence on the method performances. Indeed, ZINB generated datasets showed lower mean values for all methods because of the increase in sparsity. All methods' performances increased as the sample size and/or the fold effect increased (Supplementary Figure S11). Focusing on the amount of zero counts, we observed that the mean performance increased when the sparsity effect increased from 0.05 to 0.15, not only for edgeR and DESeq2 based method, but for limma-voom, ALDEx2 and Wilcoxon (Supplementary Figure S11).

Confirming our real data results, metagenomeSeq, scde, and edgeR with robust dispersion estimation performed poorly. On the other hand, MAST, which showed mixed results in real data, did not behave in simulations, partly because of the misspecified model with respect to the data generating distribution.

The most time-consuming methods were scde, ALDEx2, MAST and zinbwave. In simulated datasets with 40 samples per condition and less than 1000 features we observed an average elapsed time of around 5 minutes for scde; ALDEx2 took an average of 30 seconds, MAST took an average of 14 seconds, DESeq2 less than 8 seconds, and edgeR and limma-voom took less than 1 second, although observational weights estimation took an average of 18 seconds (Additional file 2: Supplementary Table S5, Supplementary Fig. S7).

In conclusion, all methods displayed drastically lowered performance for datasets with increased sparsity.  A way to decrease sparsity is to filter rare features or to impute the zero counts: in our simulations, for the sake of simplicity, we decided to keep the features with more than ten reads in at least two samples. Other works [13] or analysis pipelines [51], suggest several different filters that may have an impact on the results. Beyond the filtering choice, for a method that does not treat zero counts in any special way, it is easier to detect differentially abundant features when zero rates are clearly different between experimental groups. Indeed, the change in the proportion of zeros will affect the mean and hence the DA statistic. The same situation can be tricky for methods that downweight the contribution of zero counts (e.g., zinbwave). On the other hand, when zero counts are equally present in both groups, downweighting them is favourable (Additional file 2: Supplementary Figure S11). These two opposite situations are similar to the difference between treating the zero counts as “biological” or “technical” and more research is needed to understand whether the zero-inflated model can help classify the two.

In simulated datasets with 40 samples per condition and less than 1000 features we observed an average elapsed time of around 4 minutes for scde; ALDEx2 took an average of 25 seconds, corncob took an average of 23 seconds, MAST took an average of 5 seconds, DESeq2 less than 5 seconds, and edgeR and limma-voom took less than 1 second, although observational weights estimation took an average of 18 seconds (Additional file 2: Supplementary Table S5 and Supplementary Fig. S10).

**Supplementary Table S4:** Simulation framework variables.

| Variable | Description | Type | Values |
|---|---|---|---|
| **dataset** | Dataset names | categorical | Stool_16S, Stool_WMS, TongueDorsum_16S, TongueDorsum_WMS BritoIL_Stool, BritoIL_Oral |
| **distribution** | Generating distribution | dichotomic | NB, ZINB |
| **simulation** | Number of the simulation | numerical | 1 to 50 |
| **sampleSize** | Number of samples for each condition | numerical | 10, 20, 40 |
| **TPR** | Proportion of differential abundant features to generate | numerical | 0.1, 0.5 |
| **foldEffect** | Differential abundance multiplicative factor | numerical | 2, 5 |
| **compensation** | Balancing non differential abundant features after applying foldEffect | dichotomic | yes, no |
| **sparsityEffect** | Additive factor which increases or lowers sparsity for down-regulated and up-regulated features respectively. Only for ZINB generated data. | numerical | 0, 0.05, 0.15 |

**Supplementary Table S5**: Average computational times and standard deviations in seconds for each method, grouped by the sample size and the parametric distribution which generates the data. Boxplots in Supplementary Figure S7.

| **method\sampleSize** | **NB** | | | **ZINB** | | |
|---|---|---|---|---|---|---|
| | **10** | **20** | **40** | **10** | **20** | **40** |
| **limma_voom_TMM** | 0.06 (0.02) | 0.08 (0.03) | 0.1 (0.04) | 0.04 (0.01) | 0.06 (0.01) | 0.08 (0.01) |
| **edgeR_TMM_standard** | 0.22 (0.07) | 0.35 (0.11) | 0.51 (0.09) | 0.19 (0.04) | 0.3 (0.06) | 0.49 (0.06) |
| **edgeR_poscounts_standard** | 0.36 (0.11) | 0.5 (0.15) | 0.65 (0.13) | 0.31 (0.07) | 0.43 (0.1) | 0.62 (0.07) |
| **mgsZig_CSS** | 2.18 (0.59) | 2.2 (0.57) | 1.92 (0.31) | 1.9 (0.46) | 2.04 (0.44) | 2.11 (0.6) |
| **seurat_wilcoxon** | 3.65 (0.78) | 3.69 (0.76) | 3.3 (0.5) | 3.43 (0.86) | 3.45 (0.88) | 3.32 (0.42) |
| **DESeq2_TMM** | 1.72 (0.48) | 2.23 (0.54) | 3.06 (0.63) | 2.93 (1.52) | 4.27 (2.1) | 6.07 (2.05) |
| **DESeq2_poscounts** | 1.9 (0.53) | 2.37 (0.6) | 3.25 (2.74) | 3.1 (1.57) | 4.49 (2.22) | 6.28 (2.11) |
| **MAST** | 8.08 (4.6) | 7.89 (4.31) | 5.19 (1.69) | 4.19 (3.14) | 4.76 (2.81) | 4.33 (0.97) |
| **edgeR_TMM_robustDisp** | 3.51 (0.91) | 5.21 (1.24) | 7.33 (3.06) | 3.68 (1.9) | 6.02 (1.37) | 9.61 (0.98) |
| **limma_voom_TMM_zinbwave** | 5.08 (1.74) | 14.46 (13.19) | 16.27 (11.7) | 7.42 (4.12) | 12.84 (5.65) | 24.43 (10.85) |
| **edgeR_TMM_zinbwave** | 5.22 (1.76) | 14.71 (13.22) | 16.68 (11.75) | 7.55 (4.13) | 13.07 (5.66) | 24.84 (10.86) |
| **corncob_LRT** | 19.3 (4.96) | 22.46 (4.51) | 24.18 (2.62) | 12.53 (3.83) | 17.08 (3.32) | 21.98 (2.07) |
| **ALDEx2** | 16.58 (3.94) | 20.91 (4.89) | 25.21 (3.91) | 12.5 (3.17) | 17.01 (3.37) | 23.97 (2.51) |
| **corncob_wald** | 19.71 (5.06) | 23.16 (4.72) | 24.84 (2.94) | 12.86 (3.88) | 17.58 (3.5) | 22.59 (2.13) |
| **DESeq2_poscounts_zinbwave** | 7.06 (2.05) | 17.05 (13.46) | 20.1 (12.17) | 12.15 (6.87) | 19.92 (9.63) | 34.07 (11.97) |
| **scde** | 61.07 (22.02) | 128.39 (57.28) | 226.17 (102.25) | 46.7 (10.68) | 114.7 (21.66) | 253.12 (43.85) |

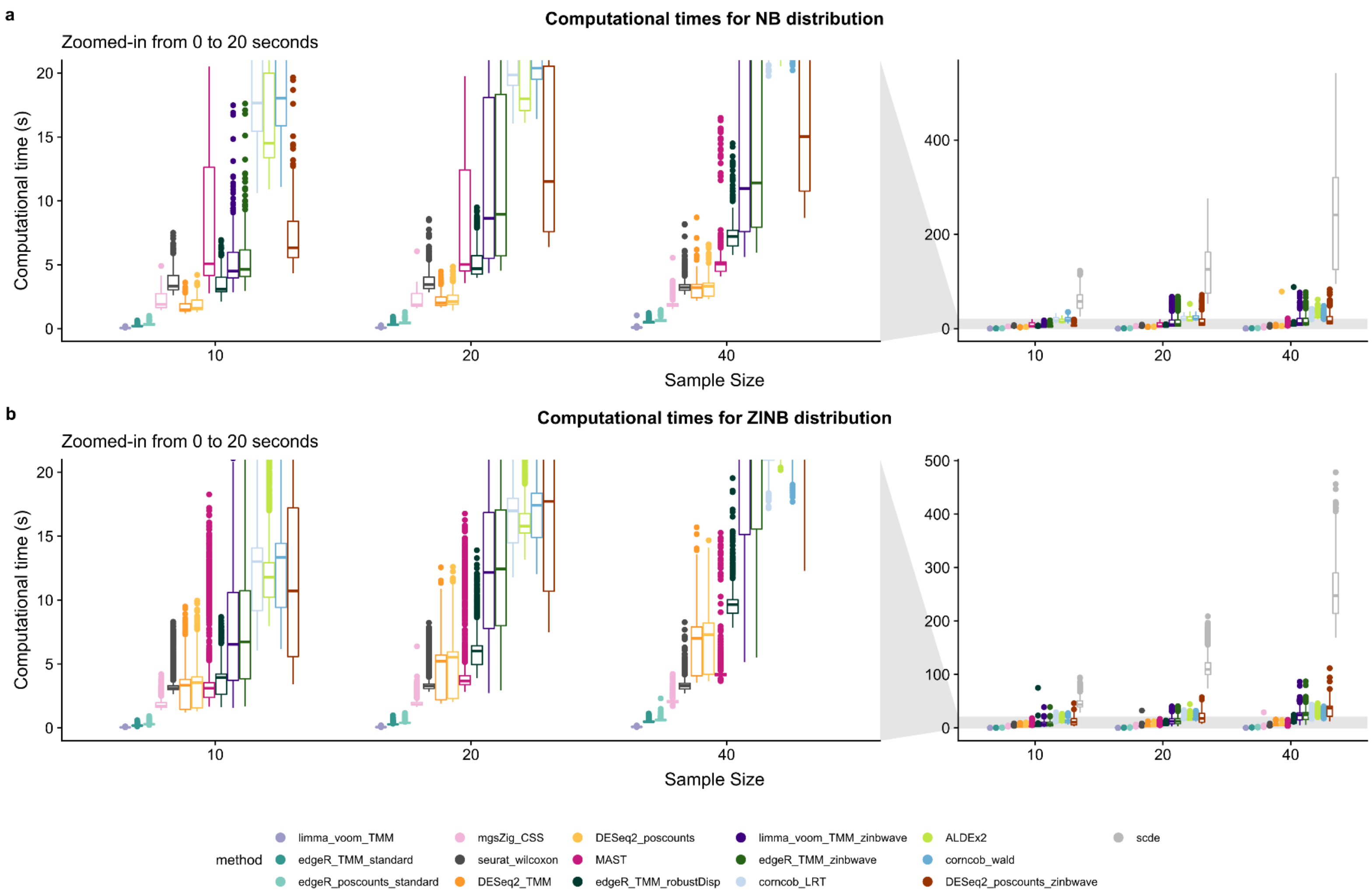


**Figure S10**: **a**. Average computational time boxplots (in seconds) for each method, grouped by sample size when data are NB generated. On the left side a zoom from 0 to 20 seconds is reported. **b.** Average computational time boxplots (in seconds) for each method, grouped by sample size when data are ZINB generated. On the left side a zoom from 0 to 20 seconds is reported.

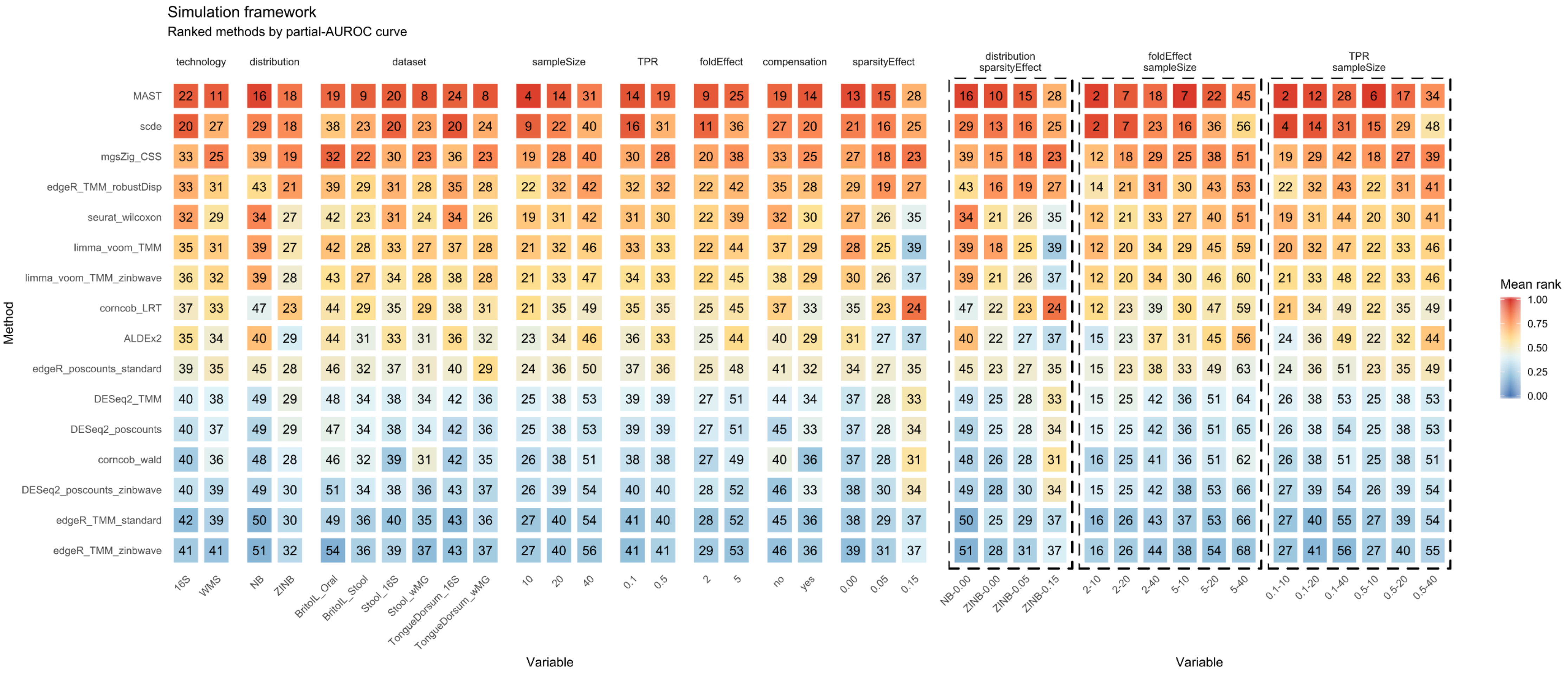


**Supplementary Figure S11:** Comprehensive univariate and bivariate (dotted line boxes) evaluation of the pAUROC curves from FPR 0 to 0.1 (For a full multivariate method comparison see next pages: Additional file 2). Average normalized ranks range from 0 to 1, lower values correspond to better performances. The value inside each tile refers to the average rescaled partial AUROC value multiplied per 100, on which ranks are computed. Higher pAUROC values correspond to better performance.

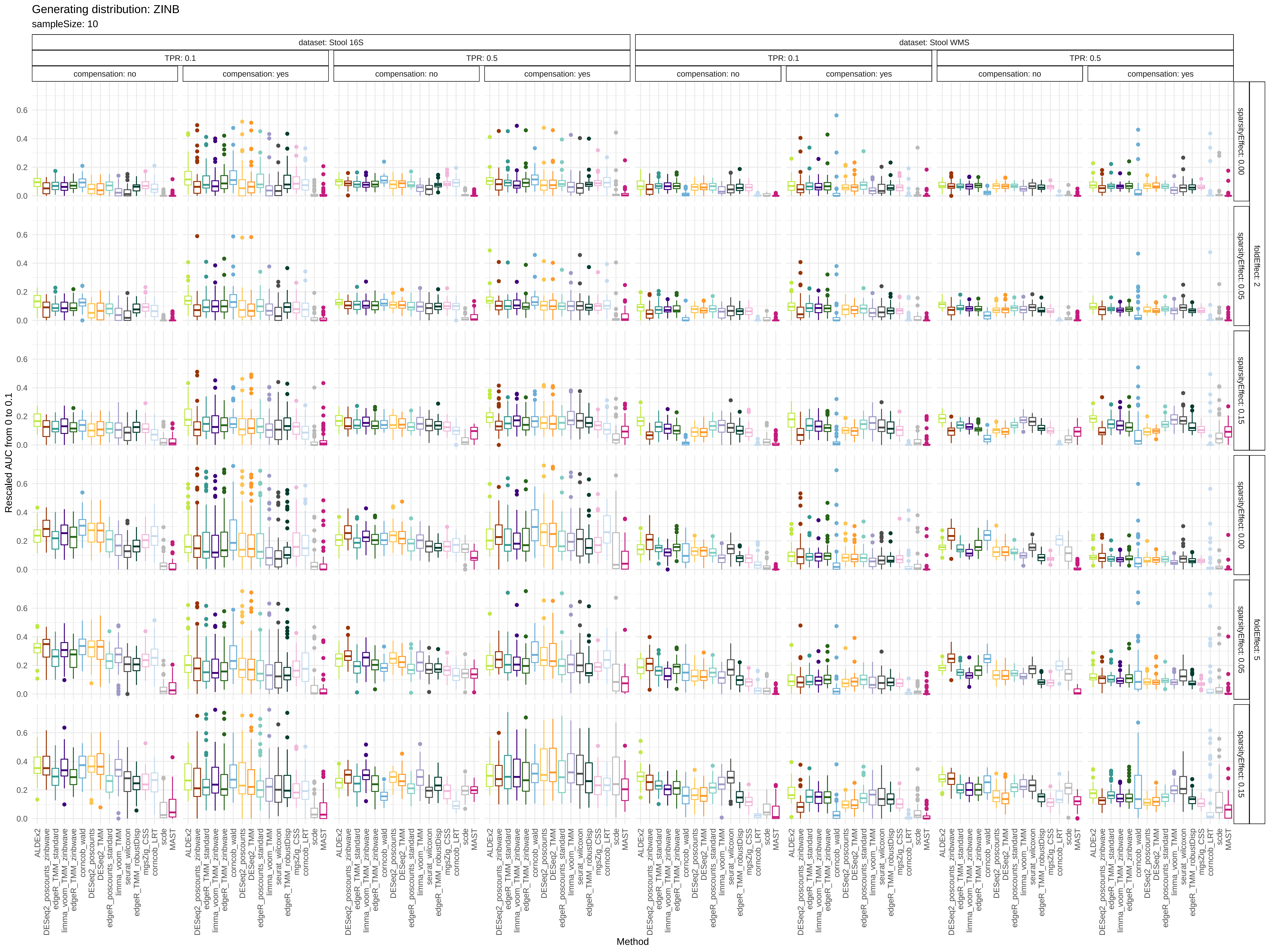

Generating distribution: ZINB
sampleSize: 10
dataset: Stool 16S
dataset: Stool WMS
TPR: 0.1
TPR: 0.5
compensation: no
compensation: yes
sparsityEffect: 0.00
sparsityEffect: 0.05
sparsityEffect: 0.15
foldEffect: 2
foldEffect: 5
Rescaled AUC from 0 to 0.1
0.0
0.2
0.4
0.6
ALDEx2
DESeq2_poscounts_zinbwave
edgeR_TMM_standard
limma_voom_TMM_zinbwave
edgeR_TMM_zinbwave
corncob_wald
DESeq2_poscounts
DESeq2_TMM
edgeR_poscounts_standard
limma_voom_TMM
seurat_wilcoxon
edgeR_TMM_robustDisp
mgsZig_CSS
corncob_LRT
scde
MAST
Method

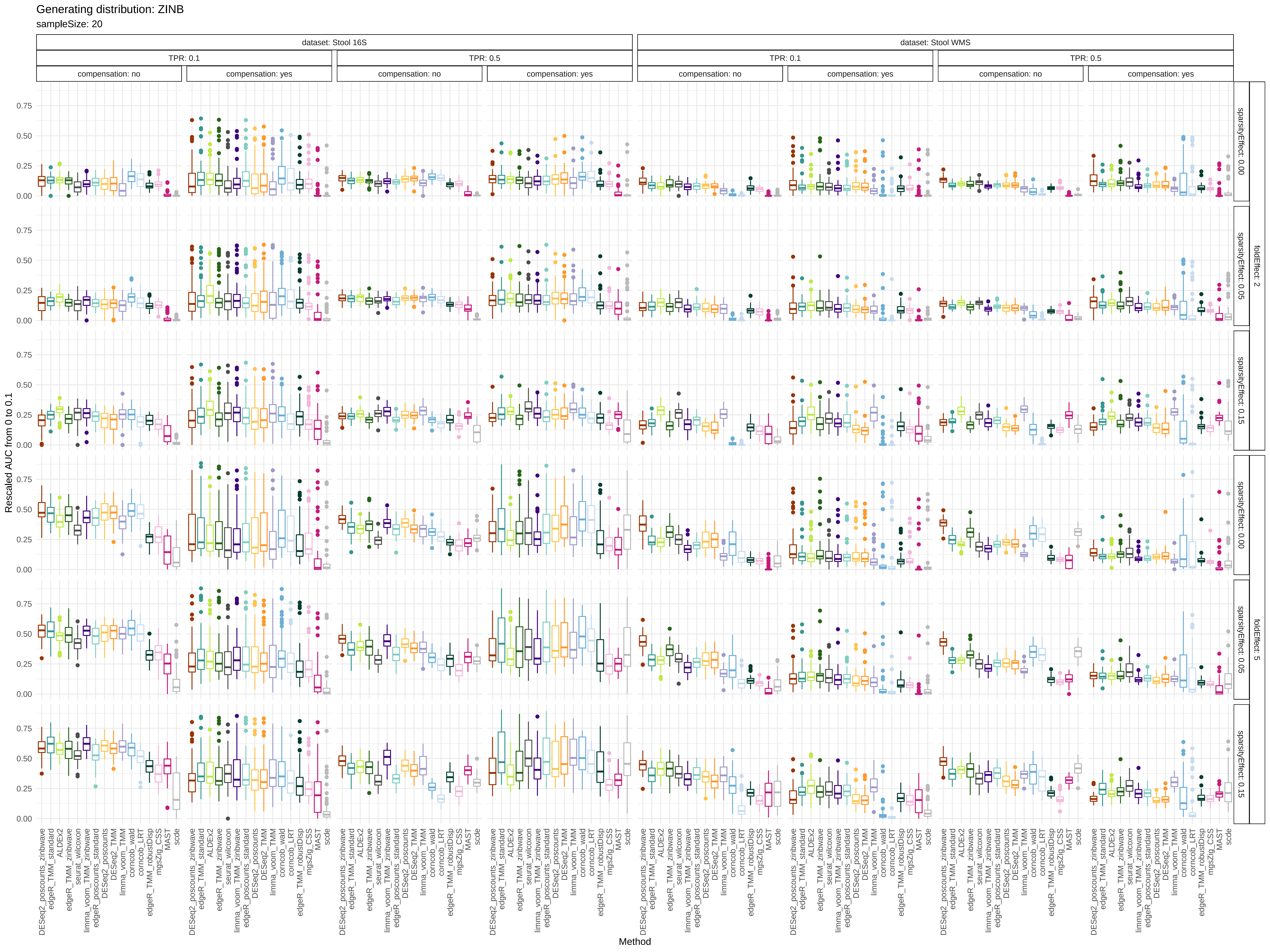

Generating distribution: ZINB
sampleSize: 20
dataset: Stool 16S
dataset: Stool WMS
TPR: 0.1
TPR: 0.5
compensation: no
compensation: yes
foldEffect: 2
foldEffect: 5
sparsityEffect: 0.00
sparsityEffect: 0.05
sparsityEffect: 0.15
Rescaled AUC from 0 to 0.1
0.00
0.25
0.50
0.75
DESeq2_poscounts_zinbwave
edgeR_TMM_standard
ALDEx2
edgeR_TMM_zinbwave
seurat_wilcoxon
limma_voom_TMM_zinbwave
edgeR_poscounts_standard
DESeq2_poscounts
DESeq2_TMM
limma_voom_TMM
corncob_wald
corncob_LRT
edgeR_TMM_robustDisp
mgsZig_CSS
MAST
scde
Method

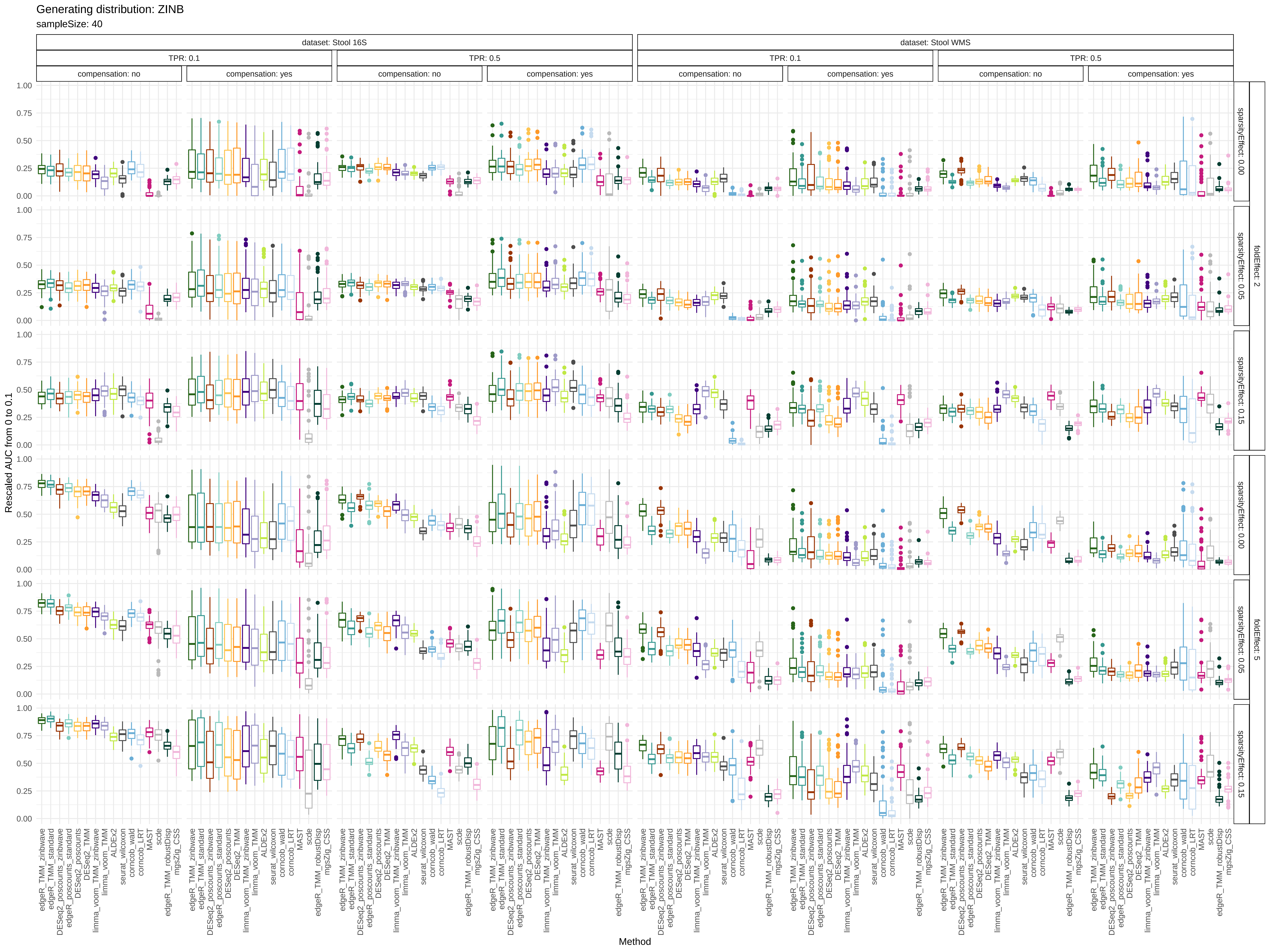

Generating distribution: ZINB
sampleSize: 40
dataset: Stool 16S
dataset: Stool WMS
TPR: 0.1
TPR: 0.5
compensation: no
compensation: yes
foldEffect: 2
foldEffect: 5
sparsityEffect: 0.00
sparsityEffect: 0.05
sparsityEffect: 0.15
Rescaled AUC from 0 to 0.1
Method
edgeR_TMM_zinbwave
edgeR_TMM_standard
DESeq2_poscounts_zinbwave
edgeR_poscounts_standard
DESeq2_poscounts
DESeq2_TMM
limma_voom_TMM_zinbwave
limma_voom_TMM
ALDEx2
seurat_wilcoxon
corncob_wald
corncob_LRT
MAST
scde
edgeR_TMM_robustDisp
mgsZig_CSS

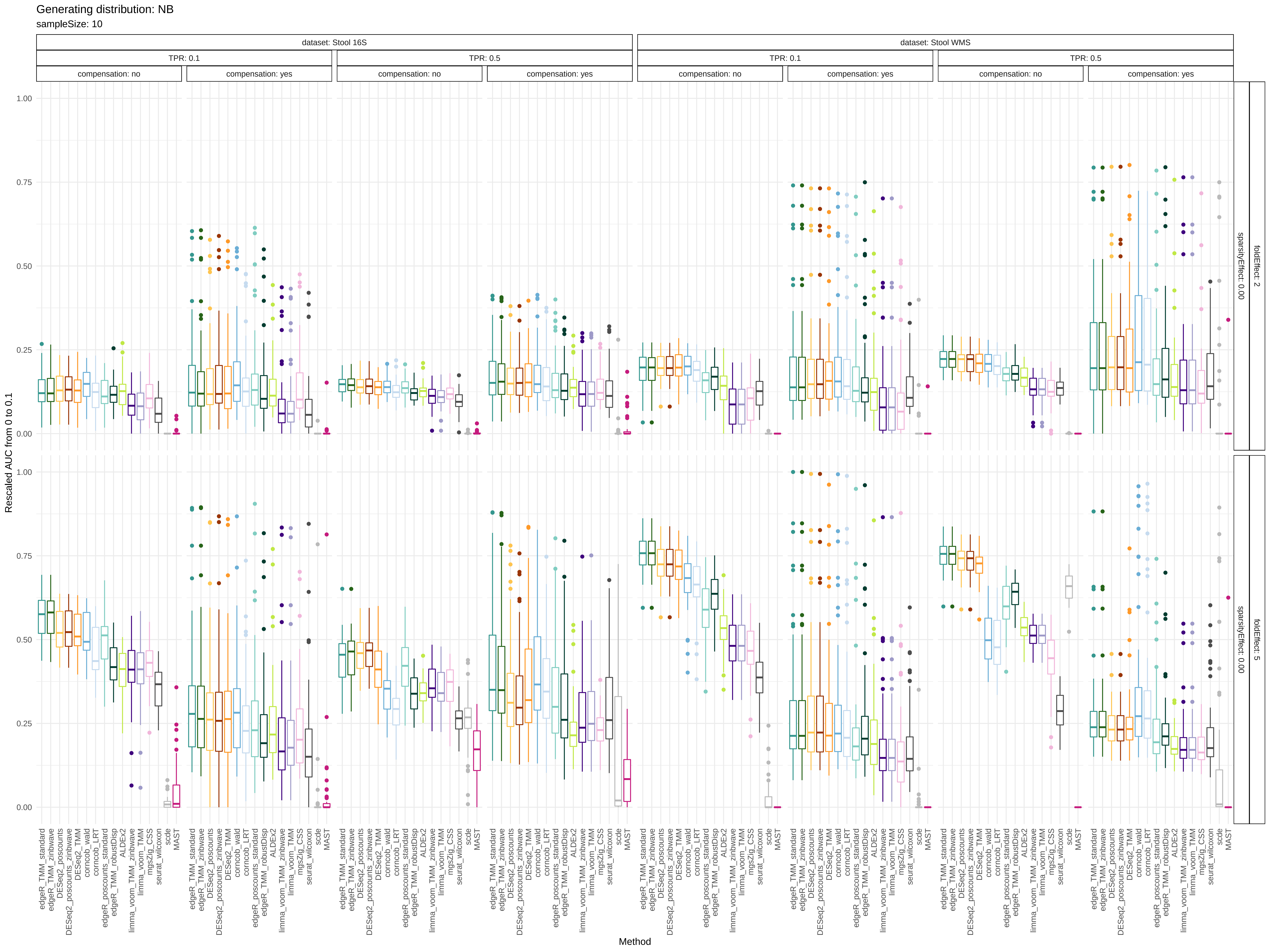
Generating distribution: NB
sampleSize: 10
dataset: Stool 16S
dataset: Stool WMS
TPR: 0.1
TPR: 0.5
compensation: no
compensation: yes
foldEffect: 2
sparsityEffect: 0.00
foldEffect: 5
Rescaled AUC from 0 to 0.1
1.00
0.75
0.50
0.25
0.00
edgeR_TMM_standard
edgeR_TMM_zinbwave
DESeq2_poscounts
DESeq2_poscounts_zinbwave
DESeq2_TMM
corncob_wald
corncob_LRT
edgeR_poscounts_standard
edgeR_TMM_robustDisp
ALDEx2
limma_voom_TMM_zinbwave
limma_voom_TMM
mgsZig_CSS
seurat_wilcoxon
scde
MAST
Method

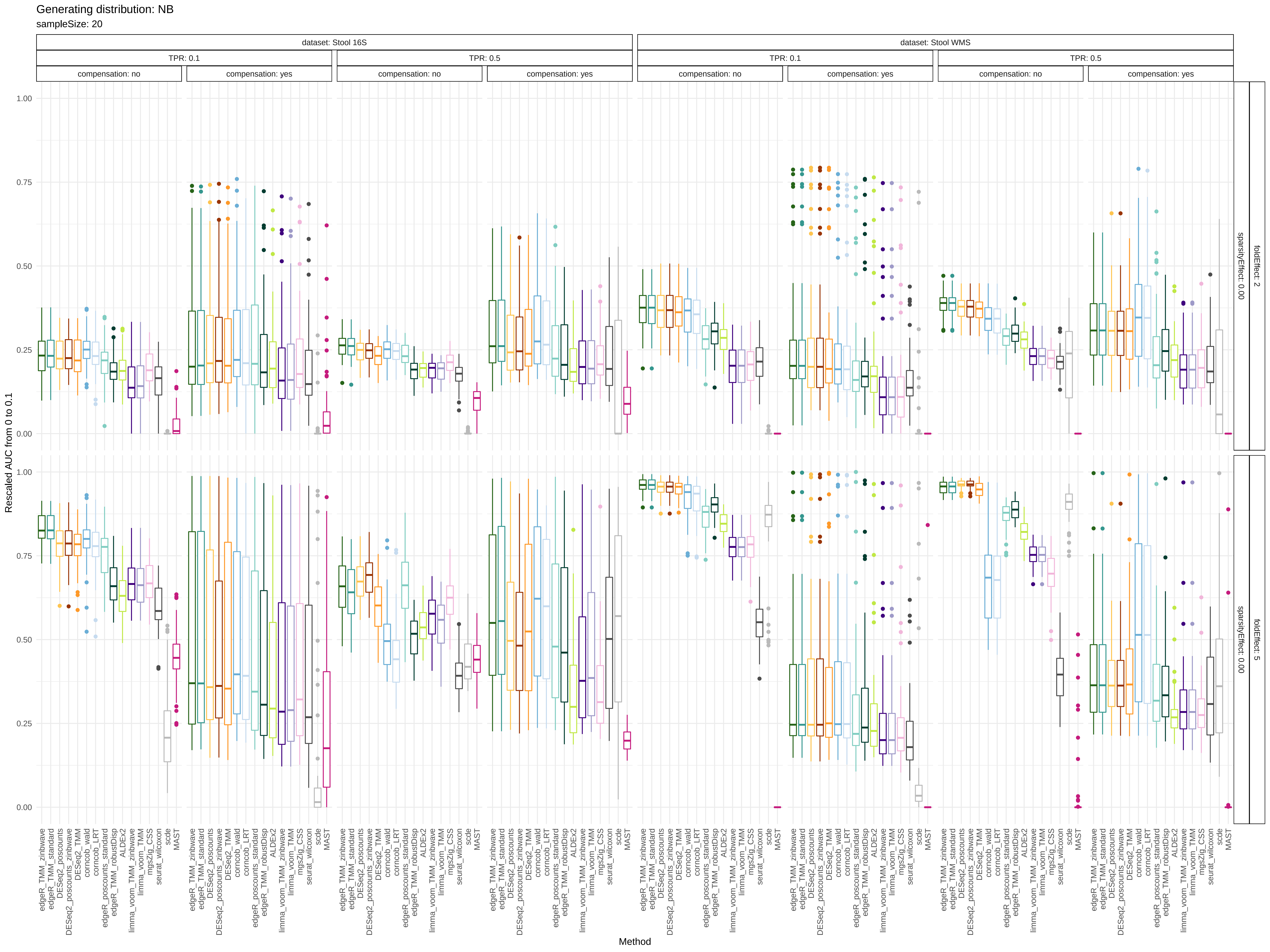
Generating distribution: NB
sampleSize: 20
dataset: Stool 16S
dataset: Stool WMS
TPR: 0.1
TPR: 0.5
compensation: no
compensation: yes
foldEffect: 2
foldEffect: 5
sparsityEffect: 0.00
Rescaled AUC from 0 to 0.1
0.00
0.25
0.50
0.75
1.00
Method
edgeR_TMM_zinbwave
edgeR_TMM_standard
DESeq2_poscounts
DESeq2_poscounts_zinbwave
DESeq2_TMM
corncob_wald
corncob_LRT
edgeR_poscounts_standard
edgeR_TMM_robustDisp
ALDEx2
limma_voom_TMM_zinbwave
limma_voom_TMM
mgsZig_CSS
seurat_wilcoxon
scde
MAST

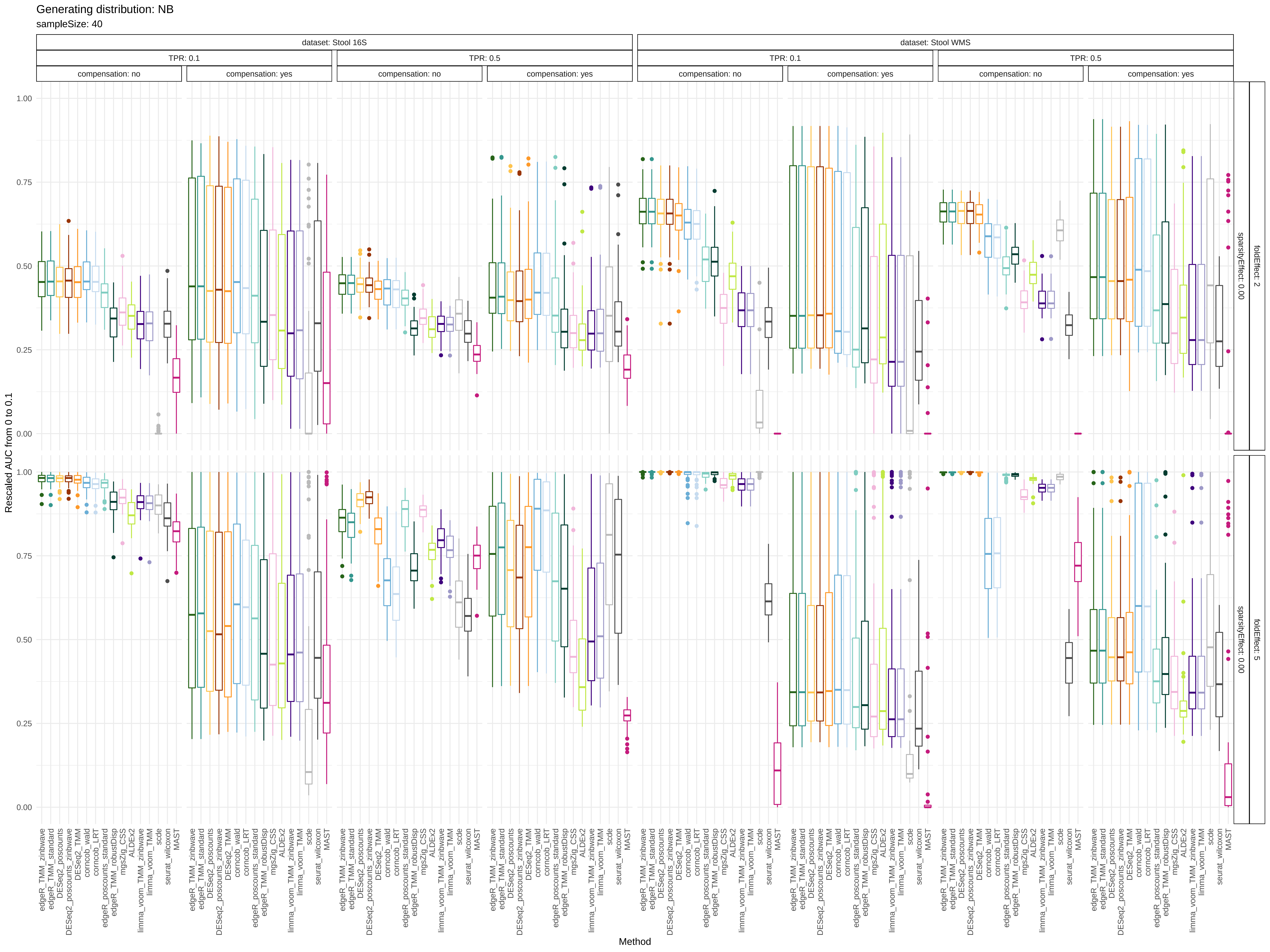
Generating distribution: NB
sampleSize: 40
dataset: Stool 16S
dataset: Stool WMS
TPR: 0.1
TPR: 0.5
compensation: no
compensation: yes
foldEffect: 2
foldEffect: 5
sparsityEffect: 0.00
Rescaled AUC from 0 to 0.1
Method
0.00
0.25
0.50
0.75
1.00
edgeR_TMM_zinbwave
edgeR_TMM_standard
DESeq2_poscounts
DESeq2_poscounts_zinbwave
DESeq2_TMM
corncob_wald
corncob_LRT
edgeR_poscounts_standard
edgeR_TMM_robustDisp
mgsZig_CSS
ALDEx2
limma_voom_TMM_zinbwave
limma_voom_TMM
scde
seurat_wilcoxon
MAST

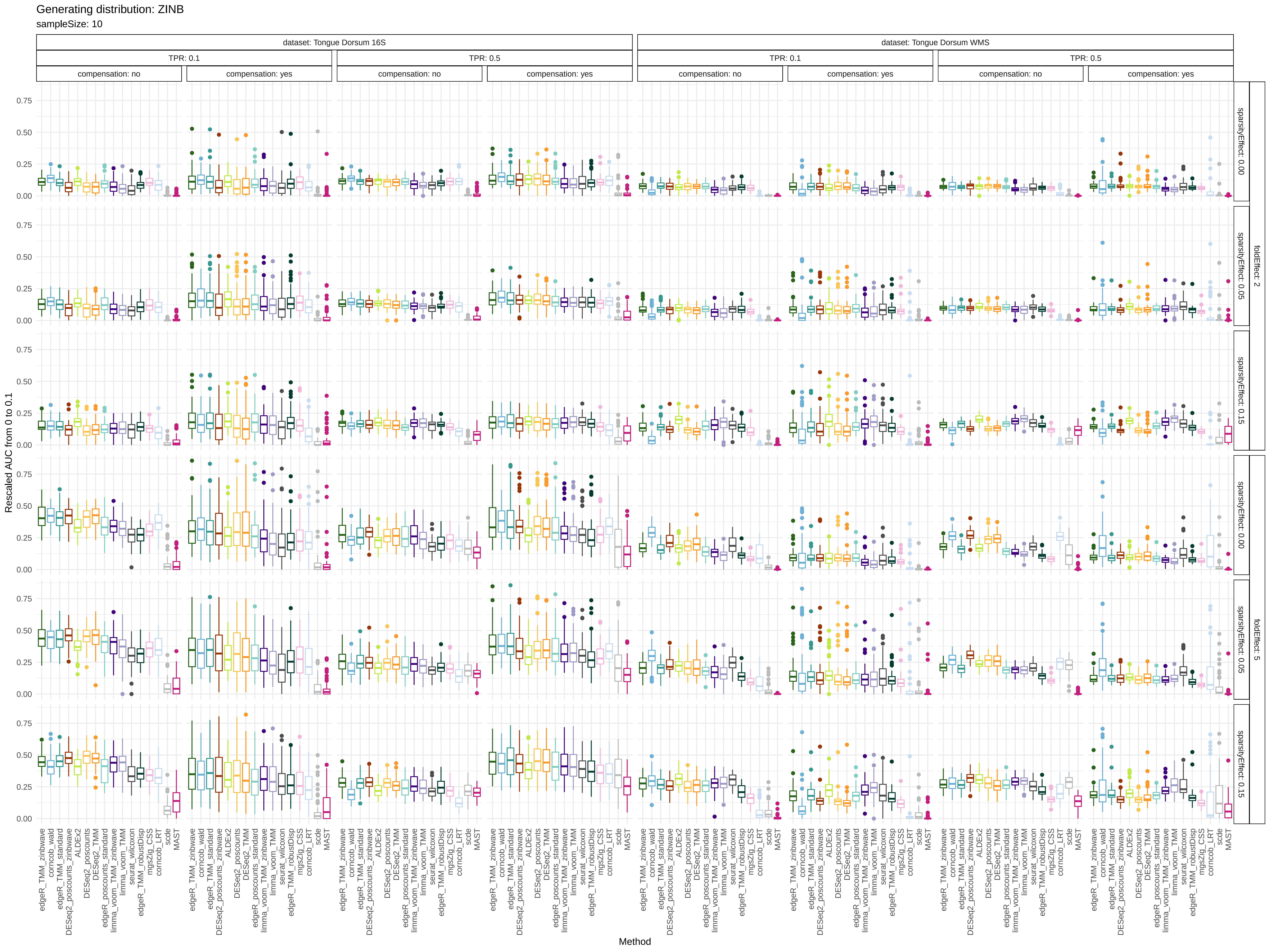

Generating distribution: ZINB
sampleSize: 10
dataset: Tongue Dorsum 16S
dataset: Tongue Dorsum WMS
TPR: 0.1
TPR: 0.5
compensation: no
compensation: yes
foldEffect: 2
foldEffect: 5
sparsityEffect: 0.00
sparsityEffect: 0.05
sparsityEffect: 0.15
Rescaled AUC from 0 to 0.1
0.00
0.25
0.50
0.75
edgeR_TMM_zinbwave
corncob_wald
edgeR_TMM_standard
DESeq2_poscounts_zinbwave
ALDEx2
DESeq2_poscounts
DESeq2_TMM
edgeR_poscounts_standard
limma_voom_TMM_zinbwave
limma_voom_TMM
seurat_wilcoxon
edgeR_TMM_robustDisp
mgsZig_CSS
corncob_LRT
scde
MAST
Method

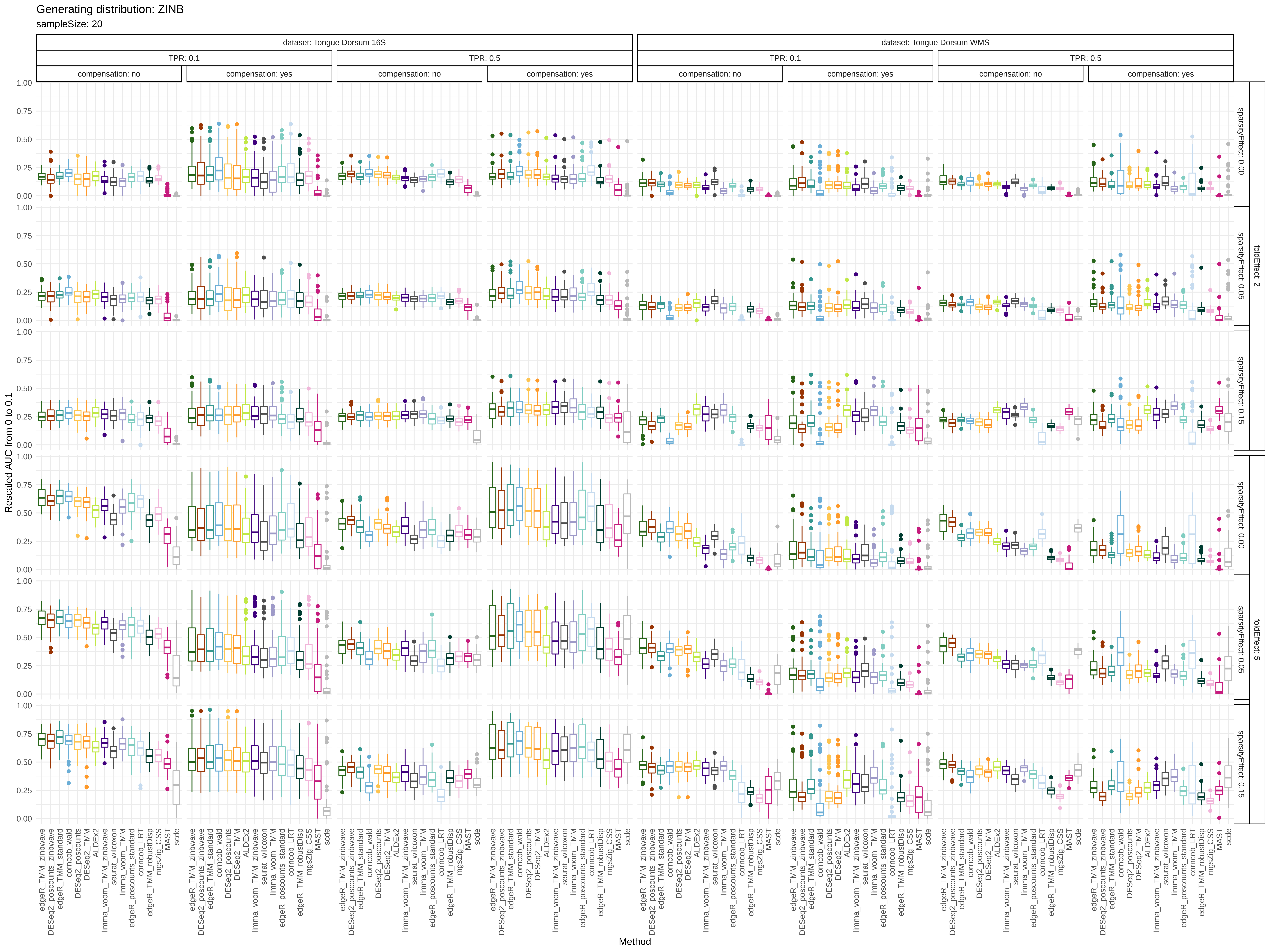
Generating distribution: ZINB
sampleSize: 20
dataset: Tongue Dorsum 16S
dataset: Tongue Dorsum WMS
TPR: 0.1
TPR: 0.5
compensation: no
compensation: yes
foldEffect: 2
foldEffect: 5
sparsityEffect: 0.00
sparsityEffect: 0.05
sparsityEffect: 0.15
Rescaled AUC from 0 to 0.1
Method
edgeR_TMM_zinbwave
DESeq2_poscounts_zinbwave
edgeR_TMM_standard
corncob_wald
DESeq2_poscounts
DESeq2_TMM
ALDEx2
limma_voom_TMM_zinbwave
seurat_wilcoxon
limma_voom_TMM
edgeR_poscounts_standard
corncob_LRT
edgeR_TMM_robustDisp
mgsZig_CSS
MAST
scde

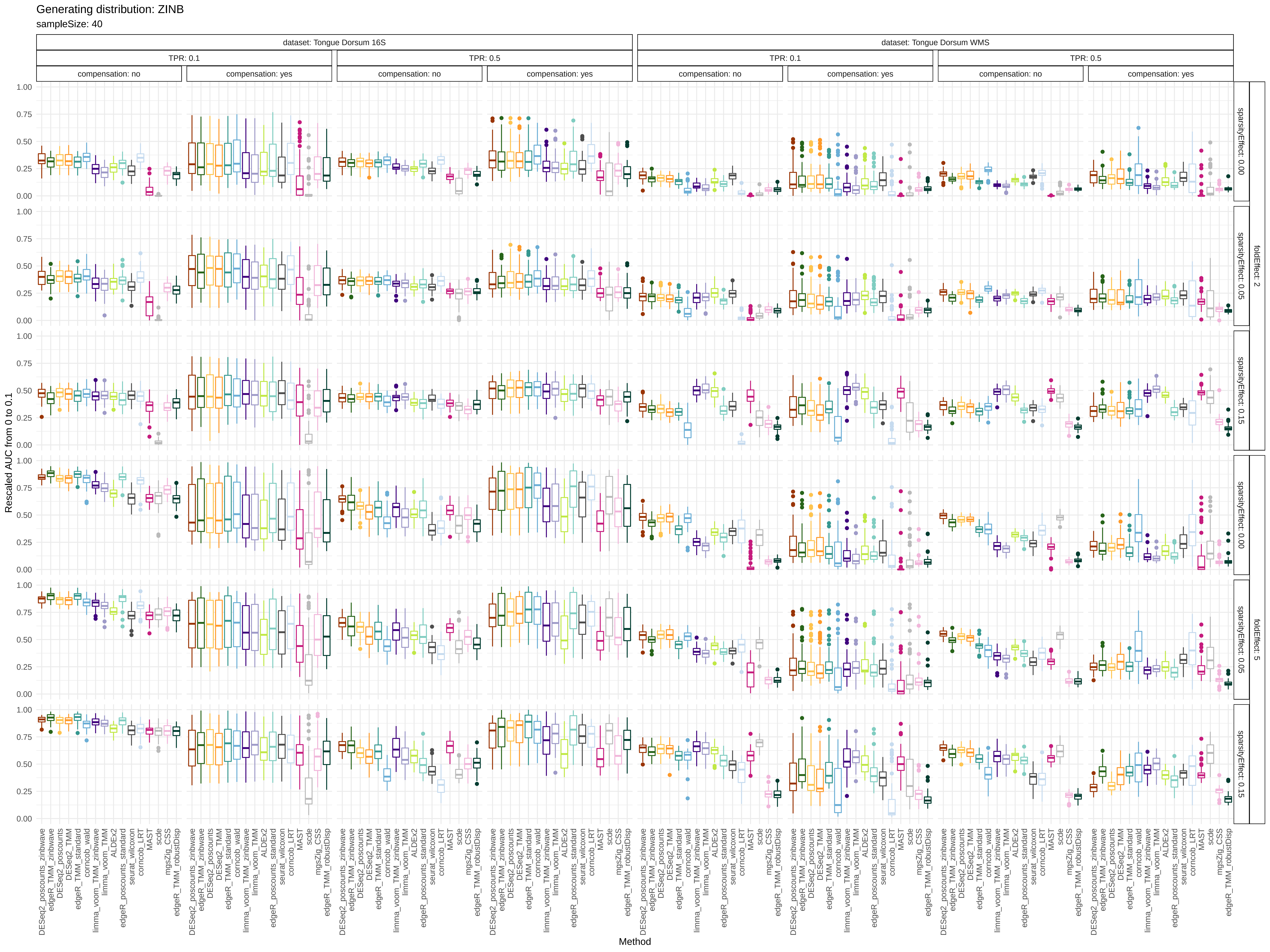

Generating distribution: ZINB
sampleSize: 40
dataset: Tongue Dorsum 16S
dataset: Tongue Dorsum WMS
TPR: 0.1
TPR: 0.5
compensation: no
compensation: yes
foldEffect: 2
foldEffect: 5
sparsityEffect: 0.00
sparsityEffect: 0.05
sparsityEffect: 0.15
Rescaled AUC from 0 to 0.1
1.00
0.75
0.50
0.25
0.00
Method
DESeq2_poscounts_zinbwave
edgeR_TMM_zinbwave
DESeq2_poscounts
DESeq2_TMM
edgeR_TMM_standard
corncob_wald
limma_voom_TMM_zinbwave
limma_voom_TMM
ALDEx2
edgeR_poscounts_standard
seurat_wilcoxon
corncob_LRT
MAST
scde
mgsZig_CSS
edgeR_TMM_robustDisp

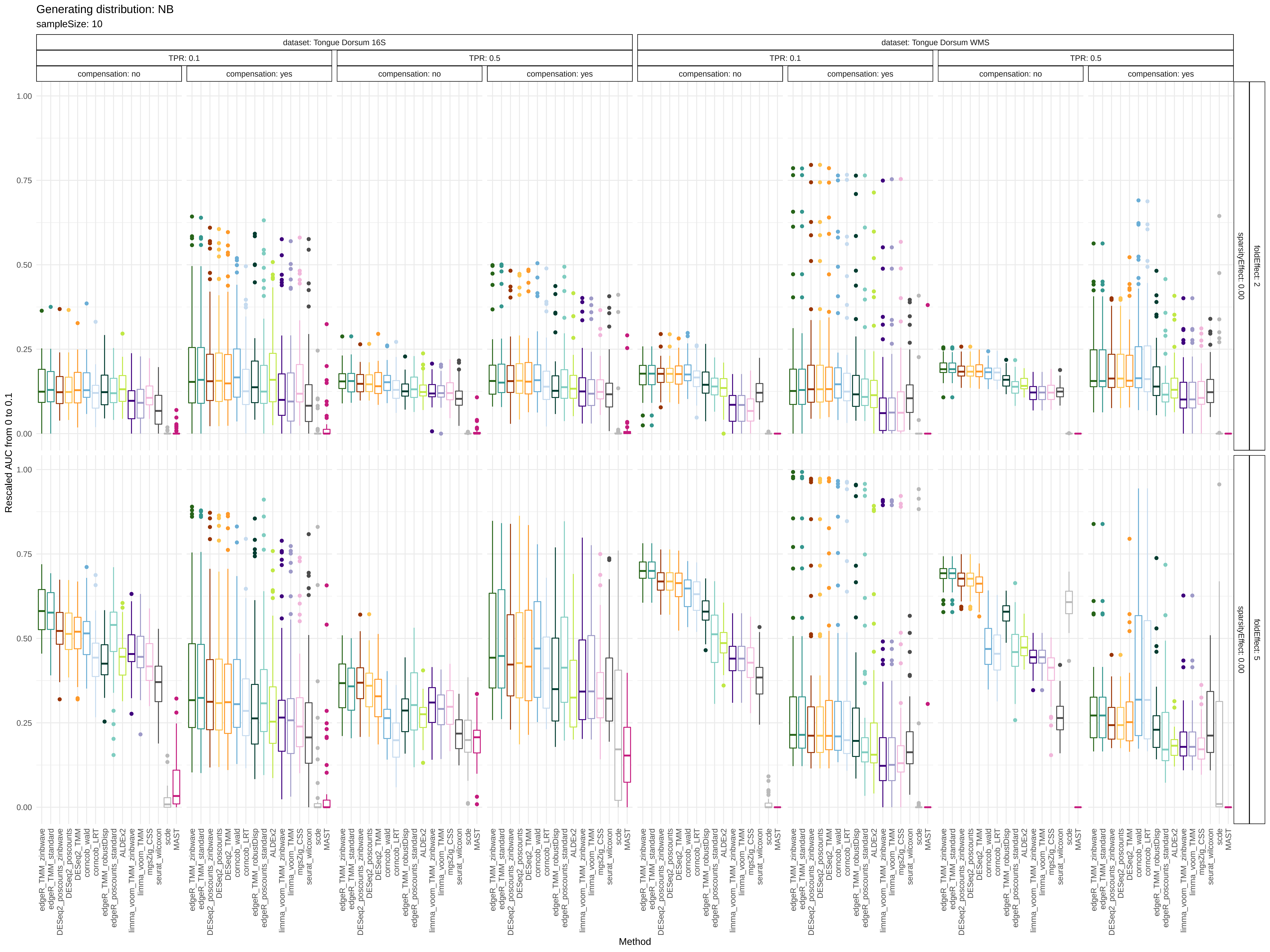

Generating distribution: NB
sampleSize: 10
dataset: Tongue Dorsum 16S
dataset: Tongue Dorsum WMS
TPR: 0.1
TPR: 0.5
compensation: no
compensation: yes
foldEffect: 2
foldEffect: 5
sparsityEffect: 0.00
Rescaled AUC from 0 to 0.1
Method
0.00
0.25
0.50
0.75
1.00
edgeR_TMM_zinbwave
edgeR_TMM_standard
DESeq2_poscounts_zinbwave
DESeq2_poscounts
DESeq2_TMM
corncob_wald
corncob_LRT
edgeR_TMM_robustDisp
edgeR_poscounts_standard
ALDEx2
limma_voom_TMM_zinbwave
limma_voom_TMM
mgsZig_CSS
seurat_wilcoxon
scde
MAST

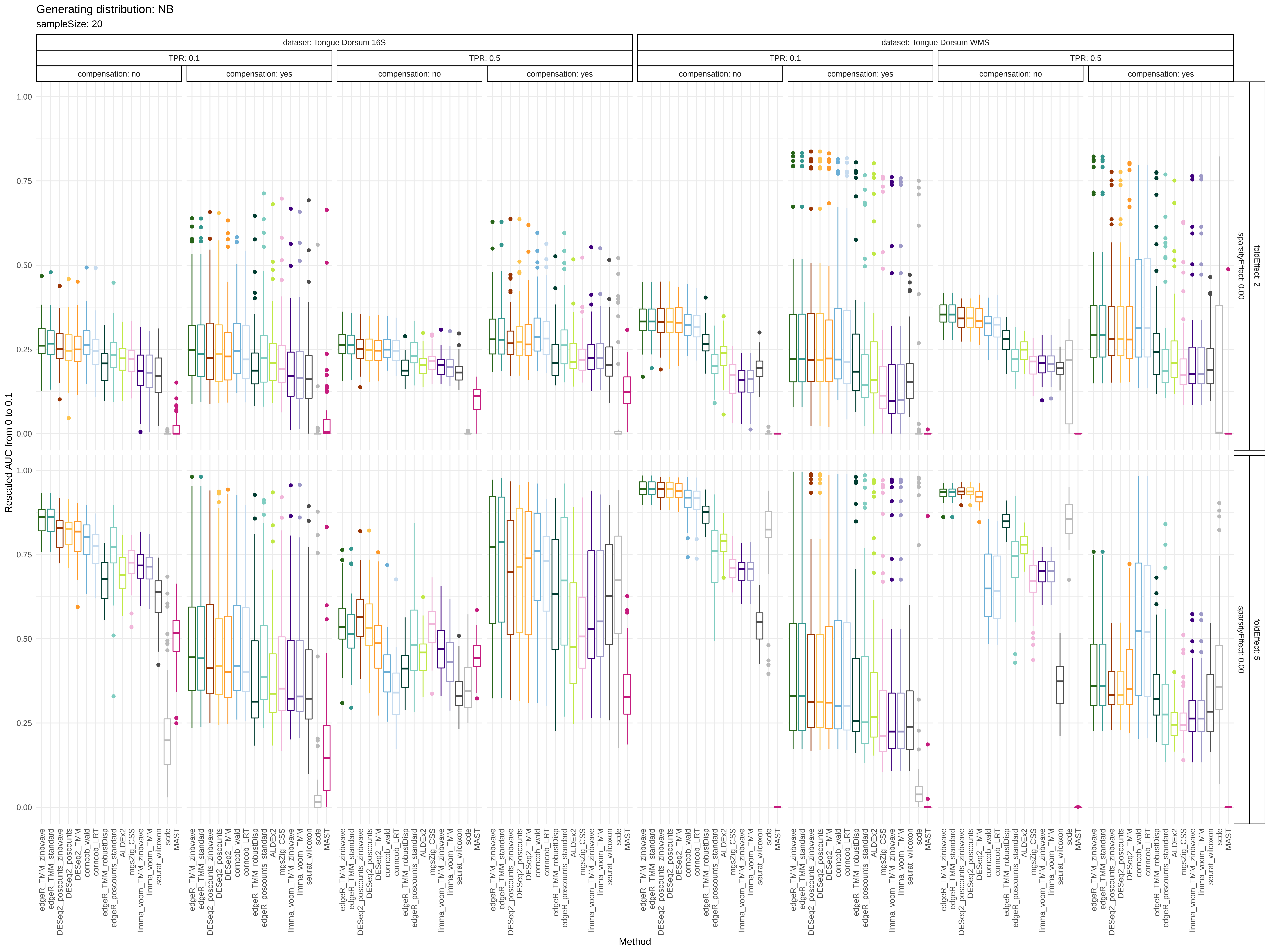
Generating distribution: NB
sampleSize: 20
dataset: Tongue Dorsum 16S
dataset: Tongue Dorsum WMS
TPR: 0.1
TPR: 0.5
compensation: no
compensation: yes
foldEffect: 2
foldEffect: 5
sparsityEffect: 0.00
Rescaled AUC from 0 to 0.1
1.00
0.75
0.50
0.25
0.00
Method
edgeR_TMM_zinbwave
edgeR_TMM_standard
DESeq2_poscounts_zinbwave
DESeq2_poscounts
DESeq2_TMM
corncob_wald
corncob_LRT
edgeR_TMM_robustDisp
edgeR_poscounts_standard
ALDEx2
mgsZig_CSS
limma_voom_TMM_zinbwave
limma_voom_TMM
seurat_wilcoxon
scde
MAST

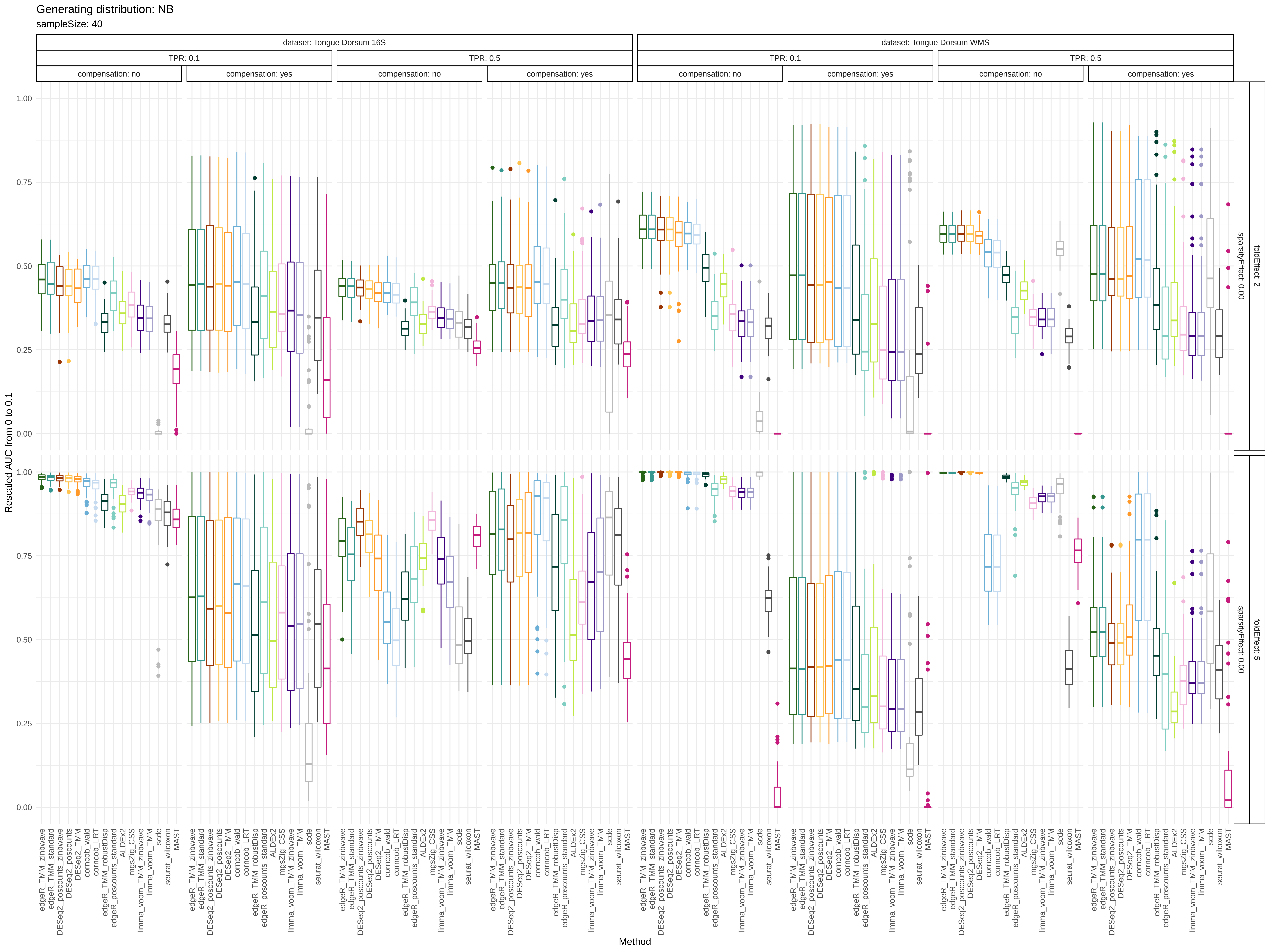

Generating distribution: NB
sampleSize: 40
dataset: Tongue Dorsum 16S
dataset: Tongue Dorsum WMS
TPR: 0.1
TPR: 0.5
compensation: no
compensation: yes
foldEffect: 2
foldEffect: 5
sparsityEffect: 0.00
Rescaled AUC from 0 to 0.1
0.00
0.25
0.50
0.75
1.00
Method
edgeR_TMM_zinbwave
edgeR_TMM_standard
DESeq2_poscounts_zinbwave
DESeq2_poscounts
DESeq2_TMM
corncob_wald
corncob_LRT
edgeR_TMM_robustDisp
edgeR_poscounts_standard
ALDEx2
mgsZig_CSS
limma_voom_TMM_zinbwave
limma_voom_TMM
scde
seurat_wilcoxon
MAST

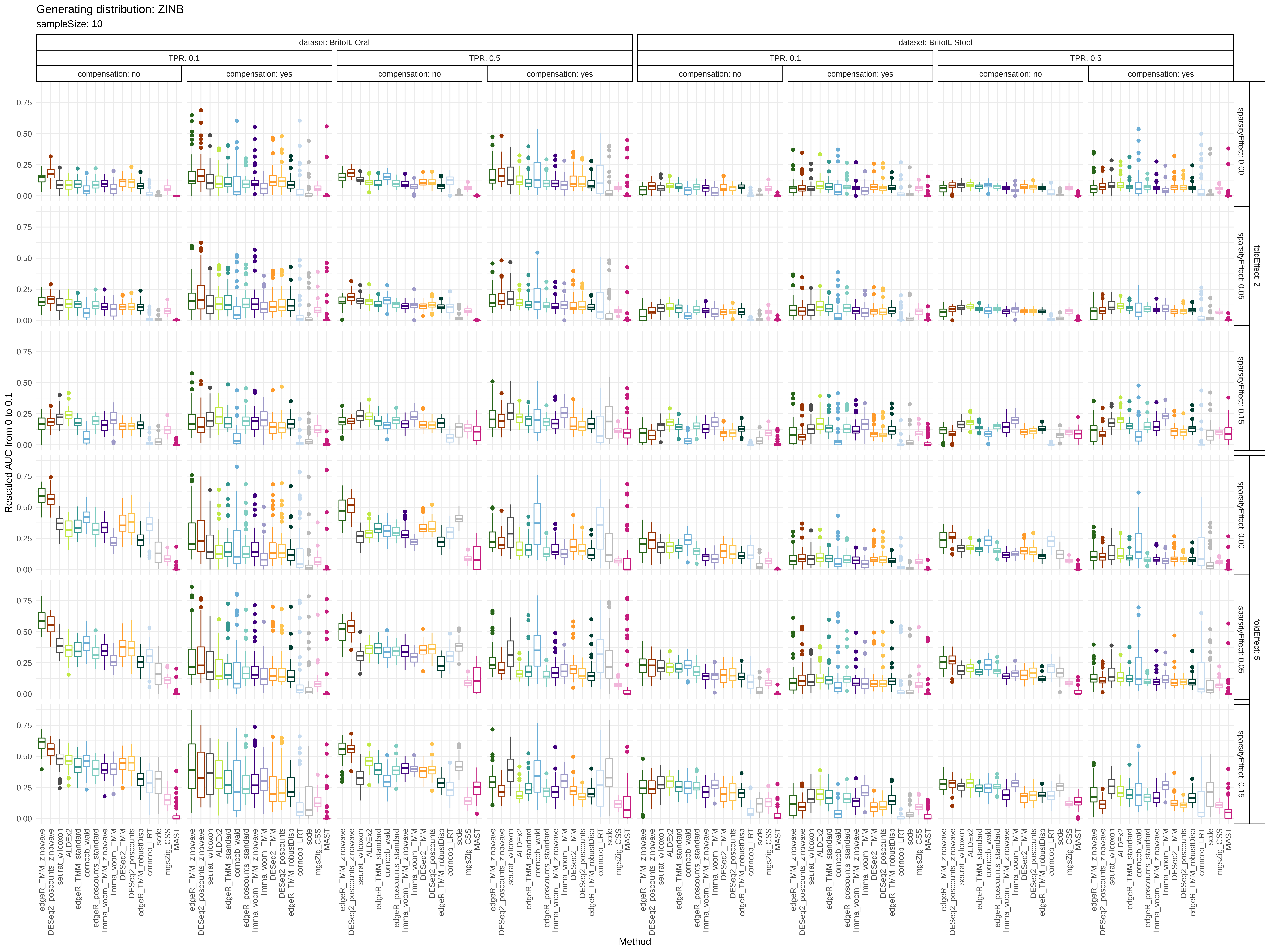

Generating distribution: ZINB
sampleSize: 10
dataset: BritoIL Oral
dataset: BritoIL Stool
TPR: 0.1
TPR: 0.5
compensation: no
compensation: yes
foldEffect: 2
foldEffect: 5
sparsityEffect: 0.00
sparsityEffect: 0.05
sparsityEffect: 0.15
Rescaled AUC from 0 to 0.1
0.00
0.25
0.50
0.75
Method
edgeR_TMM_zinbwave
DESeq2_poscounts_zinbwave
seurat_wilcoxon
ALDEx2
edgeR_TMM_standard
corncob_wald
edgeR_poscounts_standard
limma_voom_TMM_zinbwave
limma_voom_TMM
DESeq2_TMM
DESeq2_poscounts
edgeR_TMM_robustDisp
corncob_LRT
scde
mgsZig_CSS
MAST

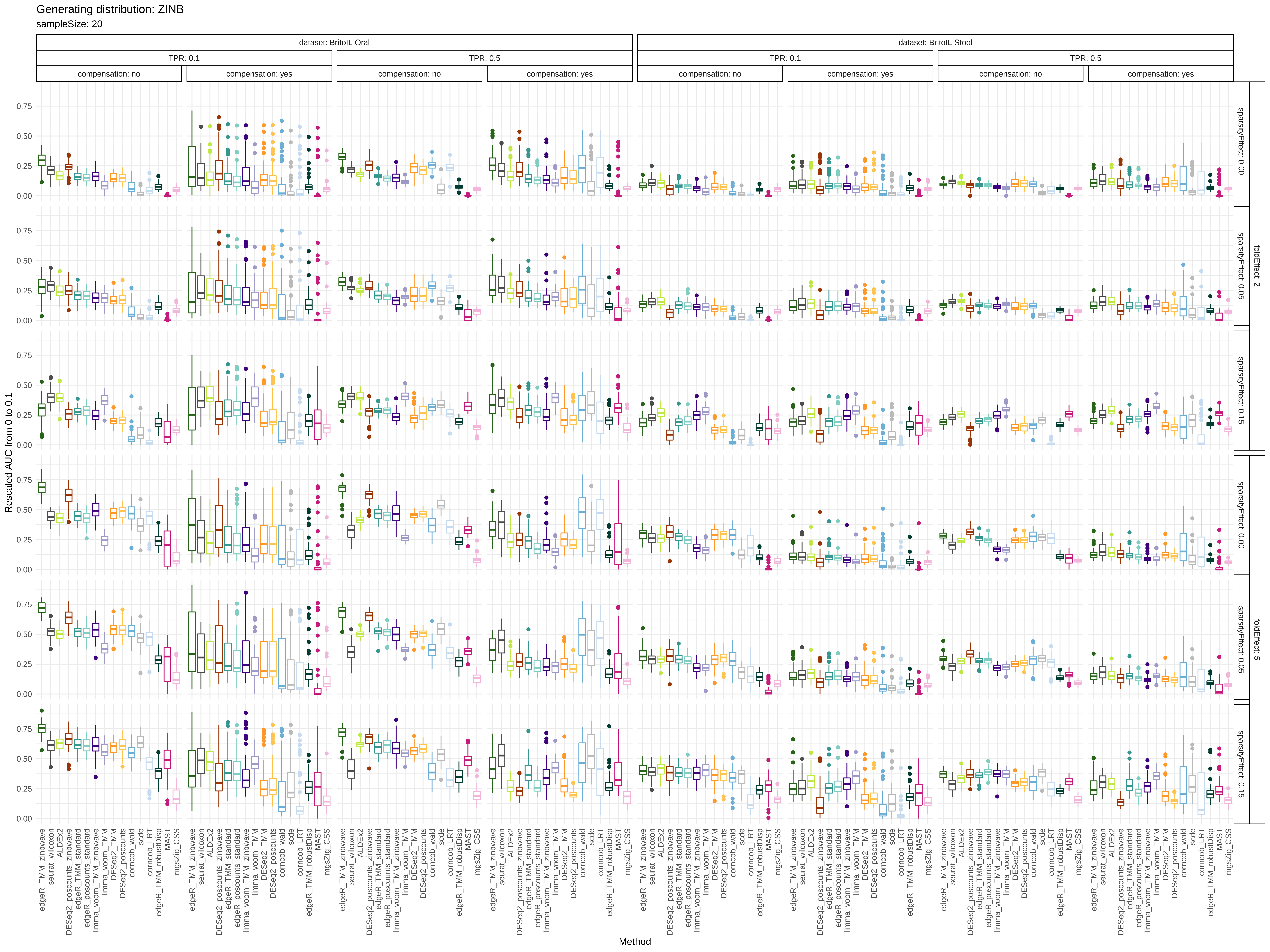

Generating distribution: ZINB
sampleSize: 20
dataset: BritoIL Oral
dataset: BritoIL Stool
TPR: 0.1
TPR: 0.5
compensation: no
compensation: yes
foldEffect: 2
foldEffect: 5
sparsityEffect: 0.00
sparsityEffect: 0.05
sparsityEffect: 0.15
Rescaled AUC from 0 to 0.1
0.00
0.25
0.50
0.75
edgeR_TMM_zinbwave
seurat_wilcoxon
ALDEx2
DESeq2_poscounts_zinbwave
edgeR_TMM_standard
edgeR_poscounts_standard
limma_voom_TMM_zinbwave
limma_voom_TMM
DESeq2_TMM
DESeq2_poscounts
corncob_wald
scde
corncob_LRT
edgeR_TMM_robustDisp
MAST
mgsZig_CSS
Method

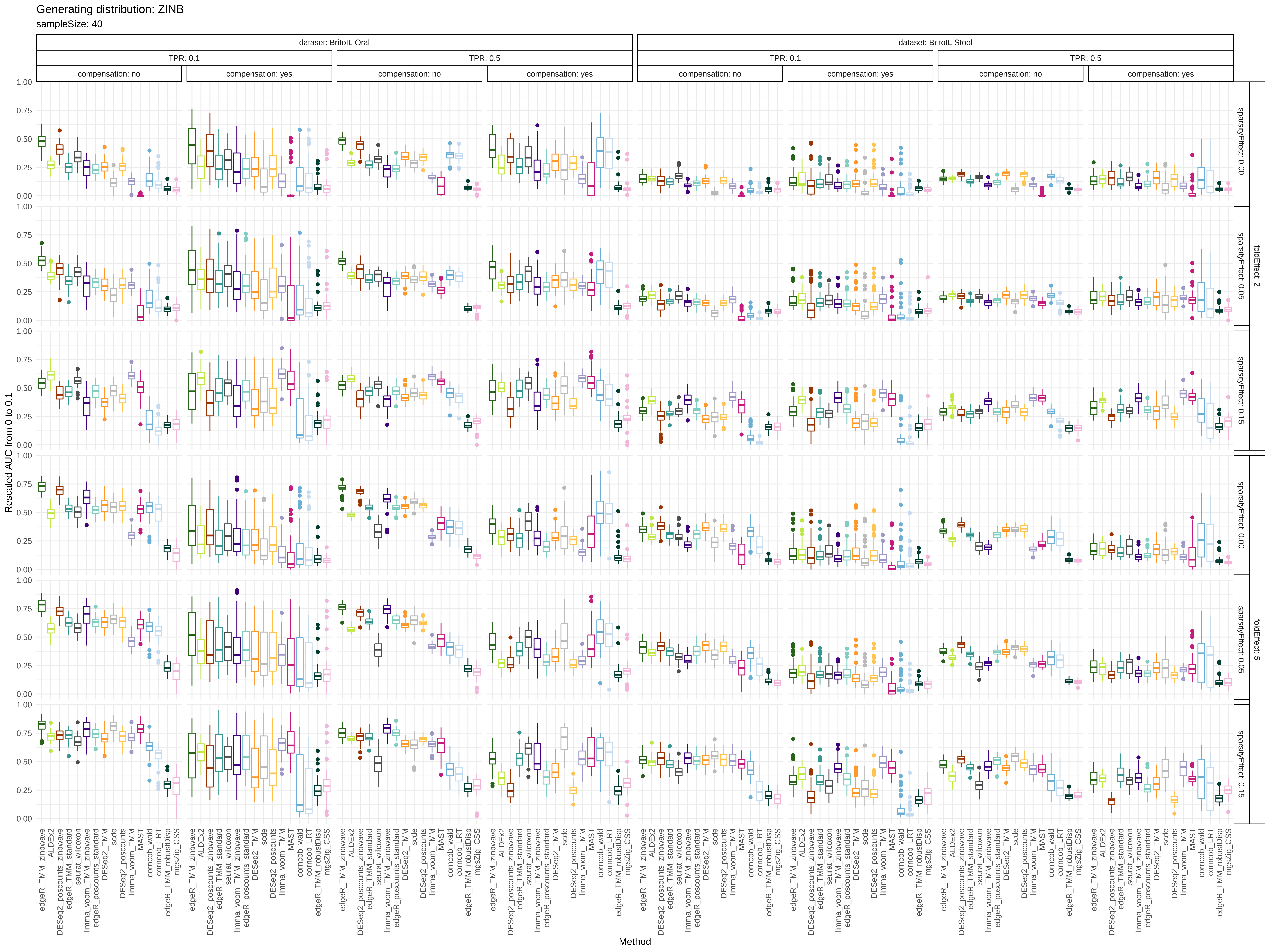
Generating distribution: ZINB
sampleSize: 40
dataset: BritoIL Oral
dataset: BritoIL Stool
TPR: 0.1
TPR: 0.5
compensation: no
compensation: yes
sparsityEffect: 0.00
sparsityEffect: 0.05
sparsityEffect: 0.15
foldEffect: 2
foldEffect: 5
Rescaled AUC from 0 to 0.1
Method
edgeR_TMM_zinbwave
ALDEx2
DESeq2_poscounts_zinbwave
edgeR_TMM_standard
seurat_wilcoxon
limma_voom_TMM_zinbwave
edgeR_poscounts_standard
DESeq2_TMM
scde
DESeq2_poscounts
limma_voom_TMM
MAST
corncob_wald
corncob_LRT
edgeR_TMM_robustDisp
mgsZig_CSS

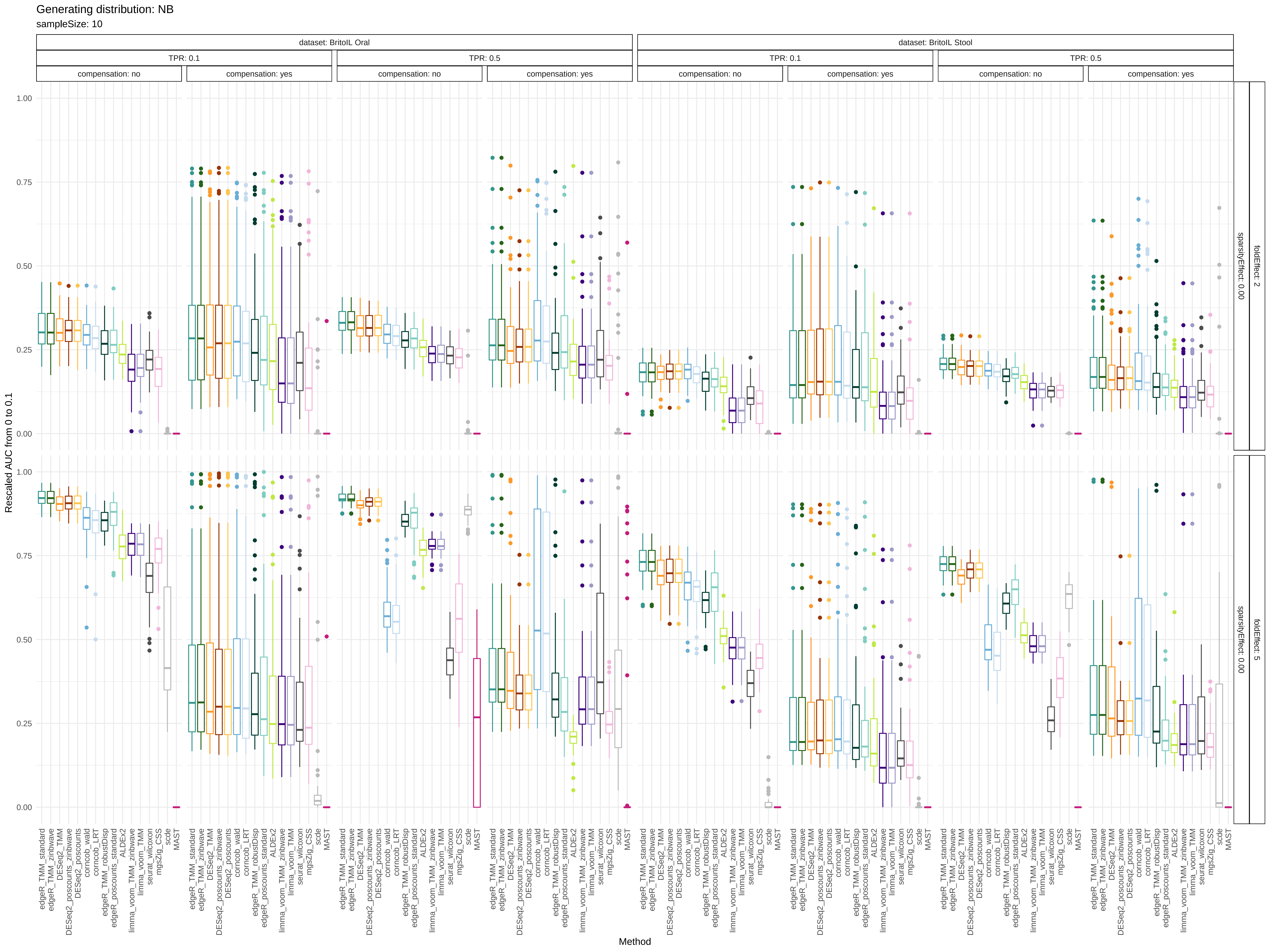

Generating distribution: NB
sampleSize: 10
dataset: BritoIL Oral
dataset: BritoIL Stool
TPR: 0.1
TPR: 0.5
compensation: no
compensation: yes
foldEffect: 2
foldEffect: 5
sparsityEffect: 0.00
Rescaled AUC from 0 to 0.1
0.00
0.25
0.50
0.75
1.00
Method
edgeR_TMM_standard
edgeR_TMM_zinbwave
DESeq2_TMM
DESeq2_poscounts_zinbwave
DESeq2_poscounts
corncob_wald
corncob_LRT
edgeR_TMM_robustDisp
edgeR_poscounts_standard
ALDEx2
limma_voom_TMM_zinbwave
limma_voom_TMM
seurat_wilcoxon
mgsZig_CSS
scde
MAST

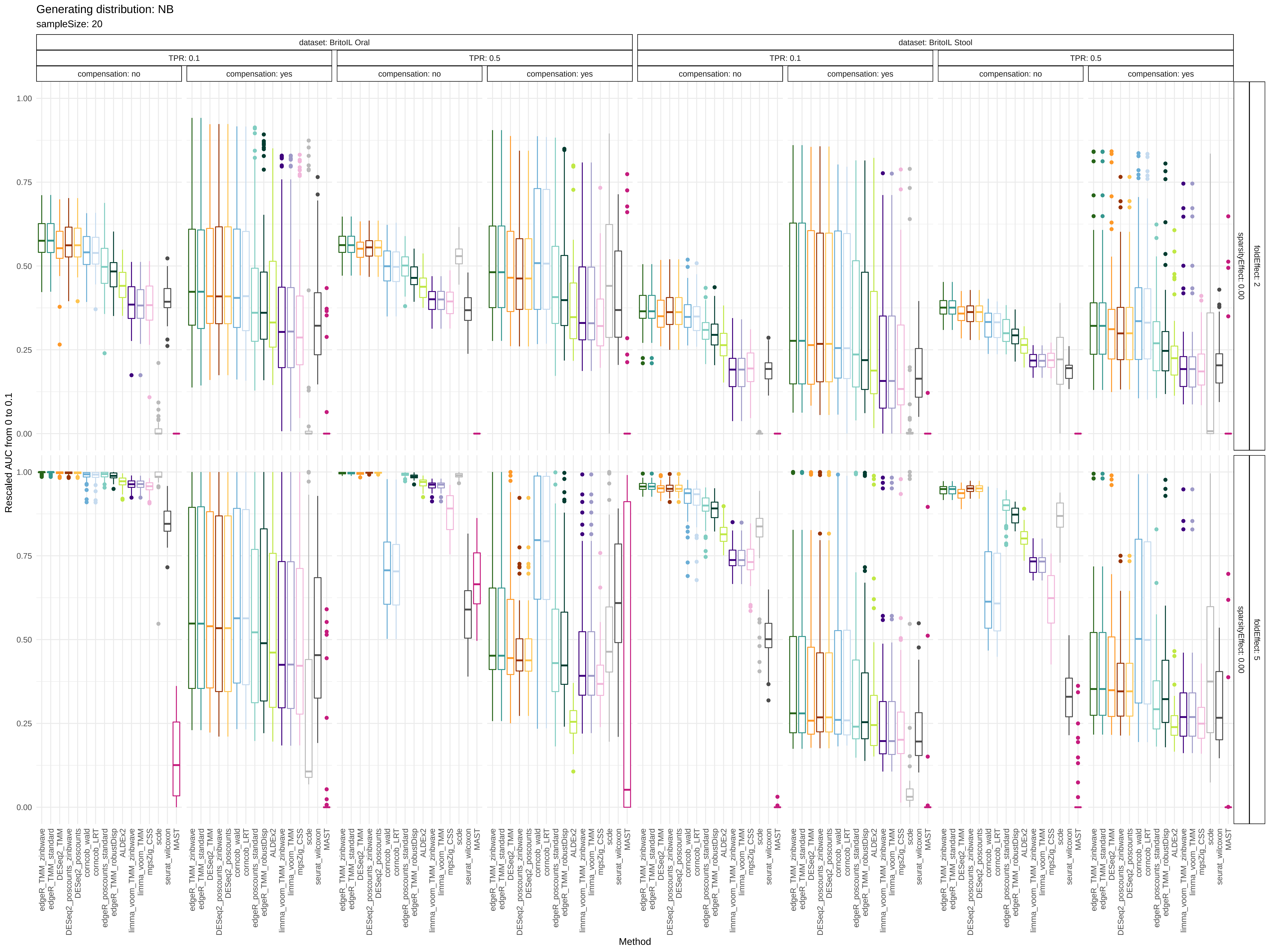

Generating distribution: NB
sampleSize: 20
dataset: BritoIL Oral
dataset: BritoIL Stool
TPR: 0.1
TPR: 0.5
compensation: no
compensation: yes
foldEffect: 2
foldEffect: 5
sparsityEffect: 0.00
Rescaled AUC from 0 to 0.1
0.00
0.25
0.50
0.75
1.00
edgeR_TMM_zinbwave
edgeR_TMM_standard
DESeq2_TMM
DESeq2_poscounts_zinbwave
DESeq2_poscounts
corncob_wald
corncob_LRT
edgeR_poscounts_standard
edgeR_TMM_robustDisp
ALDEx2
limma_voom_TMM_zinbwave
limma_voom_TMM
mgsZig_CSS
scde
seurat_wilcoxon
MAST
Method

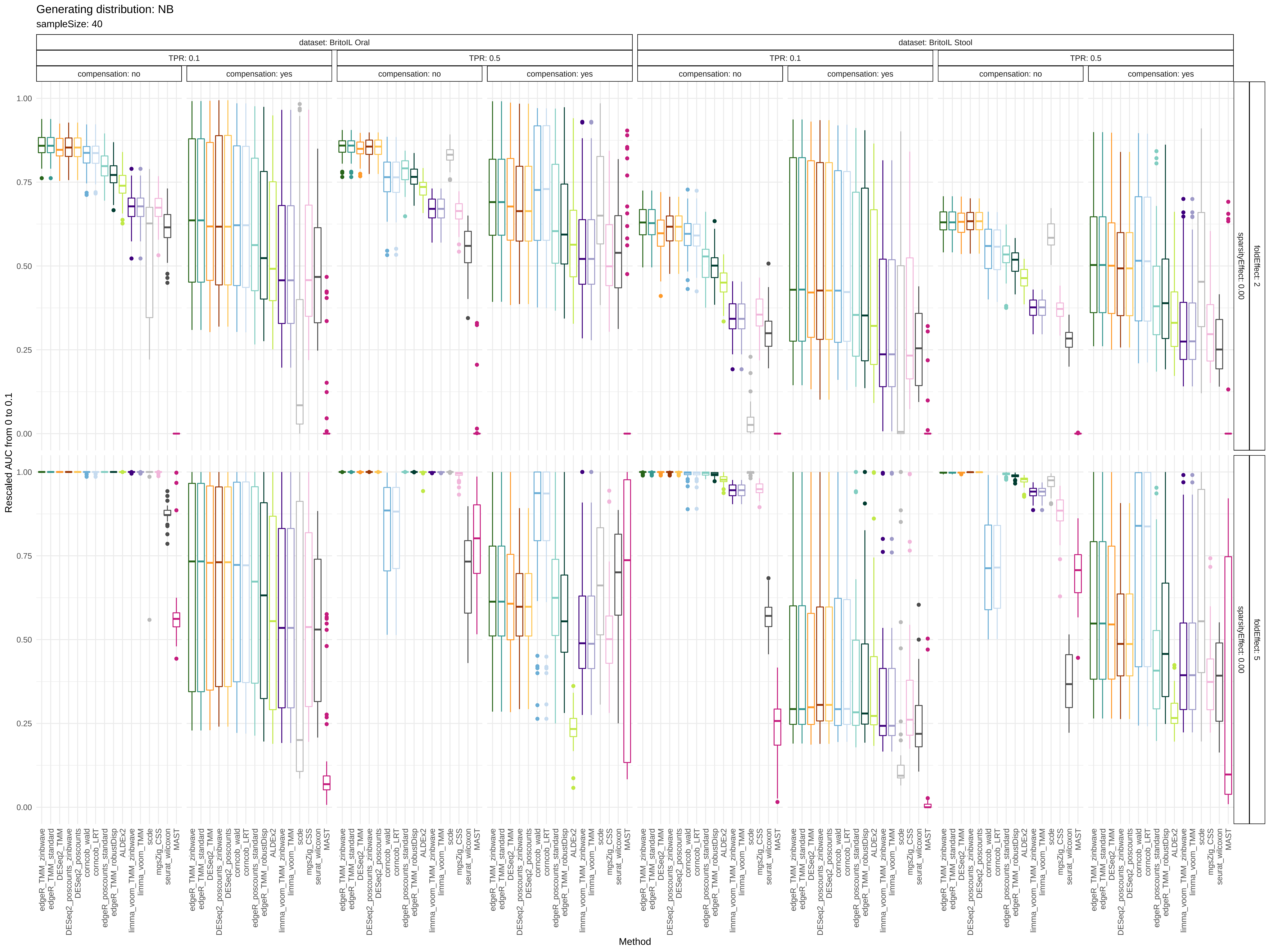

Generating distribution: NB
sampleSize: 40
dataset: BritoIL Oral
dataset: BritoIL Stool
TPR: 0.1
TPR: 0.5
compensation: no
compensation: yes
foldEffect: 2
foldEffect: 5
sparsityEffect: 0.00
Rescaled AUC from 0 to 0.1
0.00
0.25
0.50
0.75
1.00
Method
edgeR_TMM_zinbwave
edgeR_TMM_standard
DESeq2_TMM
DESeq2_poscounts_zinbwave
DESeq2_poscounts
corncob_wald
corncob_LRT
edgeR_poscounts_standard
edgeR_TMM_robustDisp
ALDEx2
limma_voom_TMM_zinbwave
limma_voom_TMM
scde
mgsZig_CSS
seurat_wilcoxon
MAST
